# A Novel Liver Cancer-Selective Histone Deacetylase Inhibitor Is Effective Against Hepatocellular Carcinoma and Induces Durable Responses with Immunotherapy

**DOI:** 10.1101/2024.03.27.587062

**Authors:** Bocheng Wu, Subhasish Tapadar, Zhiping Ruan, Carrie Q. Sun, Rebecca S. Arnold, Alexis Johnston, Jeremiah O. Olugbami, Uche Arunsi, David A. Gaul, John A. Petros, Tatsuya Kobayashi, Dan G. Duda, Adegboyega K. Oyelere

## Abstract

Hepatocellular cancer (HCC) progression is facilitated by gene-silencing chromatin histone hypoacetylation due to histone deacetylases (HDACs) activation. However, inhibiting HDACs — an effective treatment for lymphomas — has shown limited success in solid tumors. We report the discovery of a class of HDAC inhibitors (HDACi) that demonstrates exquisite selective cytotoxicity against human HCC cells. The lead compound **STR-V-53** (**3**) showed a favorable safety profile in mice and robustly suppressed tumor growth in orthotopic xenograft models of HCC. When combined with the anti-HCC drug sorafenib, **STR-V-53** showed greater in vivo efficacy. Moreover, **STR-V-53** combined with anti-PD1 therapy increased the CD8^+^ to regulatory T-cell (Treg) ratio and survival in an orthotopic HCC model in immunocompetent mice. This combination therapy resulted in durable responses in 40% of the mice. Transcriptomic analysis revealed that **STR-V-53** primed HCC cells to immunotherapy through HDAC inhibition, impaired glucose-regulated transcription, impaired DNA synthesis, upregulated apoptosis, and stimulated the immune response pathway. Collectively, our data demonstrate that the novel HDACi **STR-V-53** is an effective anti-HCC agent that can induce profound responses when combined with standard immunotherapy.

**Graphical Abstract:** 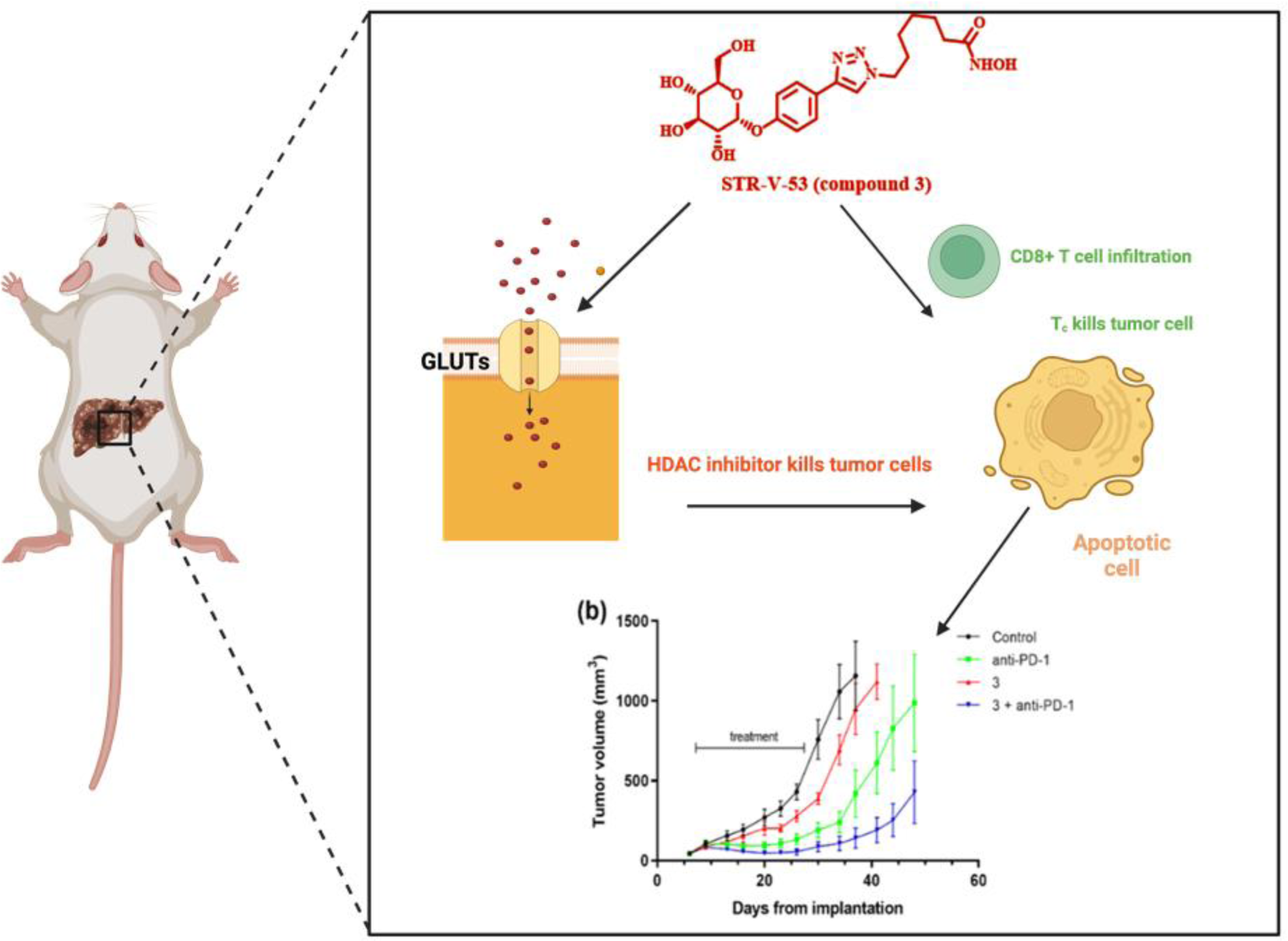

## Introduction

With more than 800,000 diagnosed cases and over 700,000 fatalities annually, liver cancer is a leading cause of global cancer deaths^1^. Hepatocellular carcinoma (HCC) is the most common liver cancer^2,3^. HCC is responsible for over 80% of all liver cancer cases, and over 90% of the time, it occurs in patients with liver damage. The prognosis of HCC is dismal, with an estimated 10-12% 5-year survival^4^. Surgical resection is a curative treatment option for HCC. However, surgery is limited to patients with the early stage of the disease and good liver function. Only a small proportion of HCC patients are eligible for surgery, and the post-surgery relapse rate is high (50-80%)^5,6^. Liver transplantation is another treatment option for localized HCC, but the shortage of organs significantly limits this option^7,8^. Systemic therapy became a main treatment option for unresectable/advanced HCC after the FDA approved sorafenib, an inhibitor of multiple kinases, including RAF, VEGFR, PDGFR, and other receptor tyrosine kinases^9^. The subsequent approval of other multikinase inhibitors, such as lenvatinib, regorafenib, and cabozantinib, has added more pharmacological options for HCC treatment^10^. However, these drugs generally extend survival in advanced HCC patients by only 1-3 months^9^. More recently, an immunotherapy strategy, involving the blockade of immune checkpoints (PD-1 alone or with CTLA-4 or VEGF antibodies), has emerged as an efficacious approach. However, most patients (>70%) do not respond to these treatments^11^. Therefore, there is a significant need for more efficacious treatment modalities for HCC that target malignant cells.

Histone deacetylase (HDAC) inhibition is a clinically validated epigenetic-based strategy for cancer treatment. HDACs, through their lysine deacetylase activities, play important roles in epigenetically moderating gene expression at the chromatin level. Dysregulation of HDACs expression has been linked to the proliferation, survival, and invasiveness of HCC^12^.

Specifically, the overexpression of class I HDACs (HDACs 1 and 2) and class IIa HDACs (HDACs 4, 7, and 9) in HCC cells and patient samples strongly correlates with reduced patient survival. To date, five HDAC inhibitors (HDACi) – vorinostat (SAHA), belinostat, chidamide, romidepsin, and panobinostat (Fig. S1a) – have been approved to treat hematological malignancies^13^. The potential of belinostat as an anti-HCC agent has been studied in a clinical trial (NCT00321594)^14^. The trial results revealed that patients on belinostat treatment had median progression-free survival (PFS) and overall survival (OS) of 2.64 and 6.60 months, respectively^14^. While this result supports the potential of HDACi for HCC treatment, enhancing their efficacy remains a key challenge.

In previous studies, we have found that bio-inspired alterations to the HDACi surface recognition cap group could furnish HDACi with cell and tissue selectivity^15–17^. More relevant to HCC, we have found that selective-liver tissue accumulation was a viable approach to improving the anti-HCC efficacy of HDACi. We discovered that integrating macrolide azithromycin into the surface recognition group of sub-class I HDAC isoform-selective HDACi resulted in macrolide-based HDACi (Fig. S1b), which preferentially accumulated in liver tissue and robustly suppressed HCC tumor growth in an orthotopic model^18^.

In this study, we explored targeting the Warburg effect to deliver HDACi to cancer cells selectively. The Warburg effect describes an altered metabolic state in which cancer cells rely on glycolysis for energy sources even in the presence of oxygen^19^. To sustain this altered metabolism and proliferation, cancer cells upregulate the expression levels of several glucose transporters (GLUT) to facilitate enhanced uptake. GLUT-1, GLUT-2, GLUT-3, and GLUT-4 are overexpressed on several cancer cell types^20,21^. HCCs are known to overexpress GLUT-2, which effectively promotes the uptake of glucose and mannose to support tumor growth, such that GLUT-2 is recognized as a novel prognostic factor of HCC^22^. GLUT-2 is a major sugar transport facilitator in the hepatocytes, having higher capacity and lower binding affinity to glucose^21^. In addition, GLUT-2 could promote the cell uptake of fructose and several glycosylated small molecules. This unique property makes GLUT-2 the preferred sugar transporter relative to the other GLUTs^21,23^. Based on this understanding, we designed and synthesized four classes of novel glycosylated HDACi having glycoside moieties (D-glucose, D-mannose, and desosamine sugar) integrated into the prototypical HDACi surface recognition cap group (Fig. 1). We evaluated the HDAC inhibitory activities of the glycosylated HDACi against representative class I and class II HDACs and screened them using a representative panel of cancer and normal cell lines.

**Figure 1.**
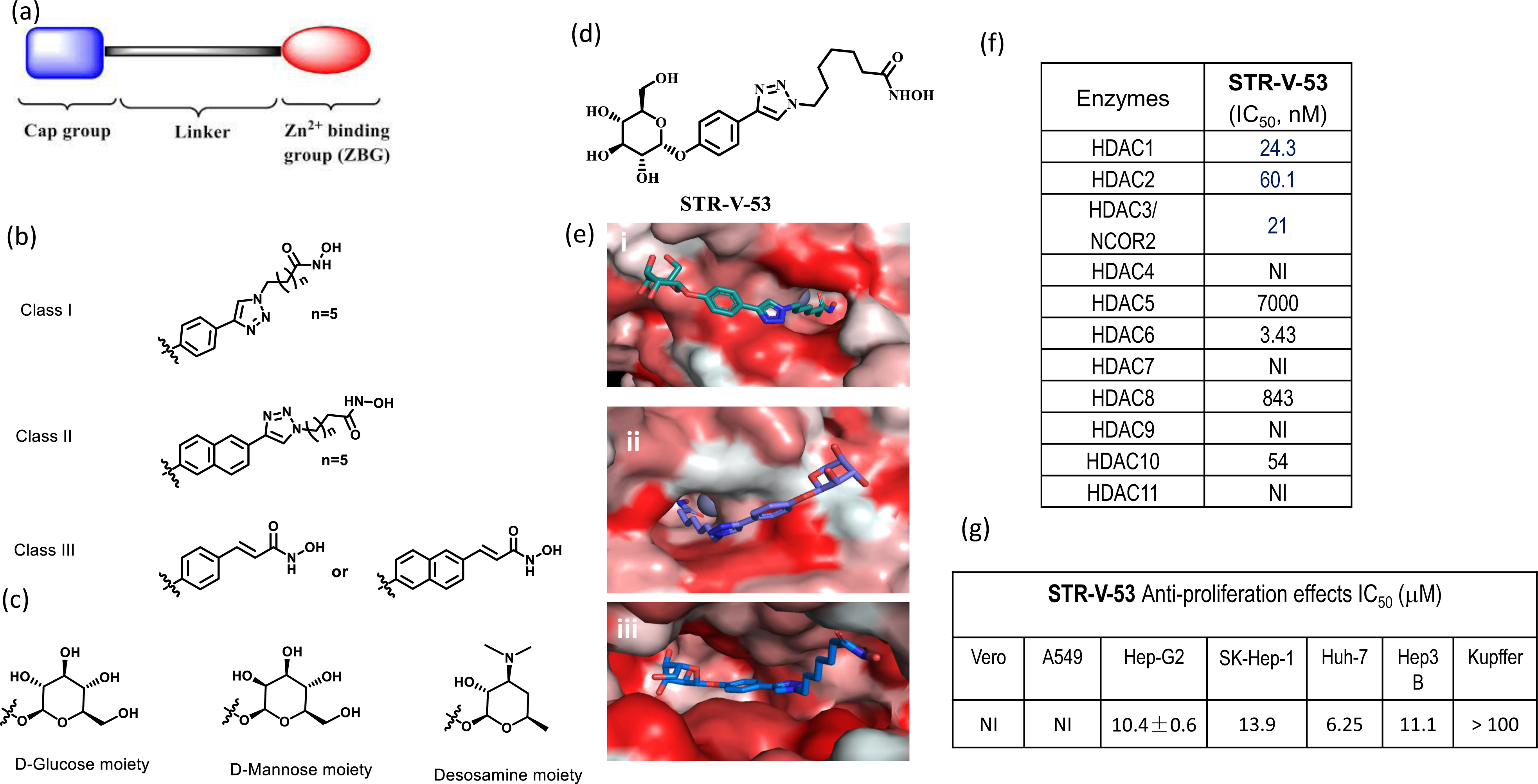
Design of glycosylated HDACi. (a) HDACi three pharmacophoric model. (b) The aglycone moieties of the three subclasses of the designed HDACi. (c) The glycoside moieties of the designed HDACi. (d) Structure of lead compound **STR-V-53**. (e) Docked poses of **STR-V-53** at the active sites of HDAC2 (PDB:4LXZ) (i), and HDAC6 (5G0G) (ii) and GLUT-1 (PDB: 4PYP) (iii). The compounds’ linker regions and the hydroxamate moieties maintained optimal interactions with hydrophobic residues lining the pockets to the base of the active sites and efficient zinc chelation, respectively; while the glycoside moieties are oriented toward the solvent-exposed hydrophilic regions at the HDACs outer rims (i and ii). Within GLUT-1, **STR-V-53** is accommodated within the nonyl beta-D-glucopyranoside binding pocket (iii). (f) HDAC isoform inhibition activities of **STR-V-53** (IC_50_ in nM). (g) Anti-proliferation effects of **STR-V-53** (IC_50_ in μM). NI= no inhibition up to 100µM.

We found that these compounds demonstrated potent HDAC inhibition activities, and a cohort of (**STR-V-53 (3)**, **STR-I-195**, and **STR-V-114**) are selectively cytotoxic to an HCC cell line. Also, we found that a representative HCC cell line (Hep-G2) largely uptakes the glycosylated HDACi through the GLUT-2 transporter. In addition, these HDACi caused HCC cell-line-dependent apoptosis through caspase 3 cleavage and p21 upregulation. We also found that a selected candidate compound, **STR-V-53** efficiently suppressed HCC tumor growth over a 21-day treatment period with no significant toxicity. Subsequently, we screened **STR-V-53** in the NCI-60 panel. Although the NCI-60 panel lacks liver cancer cell lines, we performed this experiment to obtain further information about the cancer cell-type selectivity of **STR-V-53**. In a one-dose experiment, we observed that **STR-V-53** has a negligible effect on proliferation, showing a mean cell growth of 99.6%. Our cell activity data on **STR-V-53** and the lack of growth inhibition effect of **STR-V-53** in the NCI-60 panel strongly support our observations of the HCC cell-selectivity attributes of **STR-V-53**. Finally, when combined with anti-PD1 therapy, **STR-V-53** reprogrammed the T cell responses and prolonged median overall survival in an orthotopic HCC model in immunocompetent mice. More importantly, this combination resulted in durable responses in 40% of the mice. Collectively, our data demonstrate that the novel HDACi **STR-V-53** is an effective anti-HCC agent that induces profound responses in combination with standard immunotherapy. **STR-V-53** and other glycosylated HDACi disclosed herein have high translational potential as new agents for HCC treatment.

## Results

### Design of glycosylated HDACi

Standard HDACi are based on three pharmacophoric models – surface recognition group, linker moiety, and zinc binding group (ZBG) (Fig. 1a). In designing the disclosed compounds, we individually integrated three different glycosides – D-glucose, D-mannose, or desosamine – into the HDACi surface recognition groups derived from phenyl and naphthyl moieties while we adopted the linker moieties that have afforded optimum HDAC inhibition effect based on our previous studies and those by others^15,16^. As a ZBG, we used hydroxamate, a moiety that affords pan-selective inhibition. This design furnished three classes of glycosylated HDACi (Figs. 1b-c). The synthesis of the target glycosylated HDACi was accomplished as described in Schemes 1-3 (supporting information).

### Molecular Docking

To gain insight into the prospect of productive interaction between these glucosylated HDACi and HDACs and GLUT transporter, we used the *in silico* molecular docking analysis (Autodock Vina)^26^ to interrogate the binding orientations and the docking scores of selected compounds – **STR-V-53** (Fig. 1d) and **STR-V-165** (Supp. Info Scheme 3) – against HDAC2 (PDB: 4LXZ), HDAC6 (5G0G) and, GLUT-1 (4PYP) which shared 55% sequence similarity of GLUT-2. We used GLUT-1 for this analysis due to the absence of GLUT-2 structure in the PDB.

Molecular docking analyses were performed as we described before^16–18^. We observed that both compounds were optimally accommodated within the hydrophobic residues lining the pockets to the base of the active sites and efficiently chelate the Zn^2+^ ion at the active sites of HDAC2 and HDAC6. The glycoside moieties of both compounds are oriented toward the solvent-exposed hydrophilic regions at the outer rims of either enzyme (Figs. 1ei, 1eii and S2a-b). Collectively, these interactions should be crucial to effective HDAC inhibition. The docking scores of the binding poses of **STR-V-53**, which have the most plausible interactions within HDACs 2 and 6 active sites, are -8.2 kcal/mol and -8.4 kcal/mol, respectively, while **STR-V-165** has docking scores of -8.6 kcal/mol and -8.2 kcal/mol for HDACs 2 and 6 respectively.

Against GLUT-1, we found that the glucose moiety of **STR-V-53** optimally interacts within the region where the sugar analog, n-nonyl beta-D-glucopyranoside, binds (Figs. 1eiii and S2c). More specifically, the glucose moiety of **STR-V-53** forms H-bonds with Asn-288, Gln-283, Glu-380, Asn-415 (Fig. S2c). The docked pose adopted by **STR-V-53** has -12.2 kcal/mol binding affinity, a much stronger binding affinity relative to the unmodified glucose (-6.4 kcal/mol) and nonyl beta-D-glucopyranoside (–6.9 kcal/mol). The desosamine sugar, the glycoside moiety of **STR-V-165** has fewer hydroxyl groups (Fig. S2d), which attenuated the GLUT-1 binding affinity of **STR-V-165** relative to that of **STR-V-53**. Nevertheless, the **STR-V-165** GLUT-1 binding affinity of –10.1 kcal/mol is higher than those of unmodified glucose and n-nonyl beta-D-glucopyranoside.

### HDAC inhibition and Anti-proliferation activities

We evaluated all compounds for their effects on the deacetylase activities of class I HDACs 1, 2, and 8; and HDAC6, a representative class IIb HDAC (BPS Bioscience Inc., San Diego, CA, USA) while **STR-V-53** was screened against all eleven Zn^2+^-dependent HDAC isoforms. As expected for HDACi with long methylene linkers, the class I and II compounds showed single-digit to mid-nanomolar IC_50_ inhibition of HDACs 1, 2, and 6, with a strong preference for HDAC6 as predicted by our *in silico* molecular analyses, while less inhibitory of HDAC8. The cinnamate-derived class III compounds (**STR-V-105** and **STR-V-115**) have weak HDAC inhibition activities, although they are also selective for HDAC6 (Table S1). The HDAC isoforms selectivity profile of **STR-V-53** differs slightly from that of pan-HDACi SAHA. Specifically, **STR-V-53** showed weak inhibition of HDAC 8 and no inhibition of HDAC 11 at the maximum tested concentration of 10 μM (Fig. 1f). These two HDAC isoforms are potently inhibited by SAHA with nanomolar IC_50_s.

We then screened **STR-V-53** and all the synthesized glycosylated HDACi for their effects on the viability of three cell lines: A549 (lung adenocarcinoma), Hep-G2 (HCC), and VERO (a normal kidney epithelial cell). We used SORA and **STR-V-48**, a previously disclosed non-glycosylated compound, as controls^27^. We observed that SORA is about three times more cytotoxic to the Hep-G2 cells relative to VERO. At the same time, **STR-V-48** showed potent cytotoxicity against all three cell lines with no apparent selective toxicity to the Hep-G2 cells relative to VERO (Table S2). In contrast, class I glucosylated compound **STR-V-53** showed selective toxicity against Hep-G2 HCC cells over other cell lines (Fig. 1g). Additionally, compound **STR-V-167**, the peracetylated analog of **STR-V-53**, largely retained the Hep-G2 selectivity of **STR-V-53** (Table S2). Subsequently, we evaluated the effects of **STR-V-53** on cell proliferation using a panel of human HCC cell lines (Huh-7, SK-Hep-1, and Hep3B) as well as Kupffer cells (liver resident macrophages) by MTS assay. We observed that **STR-V-53** is 7- to 16-fold more cytotoxic against these HCC cell lines relative to the Kupffer cells (Fig. 1g). We also screened the other glycosylated compounds against A549, Hep-G2, and VERO and found that they elicit varying degree of potency and selective toxicity against Hep-G2 cell. Specifically, the cinnamate-derived class III compounds **STR-V-105** and **STR-V-115** showed weak anti-proliferative effects against Hep-G2 while most of these compounds showed Hep-G2-selectivity (VERO/Hep-G2) that is somewhat inferior to that of **STR-V-53** (Table S2). The exceptions are **STR-165** and **STR-176**, which showed 8- and 10-fold Hep-G2-selectivity, respectively (Table S2). Based on their weak HDAC and cell proliferation inhibition activities, we de-emphasized further evaluation of the class III compounds. Relative to SORA, a clinically approved anti-HCC agent and a prototypical HDACi such as **STR-V-48**, this whole cell data suggests that compounds **STR-V-53**, its peracetylated analog **STR-V-167**, **STR-V-165** and **STR-V-176** could be lead selective anti-HCC agents.

To further investigate the cancer cell-type selectivity of these glycosylated HDACi, we screened **STR-V-53** (10 µM) in the NCI 60 panel. It was found that **STR-V-53** has negligible effect on the proliferation of all cancer cells in NCI panel (Fig. S3), which is lacking HCC cell lines, showing a mean cell growth of 99.6%. The anti-proliferative effect of **STR-V-53** against Hep-G2 cells and its lack of growth inhibition in the NCI-60 panel strongly support the HCC cell-selective attribute of **STR-V-53** and possibly other lead glycosylated HDACi identified herein.

### HCC cells uptake glycosylated HDACi via GLUT-2

To investigate the role of GLUT-2 in the uptake of **STR-V-53** by Hep-G2, we pharmacologically blocked GLUT-2 with a selective inhibitor, phloretin (Ph)^28^, and assess the effect of this blockage on cell cytotoxicity using MTS assay. We observed that Ph selectively mitigated the cytotoxicity of **STR-V-53** against the Hep-G2 cell line with no apparent effect on the cytotoxicity of SAHA (Fig. 2a-b). Against the VERO cell line, we did not observe any significant change in the effects of **STR-V-53** or SAHA in the presence of Ph (Fig. 2c-d). This matches our expectation because Hep-G2 cells overexpress GLUT-2 while VERO has limited expression of GLUT-2^28^.

**Figure 2.**
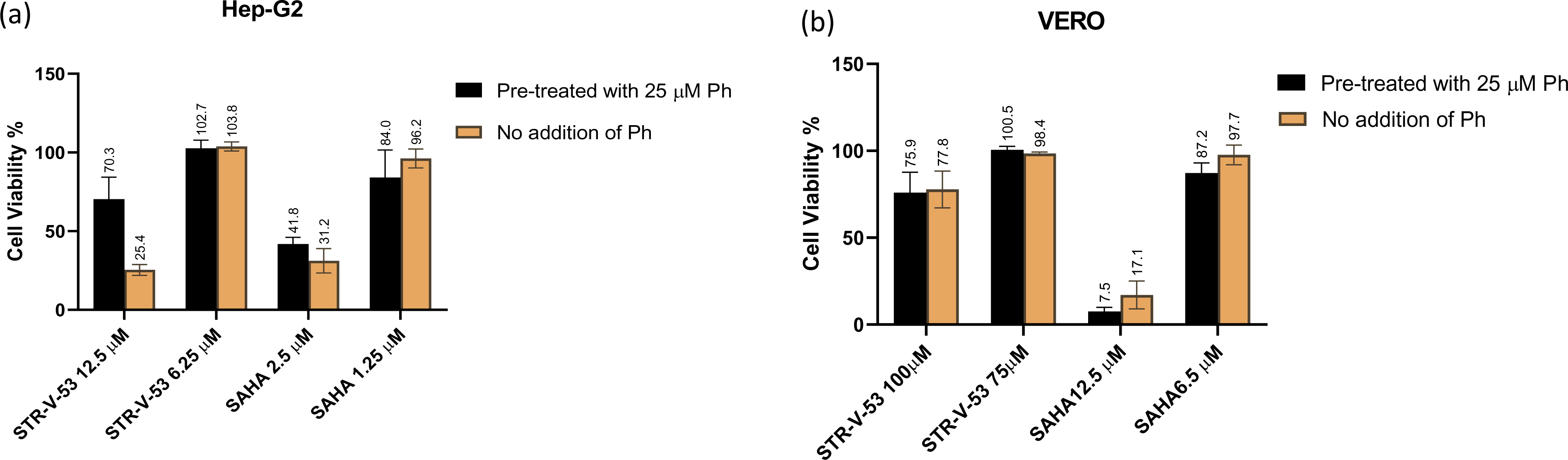
Blockade of GLUT-2 attenuates the cytotoxicity of STR-V-53 against Hep-G2 HCC cells. Hep-G2 and VERO were treated with Phloretin (Ph) for 24 h before incubation with **STR-V-53** or SAHA. (a) Hep-G2 treated with **STR-V-53** or SAHA with or without Ph. (b) VERO treated with **STR-V-53** or SAHA with or without Ph.

### Glycosylated HDACi show intracellular HDAC inhibition

To determine the contributions of HDAC inhibition to the antiproliferative activities of the glycosylated HDACi compounds, immunoblotting was used to investigate the acetylation status of histone H4 and α-tubulin, biomarkers for HDAC class I and class IIb intracellular inhibition, respectively^29^, in Hep-G2 cells in response to exposure to representative compounds **STR-V-53, STR-V-114,** and **STR-I-195**. We used SAHA as a positive control for HDAC inhibition and GAPDH level as a protein loading control. As expected, the exposure of cells to **STR-V-53** and **STR-V-114** at ¼ IC_50_ and ½ IC_50_ induced accumulation of acetylated H4 and acetylated tubulin in a similar manner to SAHA (Figs. 3a and S4i). On the other hand, **STR-I-195** at 0.5 µM did not show significant acetylation of H4 and α-Tubulin, but at 1.25µM and 2.5 µM showed significant acetylation on α-Tubulin, and mild acetylation on H4 (Fig. S4ii). This data indicates that compounds **STR-V-53, STR-I-195,** and **STR-V-114** intracellularly inhibit HDAC isoform I and HDAC 6 at low micromolar concentrations.

**Figure 3.**
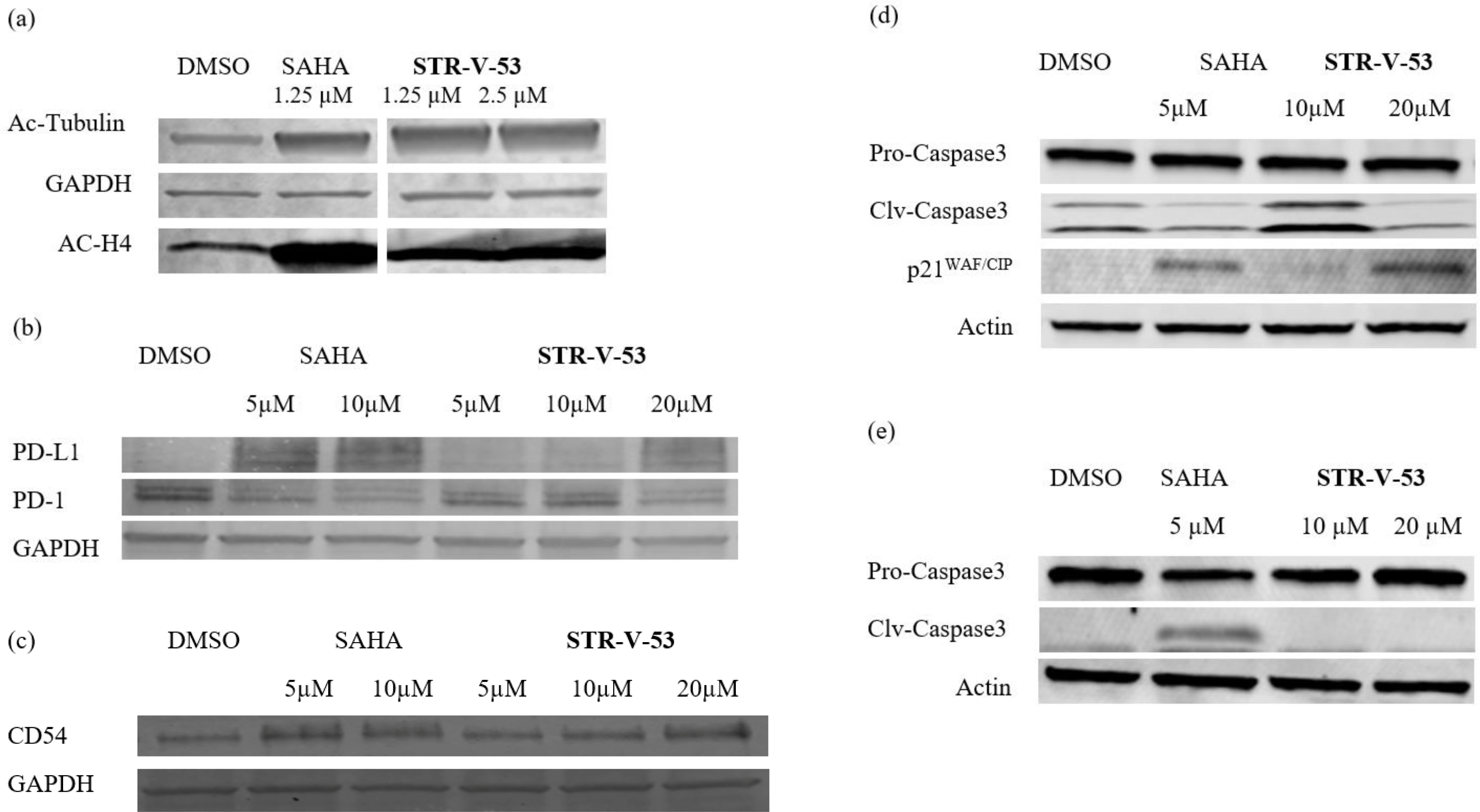
Effects of STR-V-53 on intracellular markers of HDAC inhibition, anti-tumor immune response, and apoptosis. (a) **STR-V-53** upregulates the levels of acetylated H4 and tubulin in Hep-G2 cells. Hep-G2 cells were treated with DMSO or 0.1% DMSO solution of SAHA (1.25µM) and **STR-V-53** (1.25µM, 2.5µM). Cells were treated for 5 h before lysis. Data are from two independent experiments. (b) **STR-V-53** upregulates PD-L1 and downregulates PD-1 expressions in Hep-G2 cells. Cells were treated with DMSO or 0.1% DMSO solution of SAHA (5µM and 10 µM) and **STR-V-53** (10 and 20 µM) for 24 h. (c) **STR-V-53** and SAHA caused upregulation of the levels of CD54 in the Hep-G2 cell line. Cells were treated with DMSO or 0.1% DMSO solution of SAHA (5µM and 10 µM), **STR-V-53** (5, 10, and 20 µM) for 24 h. (d and e) **STR-V-53** caused upregulation of p21^WAF/CIP^, and selectively induced apoptosis in tumor cells (Hep-G2 vs. Vero) evidenced by pro-caspase-3 activation (clv-caspase), while SAHA, a pan-HDACi, indiscriminately caused apoptosis in both tumor and normal cells. Cells were treated with DMSO or 0.1% DMSO solution of SAHA (5µM) and **STR-V-53** (10 and 20 µM).

### STR-V-53 upregulates PD-L1 and downregulates PD-1 expression in HCC

One of the consequences of histone deacetylase inhibition is the alteration of tumor immunogenicity in favor of enhanced anti-tumor immune responses^30,31^. Specifically, HDACi robustly upregulates PD-L1 in human and murine cell lines and patient tumors^32,33^. To investigate the effects of our glycosylated HDACi on immune checkpoint molecules, we used immunoblotting to probe for the expression levels of PD-L1 and PD-1 in Hep-G2 cells incubated with lead compound **STR-V-53** for 24 h. We used SAHA and DMSO as positive and negative controls, respectively. We found that both SAHA and **STR-V-53** upregulate the PD-L1 and downregulate the PD-1 expression in this cell line (Figs 3b and S5a).

### STR-V-53 upregulates CD54 (ICAM-1) in HCC cells

HDACi regulates the expression levels of other tumor antigens such as CD40, CD54 (ICAM-1), CD80, and CD86 in tumor cells^34–37^. More specifically, the repression of CD54 expression is another strategy tumors use to escape the immune response^38^. HDACi have been shown to induce upregulation of CD54^34–38^. To investigate the effect of our glycosylated HDACi on the expression of CD54, we incubated Hep-G2 cells with **STR-V-53** at IC_50_ and 2x IC_50_ for 24 h.

We used immunoblotting to determine the effect of this treatment on CD54 expression. Cells treated with SAHA and DMSO were positive and negative controls, respectively. We found that both **STR-V-53** and SAHA upregulated the expression of CD54 in Hep-G2 cells (Figs. 3c and S5b). This result indicates that, similar to other HDACi, **STR-V-53** possesses anti-tumor immune-modulatory activities.

### STR-V-53 selectively induces apoptosis in HCC cells

HDAC inhibition could cause cell apoptosis through caspase cleavage and p21^WAF/CIP^ upregulation^39^. To probe if our compounds elicit a similar phenotype, we investigated the effects representative examples on the cellular levels of cleaved Caspase 3 and p21^WAF/CIP^ through immunoblotting. Specifically, Hep-G2 cells (1 x10^6^ count/well) were treated with SAHA, **STR-V-53,** and **STR-I-195** at IC_50_ or 2x IC_50_ for 18 h prior to the cell lysis. Western blotting on the cell lysates revealed that **STR-V-53** caused a significant upregulation of cleave Caspase 3/Pro-caspase 3 ratio, while SAHA and **STR-I-195** also showed cleavage of Caspase 3 (Figs. 3d, S6a and S7a). We found that p21^WAF/CIP^ was also upregulated significantly at higher concentrations of **STR-V-53** and **STR-I-195** (Figs. 3d, S6a and S7a). These data show that **STR-V-53** and **STR-I-195** act through HDAC inhibition, similar to SAHA, to induce apoptosis in Hep-G2 cells.

Interestingly, **STR-V-53** or **STR-I-195** did not induce cleavage of Caspase 3 in VERO cells even at the same concentrations that would induce Caspase 3 activation in Hep-G2 cells (Figs. 3e, S6b and S7b). In contrast, SAHA induced cleavage of Caspase 3 in VERO cells (Figs. 3e, S6b and S7b). This data shows that these glycosylated HDACi are selectively toxic to HCC cells.

### STR-V-53 causes cell cycle arrest at the S stage

To determine if the HCC cell selective cytotoxicity of **STR-V-53** results from its perturbation of the cell cycle pattern, we evaluated the effect of **STR-V-53** on the cycle progression by flow cytometry using SAHA as a control (Fig. 4). We observed that SAHA (5 µM) and **STR-V-53** (15 µM) caused significant S phase arrest of Hep-G2 HCC cells after 48 h treatment. This data shows that the **STR-V-53** may act like SAHA to induce cell death at the concentration investigated.

**Figure 4.**
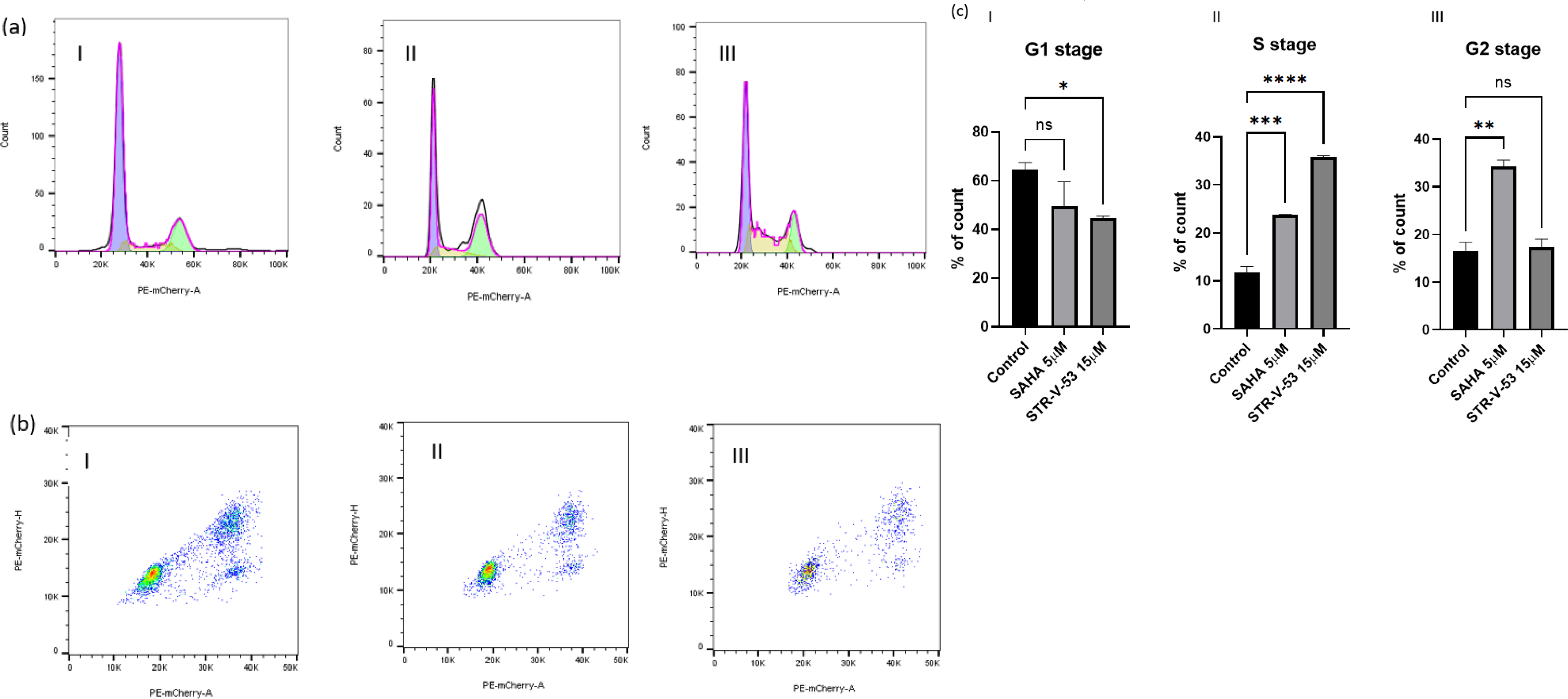
Effect of STR-V-53 treatment on HCC cell cycle progression. (a-c) Human Hep-G2 HCC cells were cultured until 80% confluence in a 10 cm petri dish. The cells were serum-starved overnight before drug treatment. Then the cells were treated with 10 mL of 0.1% DMSO medium and 0.1% DMSO solution of SAHA (5 µM) or **STR-V-53** (15µM), respectively, for 48 h. The control group (ai); SAHA (5 µM) treated group (aii) and **STR-V-53** (15µM) treated group (aiii and biii); and (c) quantification of the cell cycle data from two independent experiments. Bars show mean plus standard deviation; * P < 0.0332; ** P < 0.0021; ***P<0.0002; ****P<0.00001.

### STR-V-53 treatment suppresses HCC growth in vivo

Based on the data from the aforementioned *in vitro* studies, we selected **STR-V-53** as a candidate for further evaluation in xenograft and murine models of HCC. Before the efficacy studies, we first determined the plasma stability (murine and human) and maximum tolerated dose (MTD) of **STR-V-53** in healthy C57Bl/6 mice. We found that **STR-V-53** plasma stability is species-dependent, with a half-life of >20 h and 46 min in human and murine plasma samples, respectively (Fig. S8). For MTD determination, **STR-V-53** was administered via i.p. injection using a formulation containing excipients found in FDA-approved drugs – dimethylacetamide (DMA)/Cremophor RH 40 (CRH)/Water (10%/20%/70%) – that we developed^18^. We exposed cohorts of animals (6 per group, equal number of both sexes) to the drug at two concentrations – 50 mg/kg, 100 mg/kg (limiting injection volume to 100 µL per dose) – daily for 6 days. Using body weight as an indicator of toxicity, we observed no overt toxicity, as **STR-V-53** caused no significant body weight loss at 100 mg/kg (Fig. 5a). Moreover, we found that **STR-V-53** has a clean toxicity profile in *vitro* cytochrome P450 and hERG inhibition assays, showing no inhibition at the maximum tested concentration of 100 μM. Based on this MTD and preliminary toxicity data, we selected 25 and 50 mg/kg body weight as suitable dose levels for *in vivo* efficacy studies on **STR-V-53**.

**Figure 5.**
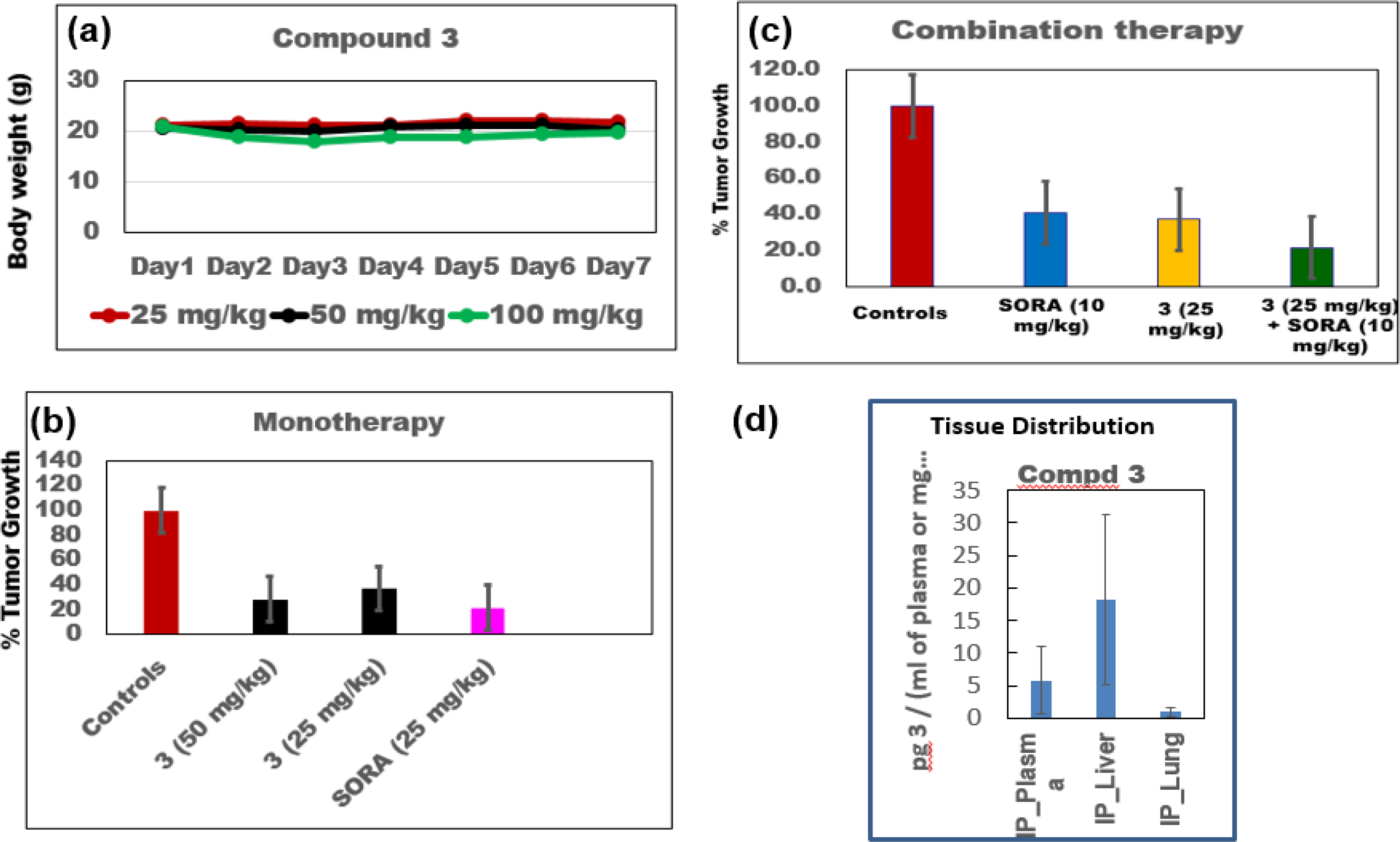
*In vivo* efficacy and tissue distribution of lead compound STR-V-53 (3). (a) **STR-V-53 (3)** showed no adverse effects on mice, based on the change in body weight, at doses as high as 100 mg/kg body weight. There are 6 mice per group, an equal number of both sexes. (b) **STR-V-53 (3)** robustly suppressed tumor growth as a standalone agent in an orthotopic xenograft model using Hep-G2 human HCC. (c) The combination of **STR-V-53 (3)** with SORA enhanced the potency of SORA. There are 10 mice (5 male and 5 female) per treatment group. (d) **STR-V-53 (3)** selectively accumulated in the liver tissues. Data from 4 randomly selected mice (2 male and 2 female).

Next, we evaluated the effects of **STR-V-53** on tumor growth in an orthotopic murine model of Hep-G2-Red-FLuc Bioware^®^ Brite Cell Line (PerkinElmer) in mice. Mice (6-8 weeks old) were orthotopically implanted through a direct intrahepatic artery injection with Hep-G2-Red-FLuc cells according to published protocols^40^. We used the Red-Fluc expression to confirm tumor implantation by bioluminescent imaging using IVIS. Treatment began after imaging confirmed the establishment of intrahepatic tumors (approx. 7 days). Mice were grouped into cohorts with similar average relative chemiluminescence. Tumor-bearing mice in the treatment groups were injected daily with 200 µL of solution of **STR-V-53**, SORA, or a combination of **STR-V-53** and SORA for 21 days via the i.p. route. No treatment group received vehicle only. After treatment, mice were imaged again, sacrificed, and liver samples were harvested to determine the effects of treatment on tumor size by measurement with calipers. We found that **STR-V-53** efficiently suppressed HCC tumor growth at 50 mg/kg and 25 mg/kg to a similar extent as SORA. **STR-V-53** at 25 mg/kg and 50 mg/kg induced tumor growth inhibition (TGI) of 60% and 68%, respectively (Fig. 5b). Moreover, the combination therapy of SORA (10 mg/kg) and **STR-V-53** (25 mg/kg) is more efficacious than either drug as a standalone agent. Relative to the effect of each agent alone, the combination of **STR-V-53** (25 mg/kg) and SORA (10 mg/kg) additively reduced tumor volume with a TGI of approximately 80% (Fig. 5c).

To determine if liver tissue accumulation plays a role in the observed *in vivo* efficacy of **STR-V-53**, we used LC-MS to evaluate the distribution of **STR-V-53** in selected tissues (liver and lungs) and plasma. We analyzed samples from healthy mice exposed to **STR-V-53** (i.p. injection) at 25 mg/kg for 8 hr. In the lungs, the level of **STR-V-53** is below the detection limit of our instrument for both sexes. The average level of **STR-V-53** in circulation in the plasma of healthy mice was ∼10 pg/mL 8 hr post-injection. In contrast, the level of **STR-V-53** in the liver was approximately 4 times its plasma level (Fig. 5d). Of note, we found a sex dependence in the plasma and liver distribution of **STR-V-53**, with male mice showing significantly higher levels (Fig. S9).

HDACi may elicit potent immunomodulatory activity through multiple mechanisms^30–33,41^, and synergize with immunotherapy, one of the current standards of care for HCC^42,43^. Since the glycosylated HDACi disclosed herein showed HCC cell-selectivity, anti-tumor immune-modulatory activities, and dual antibody blockade of VEGF and PD-L1 pathways is less toxic than SORA alone, their combination with anti-PD1 therapy might be safe and highly effective. Indeed, when we investigated the effect of **STR-V-53** (25 mg/kg, daily, i.p.) on the efficacy of anti-PD-1 antibody (10 mg/kg, 3 doses per week, i.p.) in a syngeneic murine model, we found that combination of **STR-V-53** with anti-PD-1 antibody, relative to anti-PD-1 antibody alone, more potently suppressed tumor growth, increased overall survival and caused remarkably durable responses in ∼40% of the mice (Fig. 6a-c). Remarkably, we detected no significant overt toxicity from **STR-V-53**, alone or combined with a PD-1 antibody, during the treatment period (Fig. 6d). The HCC cell selectivity, favorable toxicity profiles, and increased liver exposure of **STR-V-53** are unique attributes that potentially contributed to its *in vivo* efficacy.

**Figure 6.**
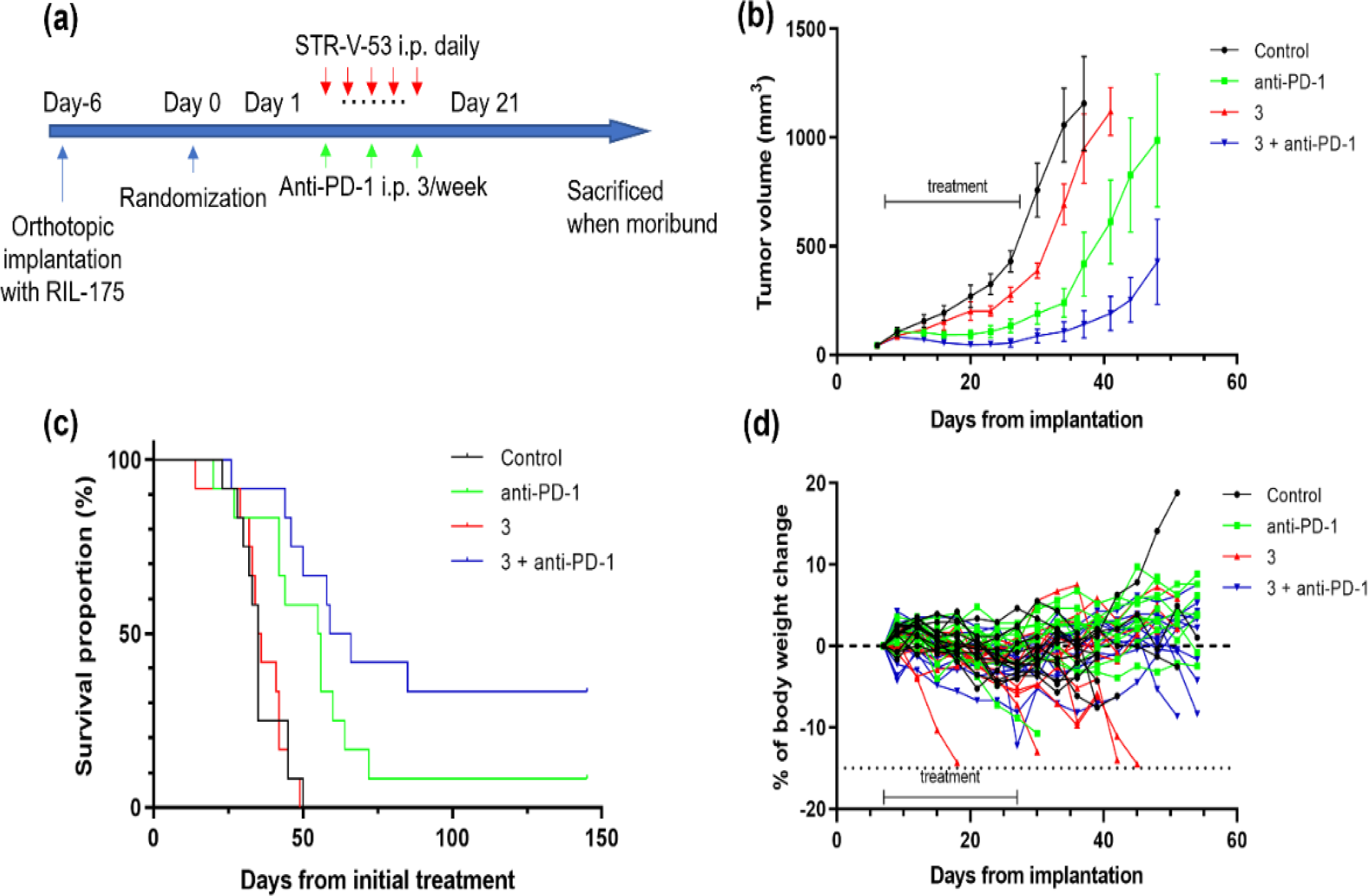
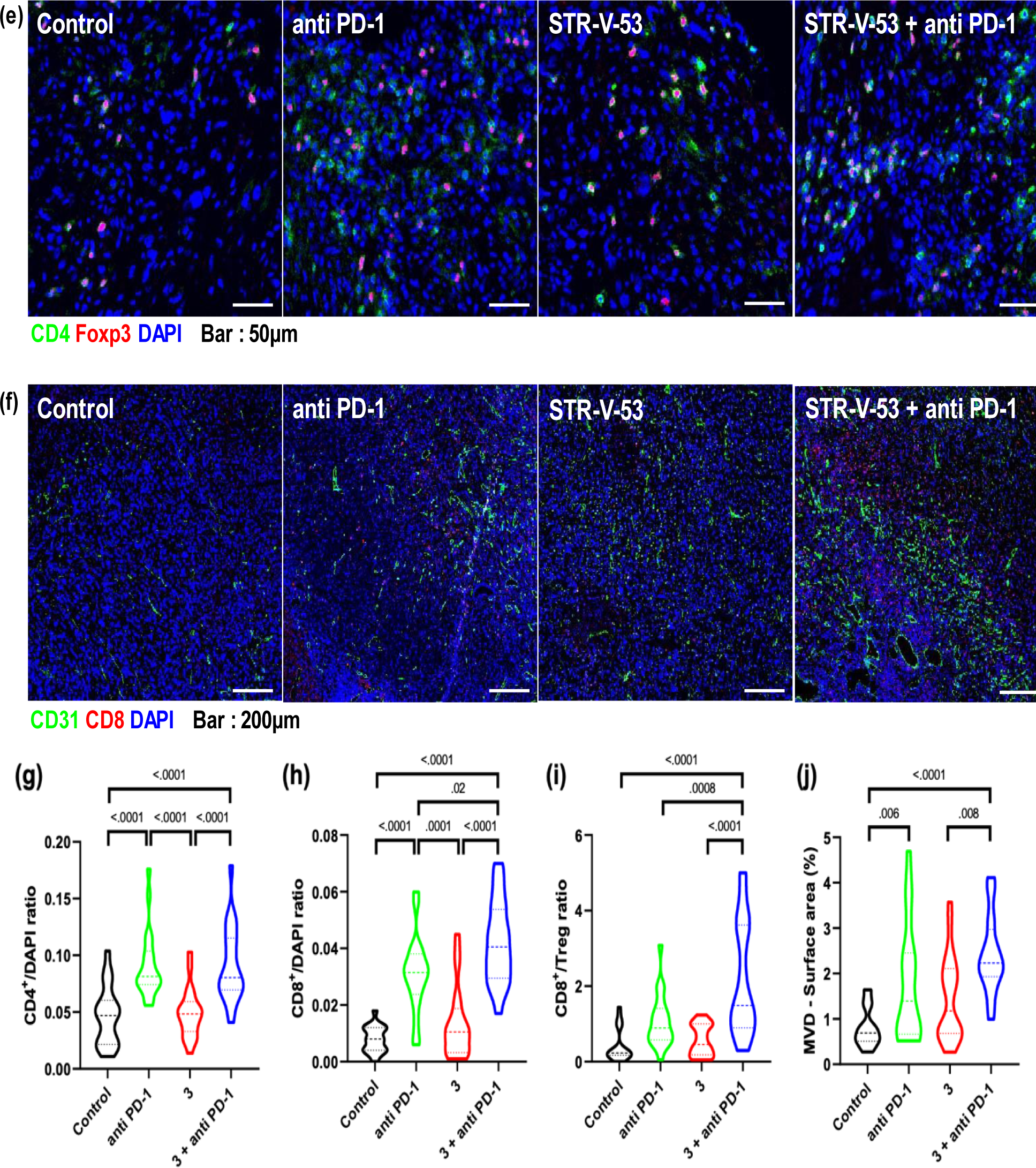
Combined treatment with STR-V-53 (3) and anti-PD-1 antibody shows efficacy in an orthotopic murine model of HCC. (a) Treatment schedule for survival studies in the RIL-175 murine HCC model. (b) Tumor growth was significantly delayed in the group treated with a combination of **STR-V-53 (3)** and anti-PD1 therapy compared with the other groups (n=12). (c) Kaplan-Meier survival distributions in mice bearing orthotopic RIL-175 murine HCC and treated with **STR-V-53 (3)** alone, anti-PD-1 antibody alone, or their combination compared to IgG control. The combination of **STR-V-53 (3)** and anti-PD1 therapy significantly prolonged overall survival (n=12). (d) Body weight change of tumor-bearing mice in the four treatment groups did not indicate toxicity. (e) Representative immunofluorescence for CD4^+^ and Foxp3^+^ TILs in tumor tissues from the 4 treatment groups. (f) Representative immunofluorescence for the endothelial marker CD31 and CD8^+^ TILs in tumor tissues from the 4 treatment groups. The frequencies of tumor-infiltrating CD4^+^ T cells (g) and CD8^+^ T cells (h), shown as the mean [SD] ratio of each positive cell counts to DAPI positive cells per field (320*320 mm square area). (i) The ratios of CD8^+^ T cell counts to Tregs (CD4^+^Foxp3^+^) from the 4 treatment groups. (j) Quantification of tumor MVD, shown as area fraction covered by vessels per field (1.25*1.25 mm square area). Scale bars: 50 μm (e) and 200 μm (f). *p-value* from Kruskal-Wallis test (two-sided). TIL = tumor-infiltrating lymphocytes; MVD = microvessel density.

Immunofluorescence (IF) evaluation of liver tumor samples from this *in vivo* experiment also showed a significant increase in tumor-infiltrating CD4^+^ and CD8^+^ T cells in the anti-PD-1 antibody monotherapy group and in the **STR-V-53** plus anti-PD-1 antibody group (Fig. 6e-h). Analysis of the ratio of CD8^+^ T cells to Tregs and the extent of microvessel density, which are thought to correlate with a favorable response to clinical treatment, were both significantly higher in the combination treatment group and were associated with a longer survival period for the mice in this group (Fig. 6i-j).

### Transcriptomic analysis of Hep-G2 cells reveals antitumor activity of STR-V-53

To obtain transcriptome level evidence that may shed light on the HCC-selectivity and/or the enhanced potency of the combination of **STR-V-53** with anti-PD-1 antibody, we performed RNA sequencing (RNA-seq) analysis on **STR-V-53**-treated Hep-G2 cells, incubated with at IC_50_ and 2 x IC_50_ for 24 hr, relative to the untreated (DMSO) control. We first conducted gene set enrichment analysis (GSEA) using hallmark and gene ontology biological process (GOBP) gene sets to analyze the effect of **STR-V-53** vs DMSO-treated control on the Hep-G2 transcriptome^45,46^. This analysis revealed that p53 pathway was positively enriched (NES +1.6) at 2x IC_50_ while the PI3K/Akt**/**mTOR signaling (-1.7; 2x IC_50_), MYC targets v1 (-2.6; IC_50_ and - 2.3; 2x IC_50_), MYC targets v2, (-2.3; IC_50_ and -2.6; 2x IC_50_), and G2M checkpoint pathways, (- 2.9; IC_50_ and -2.2; 2x IC_50_) were negatively enriched (Fig. 7). Negative enrichment of the G2M checkpoint pathway supports the finding that **STR-V-53** induced S phase arrest in Hep-G2 cells (see Fig. 4).

**Figure 7.**
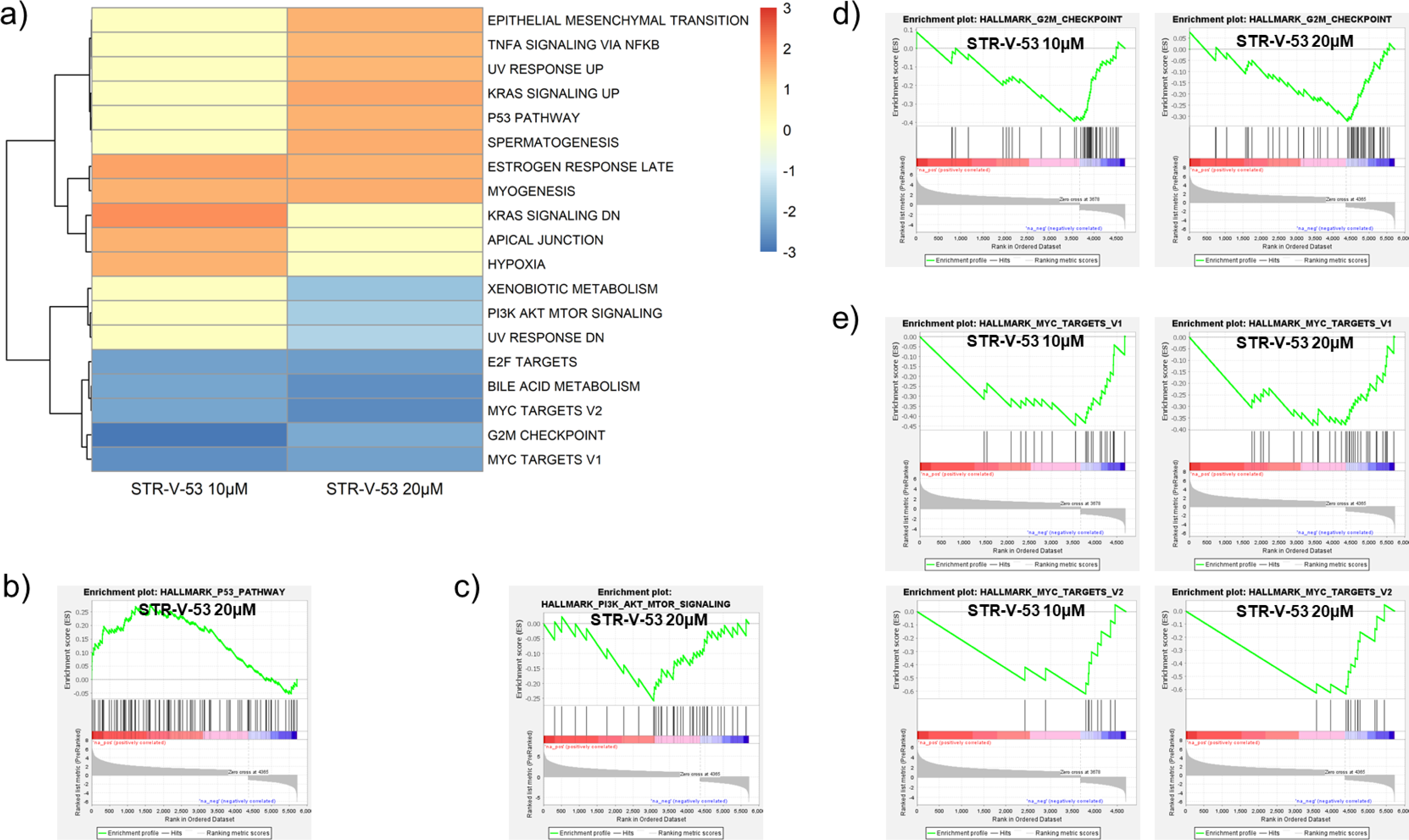
Hallmark GSEA results of STR-V-53 treatment to human HCC cells. a) Heatmap of the normalized enrichment scores (NES) of significantly enriched hallmark gene sets from the GSEA Molecular Signatures Database (MSigDB) (p < 0.05, FDR < 0.25). b) p53 pathway enrichment plot for **STR-V-53** 20µM (NES = 1.6). c) PI3K-Akt-mTOR signaling enrichment plot for **STR-V-53** 20µM (NES = –1.7). d) G2M checkpoint enrichment plots for **STR-V-53** 10µM (NES = –2.9) and 20µM (NES = –2.2). e) MYC targets v1 and v2 enrichment plot for **STR-V-53** 10µM (NES = –2.6, –2.3) and 20µM (NES = –2.3, –2.6).

Additionally, GOBP GSEA revealed negative enrichment (–1.8) of glucose-regulated transcription genes at both IC_50_ and 2x IC_50_ (Fig. 8). Other related pathways negatively enriched by **STR-V-53** at 2x IC_50_ include positive regulation of glucose import (–1.6), positive regulation of glucose transmembrane transport (–1.6) and the cytokine-mediated signaling pathway (-1.4) (Fig. 8).

**Figure 8.**
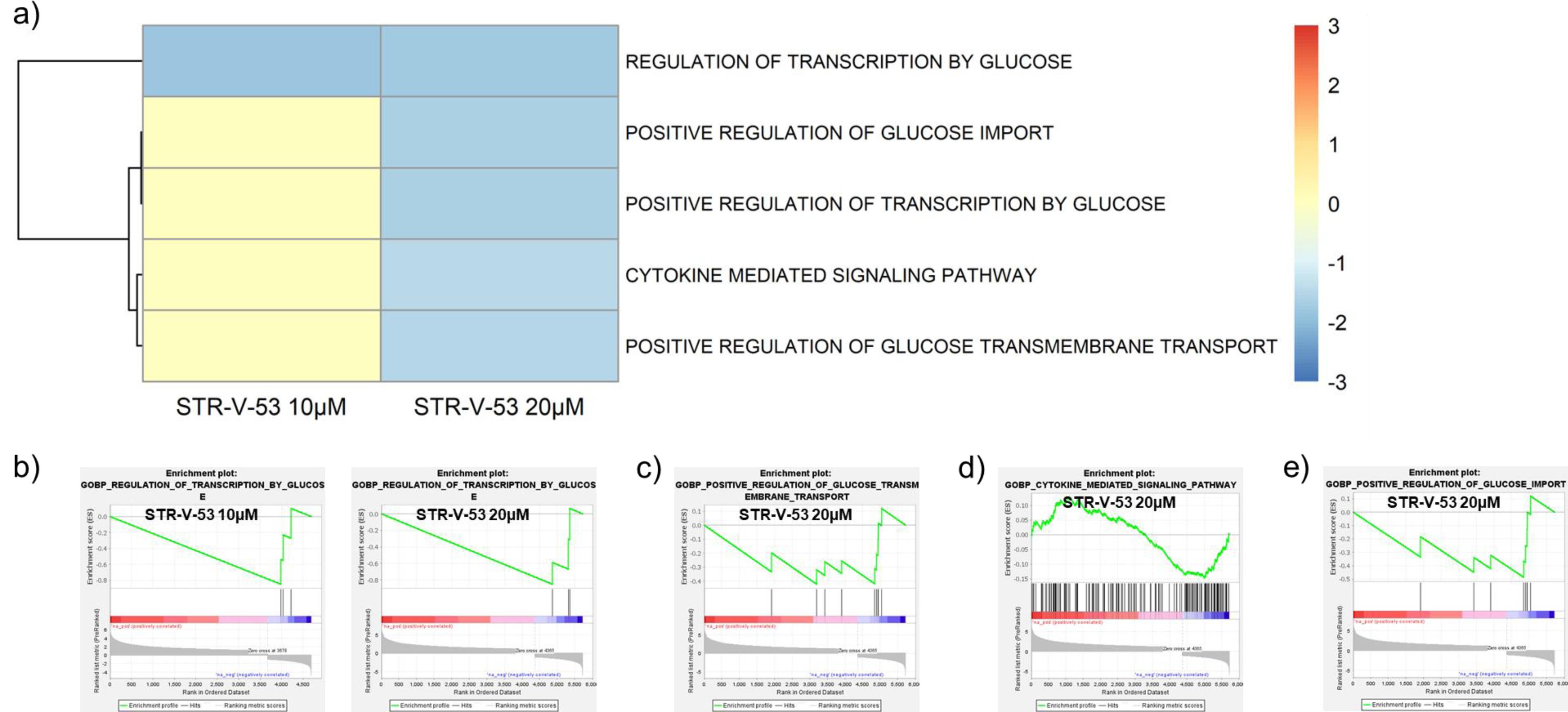
Gene Ontology-Biological Processes (GOBP) GSEA results of STR-V-53 treatment to HCC cells. a) Heatmap of the normalized enrichment scores (NES) of significantly enriched GOBP gene sets selected from the GSEA Molecular Signatures Database (MSigDB) (p < 0.05, FDR < 0.25). b) Regulation of transcription by glucose enrichment plot for **STR-V-53** 10µM (NES = –1.8) and 20µM (NES = –1.8). c) Positive glucose import enrichment plot regulation for **STR-V-53** 20µM (NES = –1.6). d) Cytokine-mediated signaling pathway enrichment plot for **STR-V-53** 20µM (NES = –1.4). e) Positive regulation of glucose transmembrane transport enrichment plot for **STR-V-53** 20µM (NES = –1.5). Of note, the negative enrichment of glucose transport genes is not due to a uniform downregulation of GLUTs. We observed that within 24 h of uptake into the cells, **STR-V-53** caused the downregulation of GLUTs 1, 2, 9, and 14, which may be compensated for by its concomitant induction of the upregulation of GLUTs 3, 4, 6, and 12 (Fig. S10). Nevertheless, the **STR-V-53**-induced negative enrichment of the glucose-regulated transcription will likely impair the ability of the Hep-G2 cells to utilize glucose to fuel growth.

Subsequently, we focused our analysis on specific genes of interest, including gene signatures for HDAC inhibition and immune response pathways. HDAC inhibition has been linked to the regulation of specific genes, and representative examples are shown in Figure 9 for clarity^47^. In a similar manner to other HDACi, **STR-V-53** at both concentrations significantly upregulates DKK-1 (+6.5, IC_50_; p < 4.5×10^-76^ and +6.2, 2x IC_50_; p < 6.5×10^-69^), providing further confirmation of its class I and II HDAC inhibition activities which have been linked to DKK-1 upregulation in several cancer types^48,49^. **STR-V-53** also upregulates CDKN1A (p21), log2 fold change +3.1 (IC_50_) and +3.3 (2x IC_50_), correspondingly supporting apoptosis-induced HCC cell death; MT1X, log2 fold change +2.9 (IC_50_) and +3.4 (2x IC_50_); PDCD1, log2 fold change +2.1 (IC_50_) and +3.0 (2x IC_50_); TUBA1A, log2 fold change +4.7 (IC_50_) and +5.4 (2x IC_50_), while downregulating ANP32B, log2 fold change -1.8 (IC_50_) and +-2.3 (2x IC_50_); TYMS, log2 fold change -3.2 (IC_50_) and -3.6 (2x IC_50_), indicating disruption of DNA synthesis and providing further confirmation of **STR-V-53**-induced S phase arrest^47,50^.

**Figure 9.**
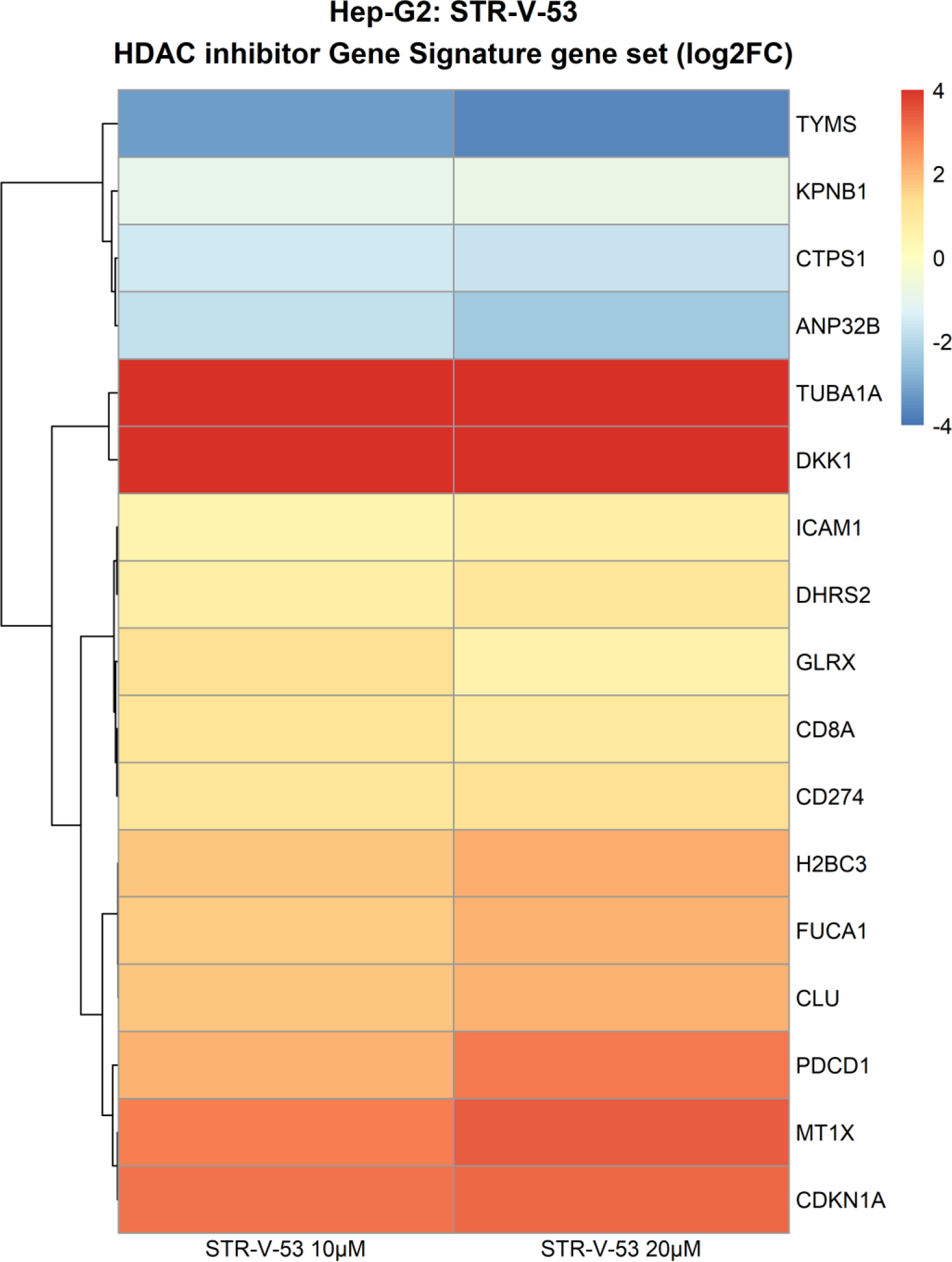
STR-V-53 effect on signature genes influenced by histone deacetylase (HDAC) inhibition. TUBA1A (log2 fold change 4.7, 10µM, and 5.4, 20µM) and DKK-1 (log2 fold change 6.5, 10µM, and 6.2, 20µM) were substantially upregulated, and TYMS (log2 fold change –3.2, 10µM, and –3.6, 20µM) downregulated.

HDAC inhibition is also associated with the upregulation of immune response pathways, making them promising candidates for combinational therapy with immunotherapy agent^51,52^. We noticed that **STR-V-53** upregulates the MHC (HLA) complex, CD8+, and PD-1 (PDCD1) /PD-L1 (CD274), all genes that are upregulated by class I and II HDACi (Fig. 10a)^52^. The upregulation of PD-1 is inconsistent with Western blot data (Fig. 3b). This discrepancy may be due to different drug incubation times used in these experiments, as immune checkpoint expression is inducible and dynamic. GOBP immune gene sets were selected for additional enrichment analysis^46^. The innate immune response in mucosa, 2.1 (IC_50_) and 2.3 (2x IC_50_), antimicrobial humoral immune response mediated by antimicrobial peptide, 1.9 (IC_50_) and 1.9 (2x IC_50_), and humoral immune response gene sets, 1.9 (IC_50_) and 1.7 (2x IC_50_), were positively enriched (Fig. 10b-e). Enrichment of these gene sets provides strong indication for the mechanistic basis for the enhanced potency of the combination of **STR-V-53** with anti-PD-1 antibody^53^.

**Figure 10.**
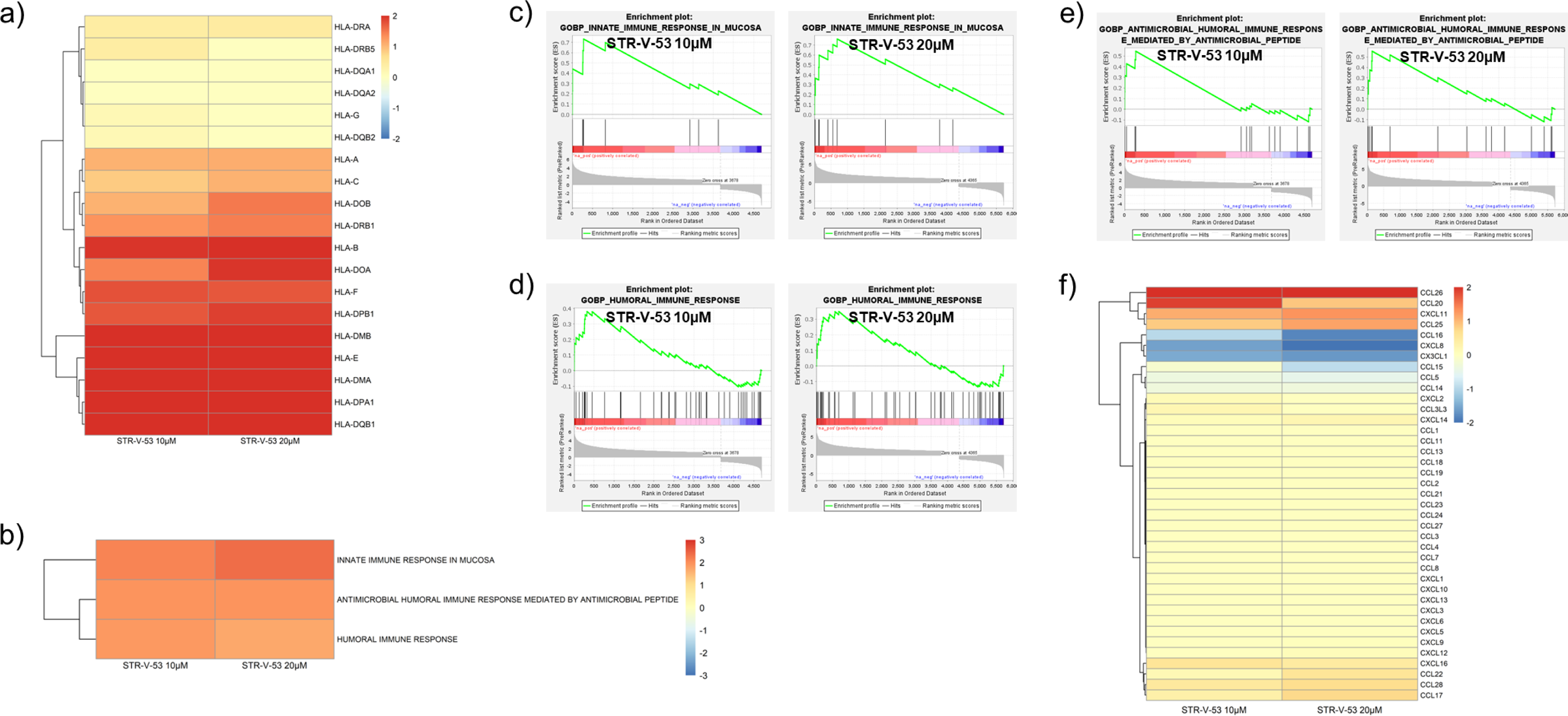
STR-V-53 effect on immune response pathways and gene patways. a) Log2 fold change heatmap of major histocompatibility complex/ human leukocyte antigen (HLA) class I genes (HLA-A, HLA-B, HLA-C, HLA-E and HLA-F) and class II genes (-DMA, -DMB, -DOA, -DOB, -DPA1, -DPB1, -DQB1 and -DRB1) determined to be upregulated by **STR-V-53**. b) Heatmap of the normalized enrichment scores (NES) of significantly enriched GOBP immune gene sets (p < 0.05, FDR < 0.25). c) Innate immune response in mucosa enrichment plot for **STR-V-53** 10µM (NES = 2.1) and 20µM (NES = 2.3). d) Humoral immune response enrichment plot for **STR-V-53** 10µM (NES = 1.9) and 20µM (NES = 1.7). e) Antimicrobial humoral immune response mediated by antimicrobial peptide enrichment plot for **STR-V-53** 10µM (NES = 1.9) and 20µM (NES = 1.9). f) Log2 fold change heatmap of chemokines up/downregulated by **STR-V-53**: CCL26 (log2 fold change 2.1, 10µM, and 2.9, 20µM), CCL20 (log2 fold change 1.9, 10µM, and 0.9, 20µM), CXCL11 (log2 fold change 1.1, 10µM, and 1.3, 20µM), CCL25 (log2 fold change 0.9, 10µM, and 1.2, 20µM), CCL16 (log2 fold change -1.0, 10µM, and –1.8, 20µM), CXCL8 (log2 fold change –1.6, 10µM, and -2.0, 20µM), and CX3CL1 (log2 fold change –1.5, 10µM, and –1.7, 20µM).

Further probing into the effect of **STR-V-53** on the expression of chemokines revealed a selective downregulation of CCL16, log2 fold change –1.0 (IC_50_) and –1.8 (2x IC_50_), CXCL8, log2 fold change –1.6 (IC_50_) and -2.0 (2x IC_50_), and CX3CL1, log2 fold change –1.5 (IC_50_) and - 1.7 (2x IC_50_) and upregulation of CCL26, log2 fold change +2.1 (IC_50_) and +2.9 (2x IC_50_), CCL20, log2 fold change +1.9 (IC_50_) and +0.9 (2x IC_50_), CXCL11, log2 fold change +1.1 (IC_50_) and +1.3 (2x IC_50_), and CCL25, log2 fold change +0.9 (IC_50_) and +1.2 (2x IC_50_) (Fig. 10f). The expression of CCL16, CXCL8 and CX3CL1 has been implicated in HCC cell invasion, metastasis, migration and M2 polarization of tumor-associated macrophages, while the suppression of is associated with improved patient prognosis^54–46^. In contrast, CCL20, CCL26, CCL25 and CXCL1 have been shown to play multifaceted roles, including oncogenic functions and predictors of better prognosis, in several tumor types^55,57–60^. However, chemokines are involved in a network of interactions to regulate tumor growth and progression. The perturbation of this network by **STR-V-53** may tilt the balance to favor the antitumor effect.

## Discussion

HCC is a frequent and lethal cancer worldwide, and available treatments have limited efficacy. Previous studies have shown that HCC is strongly driven by epigenetic dysregulation that leads to chromatin histone hypoacetylation. HDAC inhibition therapy could be a promising treatment option for HCC. However, HDACi has been ineffective against solid tumors such as HCC. In this study, we investigated a design strategy exploring the Warburg effect to deliver HDACi to HCC cells selectively. HCC cell lines and tumor samples overexpress GLUT-2, facilitating glucose and mannose uptake to support tumor growth^28^. GLUT-2 is a major facilitator of sugar transport in the hepatocytes and a prognostic factor of HCC ^22^. Our *in silico* docking analysis predicted that integrating glycoside moieties into the prototypical HDACi surface recognition group could be compatible with HDAC inhibition activities as a cohort of the resulting glycosylated HDACi bind to representative HDACs with strong binding affinities. The glycoside moieties of these compounds are also accessible for GLUT-1 binding. As expected, the glucose moiety enabled optimal GLUT-1 interaction, while the desosamine moiety, due to its fewer hydroxyl groups, has attenuated GLUT-1 binding affinity. Nevertheless, integrating these glycosides into the HDACi pharmacophore affords compounds with much higher GLUT-1 binding affinities relative to unmodified glucose and n-nonyl beta-D-glucopyranoside. This suggests that the glycosylated HDACi are better cargo for selective uptake to GLUT-1- or GLUT-2-enriched cell lines. Results for HDAC inhibition and GLUT-2 competition assays fully supported our *in silico* molecular analysis predictions.

The ability of these glycosylated HDACi as cargo for GLUT-2 was indirectly evidenced by their glycoside moiety-dependent selective toxicity against Hep-G2 relative to other cell lines. The glucosylated compounds are more selectively toxic against Hep-G2 than the mannosylated congener. Additional screening of the lead compound **STR-V-53** was performed against the NCI-60 panel and a panel of human HCC cell lines (Huh-7, SK-Hep-1 and Hep3B) as well as Kupffer cells (liver resident macrophages), which are not present in the NCI-60 panel. Our choice of these HCC cell lines was informed by their p53 expression status, which may affect their response to HDAC inhibition. Unlike Hep-G2, which expresses wildtype, Huh-7 has mutant p53, while Hep3B is p53 null. Hep-G2 (p53-WT) and Hep3B (p53-null) are responsive to HDAC inhibition while Huh-7’s (p53-mutant) response is dependent on the HDACi^44,61–62^. The robust cytotoxicity of **STR-V-53** against these HCC cell lines, and its lack of strong antiproliferative effects against the cells in the NCI-60 panel, further attest to the exquisite HCC cell-selectivity of **STR-V-53**. Furthermore, the attenuation of the cytotoxicity of the **STR-V-53**, and not that of the control compound SAHA, against Hep-G2 in the presence of GLUT-2 inhibitor provided further support that **STR-V-53** derived a significant part of its Hep-G2 cell penetration through GLUT-2-mediated transport. Further evaluation of the intracellular mechanisms revealed that **STR-V-53**, **STR-I-195,** and **STR-V-114** caused upregulation of acetylated tubulin and H4 in Hep-G2 cells, confirming their intracellularly HDACs inhibition. In addition, the lead compound **STR-V-53** and **STR-I-195** elicited similar effects as SAHA on the expression of p21^WAF/CIP^, a downstream target of intracellular HDAC inhibition, in Hep-G2 cells. **STR-V-53** also induced anti-tumor immune modulatory activities on Hep-G2 cells by upregulating the expression of CD54, PD-L1, and downregulating PD-1. Tumor expression of PD-L1 has been implicated in T cell tolerance^31,63^. Paradoxically, HDACi and PD-1 blockade combination have been shown to more effectively inhibit tumor growth in murine models and increase animal survival relative to single-agent treatments^32,33^. This observation supports Taube *et al*.’s conclusion that tumor PD-L1 expression is indicative of immune-active TME and that the expression status of immunosuppressive molecules PD-1 and PD-L2 is the most important factor closely correlated with response to anti-PD-1 immunotherapy^31^. PD-1 expression in HCC may also promote tumor growth through mechanisms independent of adaptive immunity. PD-1 knockdown suppresses HCC growth, while its overexpression enhances tumorigenesis in immune-deficient mice^64^. The upregulation of PD-L1 levels in cancer cells by HDACi has been reported for some tumor types but not in others, for example, in hormone-responsive breast cancer cells^65^. However, the downregulating effect of **STR-V-53** and SAHA on PD-1 expression is uncommon for HDACi. The downregulation of PD-1 by HDACi may further contribute to their cytotoxicity to HCC cell lines since a previous study has shown that PD-1 knockdown compromised the viability of several HCC cell lines^64^. These results indicate that our HCC-selective glycosylated HDACi **STR-V-53** acts in a similar manner to the standard HDACi in regulating the expression status of these immune checkpoint molecules. Unlike prototypical HDAC inhibitors however, the toxicity of these glycosylated HDACi is restricted to the HCC cells as **STR-V-53** and **STR-I-195** induced apoptosis in Hep-G2 cells and not in VERO cells.

We confirmed the benefit of the HCC cell-selectivity of the lead compound **STR-V-53** in *vivo* using healthy mice and two murine models of HCC. We observed that **STR-V-53** is stable in human plasma, relatively non-toxic to mice, and robustly suppressed tumor growths in an orthotopic model of HCC as a standalone agent and enhanced the potency of SORA in a combination therapy experiment. Also, **STR-V-53** induced superior antitumor effects in combination with anti-PD-1 antibodies. Immunofluorescence analysis of liver cancer samples showed a significant increase in CD4^+^ and CD8^+^ T cell infiltration into the tumor and a modifying effect on the tumor microenvironment, such as increased MVD and CD8/Treg ratio, in the combination therapy group of anti-PD-1 antibody and **STR-V-53**.

While the CD8/Treg ratio has been reported to have an impact on therapeutic efficacy^66,67^ and may be a marker that correlates well with the prolonged survival of the mice in this study, the mechanism by which HDAC inhibitors affect Tregs remains unclear and will be an important topic for future validation. Nevertheless, subsequent RNA seq analysis shed some light on the transcriptome level changes induced by **STR-V-53** to sensitize HCC cells to immunotherapy. After its uptake by Hep-G2 cells, **STR-V-53** perturbs the GLUT expression pattern by downregulating GLUTs 1, 2, 9, and 14 and upregulating GLUTs 3, 4, 6, and 12. Furthermore, **STR-V-53** induced other transcriptome reprogramming that favors HCC cell death through HDAC inhibition, impaired glucose-regulated transcription, impaired DNA synthesis, upregulation of apoptosis and stimulation of immune response pathways.

Others have shown that systemic HDACi could enhance the potency of SORA and immunotherapy in murine models of HCC^52,68–69^. However, the benefit of the HDACi/SORA combination has not been borne out in the clinic due to dose-limiting toxicities, which led to early closure of clinical trials of the combination of SORA with HDACi drugs such as SAHA [NCT01075113] and panobinostat [NCT00873002]. Because these compounds are selectively cytotoxic to HCC cells, they may be less prone to the toxic side effects observed in combination therapy studies, including the HDACi/SORA combination. Collectively, our data revealed that **STR-V-53** is a novel HDACi whose potential as a targeted anti-HCC agent merits further evaluation for clinical translation.

## Author Contributions

B.W. and S.T. performed compound synthesis. B.W., J.O.O and U.A. performed all cell cytotoxicity, intracellular mechanism and *in silico* studies. C.Q.S., R.S.A. and J.A.P. contributed to the design of and performed the *in vivo* efficacy study in orthotopic murine model of Hep-G2-Red-FLuc Bioware^®^ Brite Cell Line and determined MTD. Z.R., T.K. and D.G.D. designed and performed the combination therapy study of **STR-V-53** with anti-PD1 in a syngeneic murine model and immunofluorescence (IF) evaluation of tumor samples. A.J. performed RNA seq analysis. S.T. and D.A.G. coordinated part of this study that was performed at Sophia Bioscience, Inc. B.W., S.T., D.A.G., R.S.A., J.A.P., Z.R., T.K., D.G.D. and A.K.O. contributed to the design of the study. B.W., A.J., J.O.O, S.T., J.A.P., D.G.D., and A.K.O. contributed to the writing of this manuscript. A.K.O. conceived the concept and the study leading to this work was performed in his lab. All authors have read and agreed to the published version of the manuscript.

## Acknowledgments

This work was supported by the Vasser-Woolley Fellowship (A.K.O.), National Institutes of Health grants R01CA131217, R01CA266013 (A.K.O.), R43CA224642 (S.T.) and Georgia Institute of Technology’s Systems Mass Spectrometry Core Facility. D.G.D. is supported by National Institutes of Health grants R01CA260857, R01CA254351, R01CA247441, R03CA256764 and P01CA261669, and Department of Defense PRCRP grants W81XWH-19-1-0284 and W81XWH-21-1-0738.

## Additional information

## Supplementary information

Compound synthesis and characterization (^1^H-NMR and ^13^C-NMR spectral information), general methods, molecular modeling outputs, HDAC and cell inhibition data, Western blot data and plasma stability data. This material is available free of charge at https:/

