## Supplemental Info II for "A Novel Liver Cancer-Selective Histone Deacetylase Inhibitor Is Effective Against Hepatocellular Carcinoma and Induces Durable Responses with Immunotherapy"

[1]School of Chemistry and Biochemistry, Georgia Institute of Technology, Atlanta, GA 30332-0400, USA; [2]Sophia Bioscience, Inc. 311 Ferst Drive NW, Ste. L1325A, Atlanta, GA 30332, USA; [3]Edwin L. Steele Laboratories for Tumor Biology, Department of Radiation Oncology, Harvard Medical School & Massachusetts General Hospital, Boston, MA 02114, USA; [4]Department of Urology, Emory University School of Medicine, Atlanta, GA 30322, USA; [5]Parker H. Petit Institute for Bioengineering and Bioscience, Georgia Institute of Technology, Atlanta, GA 30332-0400, USA.

[¶]These authors contributed equally to the manuscript.

**Keywords:** Hepatocellular carcinoma; Histone deacetylase; Glycosylated histone deacetylase inhibitors; Glucose transporters; Anti-PD-1 Combination therapy.

#### Supporting information

**Figure S1.** (a) Structures of approved HDACi. (b) Structure of macrolide-based, class-I selective HDACi that preferentially accumulates within liver tissues.

a.

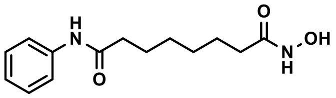


**Vorinostat**

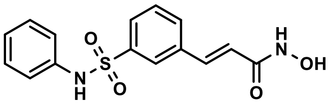


**Belinostat**

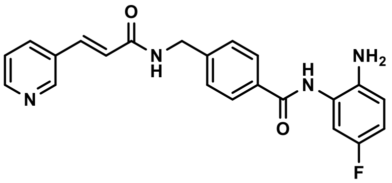


**Chidamide**

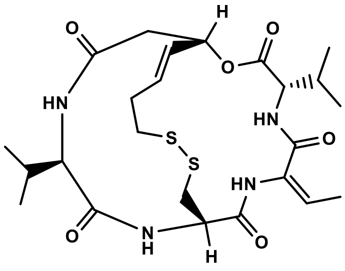


**Romidepsin**

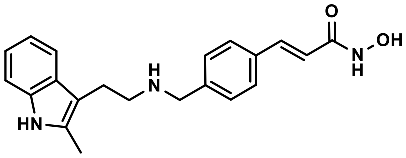


**Panobinostat**

b.

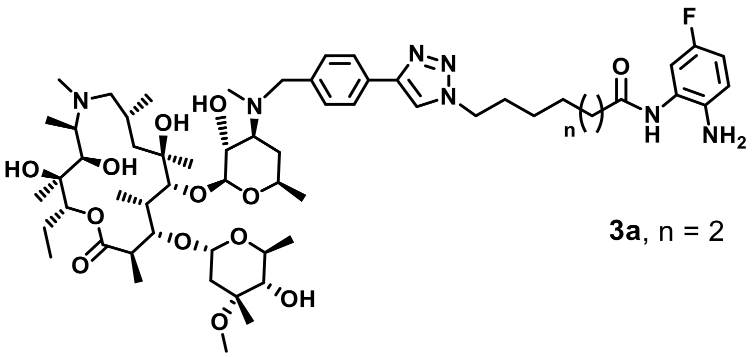


**3a**, n = 2

**Scheme 1.** Synthesis of the glucosylated HDACi **STR-V-167**, **STR-V-53**, **STR-V-114** and **STR-V-115**.

1a

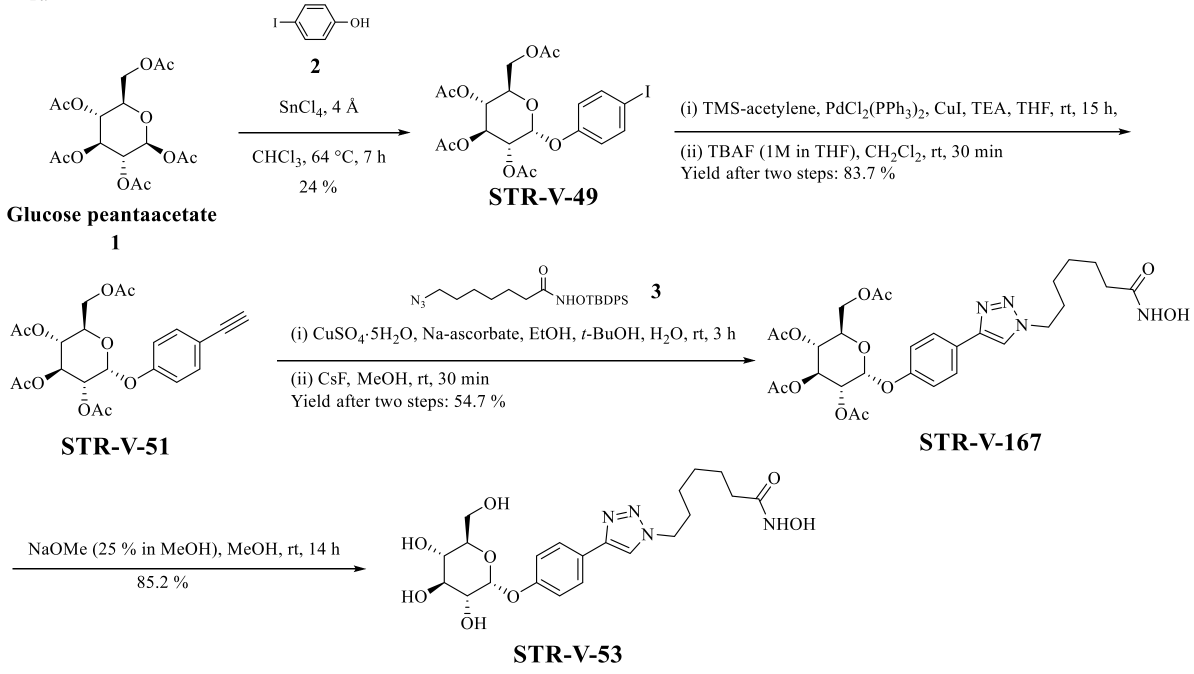


1b

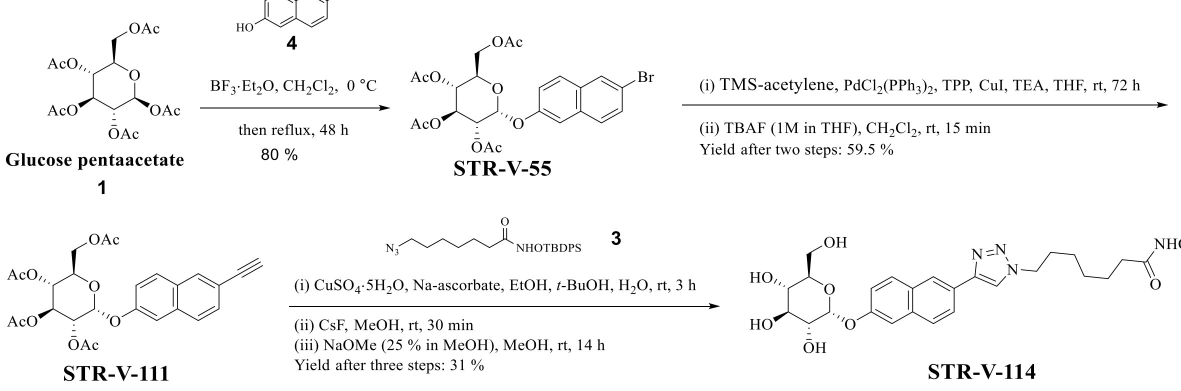


1c

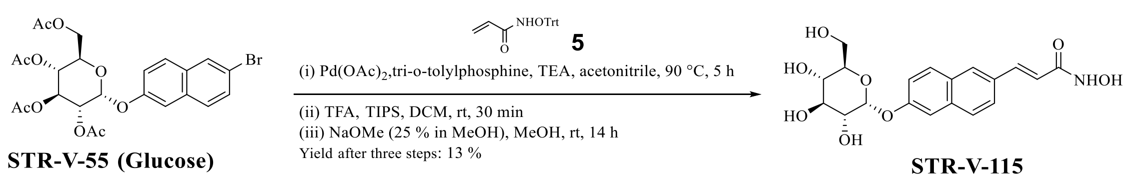

Glucose penta-acetate (compound **1**) was coupled with the 4-iodophenol (compound **2**) in the presence of Lewis acid Tin (IV) chloride in chloroform to yield **STR-V-49** (26). Sonogashira coupling of **STR-V-49** with TMS-acetylene followed by the removal of the TMS group by treatment TBAF furnished **STR-V-51**. Cu(I)-mediated Huisgen cyclization of **STR-V-51** with the TBDPS protected azido-hydroxamates (compound **3**), followed by the deprotection of the TBDPS group with cesium fluoride gave **STR-V-167**. The acetyl groups of **STR-V-167** was hydrolyzed by treatment with to sodium methoxide in methanol to furnish the target class I compounds **STR-V-53** (Scheme 1a).

To synthesize the class II glucosylated HDACi, 6-bromonaphthalen-2-ol (compound **4**) was coupled to the glucose penta-acetate (compound **1**) using Lewis acid boron trifluoride etherate in DCM under refluxing condition to furnish **STR-V-55**. Sonogashira coupling of the bromo naphthalene **STR-V-55** and TMS-acetylene using catalysts copper (I) iodide, bis(triphenylphosphine)palladium(II) dichloride, TPP in THF, followed by deprotection of the TMS moiety of the resulting intermediate using TBAF, resulted in **STR-V-111**. Subsequently, Cu(I)-mediated Huisgen cyclization of **STR-V-111** with the TBDPS protected azido-hydroxamates (compound **3**), followed by cesium fluoride deprotection and acetyl group removal furnished the target class II compound **STR-V-114** (Scheme 1b). Toward the class III glucosylated compound, Heck coupling of **STR-V-55** with *O*-trityl acrylamide (compound **5**) followed by the deprotection of the *O*-trityl group with TFA/TIPS in DCM and the acetyl group with sodium methoxide furnished the final product **STR-V-115** (Scheme 1c).

**Scheme 2.** Synthesis of the mannosylated HDACi **STR-V-176**, **STR-I-195**, **STR-V-177**, **STR-II-36** and **STR-V-105**.

2a

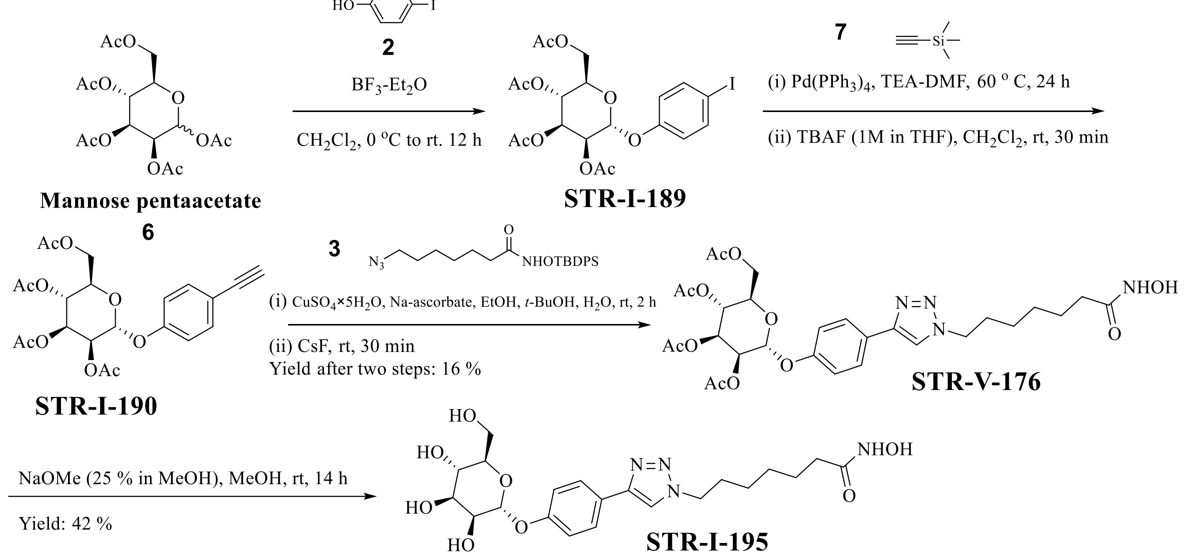


2b

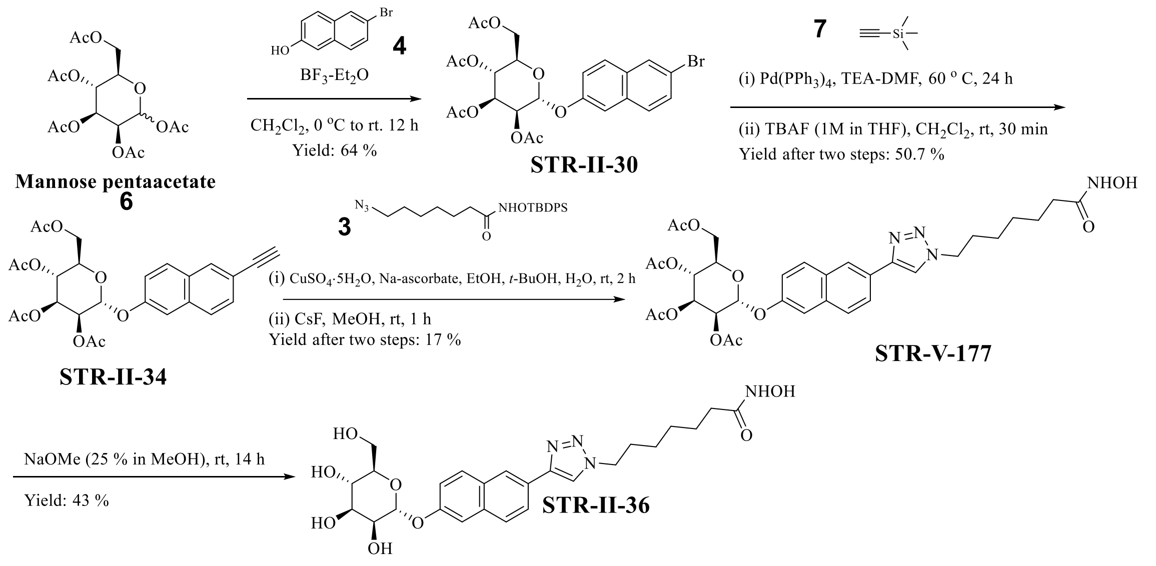

2c

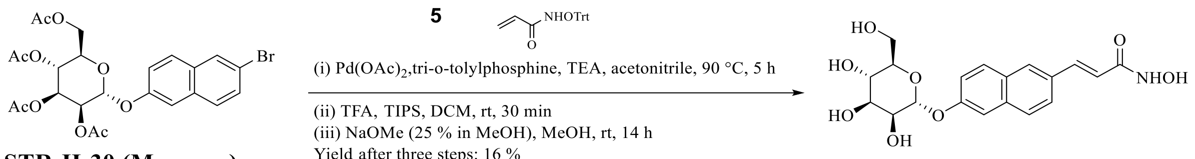


**STR-II-30 (Mannose)**

**STR-V-105**

To synthesize the mannose derivatives, mannose penta-acetate (compound **6**) was coupled with 4-iodophenol (compound **2**) using boron trifluoride etherate in DCM. The resulting **STR-I-189** was coupled with TMS-acetylene (compound **7**) using the same condition for the synthesis of **STR-V-51**. The resulting aceylated compound **STR-I-190** was reacted with TBDPS protect azido-hydroxamates (compound **3**) using the same condition as in the synthesis of **STR-V-167** to furnish **STR-V-176**. Subsequent hydrolysis of the acetyl groups yielded the target class I mannose compound **STR-I-195** (Scheme 2a).

Toward the mannosylated class II compound, the coupling of mannose penta-acetate (compound **6**) with 6-bromonaphthalen-2-ol (compound **4**) using boron trifluoride etherate, under the same condition for the synthesis of **STR-V-55**, furnished **STR-II-30**. The transformation of **STR-II-30** to the target mannosylated class II compound **STR-II-36** (Scheme 2b), via the intermediacy of **STR-V-177**, followed similar reaction steps used to synthesize analogous glucose compounds.

The class III mannosylated compounds was synthesized via Heck coupling of **STR-II-30** with *O*-trityl acrylamide (compound **5**) followed by the deprotection of the *O*-trityl group with TFA/TIPS in DCM and the acetyl group with sodium methoxide to furnish the target compound **STR-V-105** (Scheme 2c).

**Scheme 3.** Synthesis of the desosaminylated HDACi **STR-V-165**.

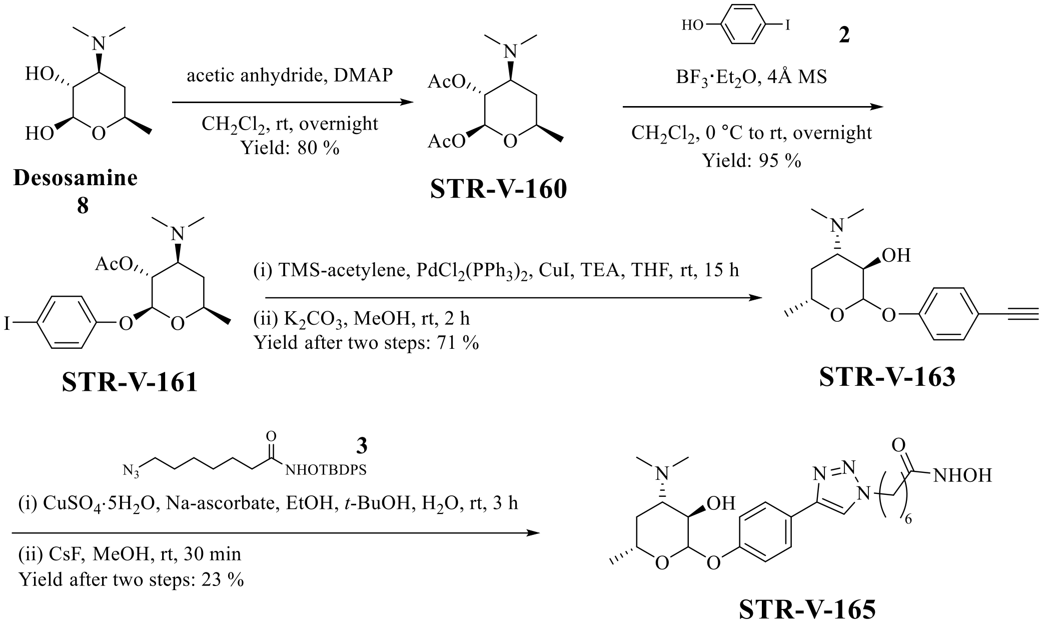


Lastly, the only desosaminylated compound disclosed herein was synthesize from diacetyl desosamine **STR-V-160**, which was obtained via acetylation of desosamine (compound **8**). The coupling of **STR-V-160** with 4-iodophenol (compound **2**), using the same method described for the synthesis of **STR-I-189**, resulted in **STR-V-161**, which was converted to **STR-V-163** using the same condition as in the synthesis of **STR-V-51**. Subsequently, **STR-V-163** was reacted with TBDPS protect azido-hydroxamate (compound **3**); the TBDPS and acetyl groups were deprotected to form **STR-V-165** (Scheme 3) in analogous manner to the synthesis of **STR-V-53**.

#### Materials and Methods

##### *Chemicals and reagents*

Anhydrous solvents and reagents were purchased from Sigma-Aldrich (St. Louis, MO, USA), Acros, VWR International (Radnor, PA, USA), Greenfield Chemicals, or Thermo Fisher

Scientific (Waltham, MA, USA) and were used without further purification. Analtech silica gel plates (60 F254) were utilized for analytical TLC, and Analtech preparative TLC plates (UV254, 2000 µm) were used for purification. Silica gel (200−400 mesh) was used in column chromatography. TLC plates were visualized using UV light, anisaldehyde, and iodine stains. High-performance liquid chromatography (HPLC) analyses were performed on an Agilent 1260 Infinity II instrument using a Luna® 5µm C-18 column 100 Å (100 mm × 4.6 mm). Eluting starts with solvent A (water, 0.1% formic acid) and solvent B (acetonitrile, 0.1% formic acid) at a gradient of 5% solvent B for the first 4 min and increase to 40% from 4 to 5 min, then continues with a gradient from 40 % to 80 % for another 20 min, and then constantly eluting with 80% solvent B for 5 min. The detection wavelength is at 280 nm and has a flow rate of 0.5 mL/min. Sample concentrations were 250 µM – 1mM, injecting 30 µL. All compounds have ≥ 95 % purity, as determined by HPLC. The solvents used LC-MS analyses were obtained from the following sources: acetonitrile (Optima, LCMS, Fischer Scientific, catalog No. A955-4); formic acid (Optima, LCMS, Fischer Scientific, catalog No. A117-50); water (Optima, LCMS, Fischer Scientific, catalog No. W6-4). Phloretin was obtained from VWR (Radnor, PA). NMR spectra were obtained on a Varian-Gemini 400 MHz and 700MHz magnetic resonance spectrometer. $^{1}$H NMR spectra were recorded in parts per million (ppm) relative to the residual peaks of $CHCl_3$ (7.24 ppm) in $CDCl_3$ or $CHD_2OD$ (4.78 ppm) in $CD_3OD$ or DMSO-d5 (2.49 ppm) in DMSO-d6. $^{13}$C spectra were recorded relative to the solvents peaks with complete hetero-decoupling. MestReNova (version 11.0) was used to process the original NMR “fid” files. High-resolution mass spectra were recorded at Georgia Institute of Technology’s Systems Mass Spectrometry Core facility.

*Cell lines and cell biology reagents*

Cell lines, including Hep-G2, SK-Hep-1, A549, MDA-MB-231, MCF-7, and VERO, were purchased from ATCC (Manassas, VA). Huh-7 and Kupffer cells were purchased from Sekisui XenoTech, LLC. Cells were cultured in the following media: Dulbecco’s Modified Eagle Medium (DMEM) (Corning, 10-017-CV); phenol red-free Minimum Essential Medium (MEM) (Corning, 17-305-CV) supplemented with 10% fetal bovine serum (FBS) (Corning, 35-010-CV); ATCC-formulated EMEM (ATCC® 30–2003™); DMEM containing 4 mM L-glutamine and 1 g/L glucose with 10% FBS (GIBCO Cat. #10099, Thermo Fisher Scientific); and OptiThaw Kupffer Cell Media (Sekisui XenoTech, LLC, Cat. #K8700). Electrophoresis supplies, TGX MIDI 4–20% gel (cat. # 5671093) and Turbo PDVF membrane (cat. # 1704273), were from Bio-Rad Laboratories, Inc (Hercules, CA, USA). Primary antibodies - Ac-Tubulin (sc-23950), Ac-H4 (sc-515319) were obtained from Santa Cruz Biotechnology (Dallas, TX, USA), anti-caspase 3 and anti-clv-caspase 3 were purchased from Cell Signaling Technology (Danvers, MA), anti-$p21^{WAF/CLIP}$ (ThermoFisher, Waltham, MA USA), anti-GAPDH (Sigma, Saint Louis, USA), secondary antibody (part. IR2173) was from ImmunoReagents (Raleigh, NC, USA). RIPA buffer (VWRVN653-100ML) was from VWR International (Radnor, PA, USA), phosphatase inhibitor (A32957) and protease inhibitor (A32955) were procured from Thermo Fisher Scientific (Waltham, MA, USA). The BCA protein assay kit (K813-2500) was purchased from BioVision, Inc (Milpitas, CA, USA). C57Bl/6 and Athymic Nude-Foxn1nu mice (5-7 weeks old; male and female) were purchased from Envigo RMS, Inc (Indianapolis, IN, USA).

*Cell culture*

Cell culture and viability assay protocols were described in our previous work. In brief, VERO and A549 cell lines were maintained in Dulbecco’s Modified Eagle Medium (DMEM) (Corning,

10-017-CV), supplemented with 10% fetal bovine serum (FBS) (Corning, 35-010-CV). Hep-G2 cells were cultured in phenol red-free Minimum Essential Medium (MEM) (Corning, 17-305-CV), supplemented with 10% fetal bovine serum (FBS). SK-Hep-1 in ATCC-formulated EMEM (ATCC® 30–2003™); Huh-7 in DMEM (containing 4 mM L-glutamine and 1 g/L glucose) with 10% FBS (GIBCO Cat. #10099, Thermo Fisher Scientific); and Kupffer cells in OptiThaw Kupffer Cell Media (Sekisui XenoTech, LLC, Cat. #K8700). Cells were seeded into a 96-well plate (2000 counts/100uL) for 24 h prior to treatment and then treated with various drug concentrations for 72 h. All drugs were dissolved in media with DMSO concentration maintained at 1%. The effect of compounds on cell viability was measured using the MTS assay (CellTiter 96 Aqueous One Solution and CellTiter 96 Non-Radioactive Cell Proliferation Assays, Promega, Madison, WI) as described by the manufacturer. $IC_{50}$s were determined using Prism GraphPad 8.

*Western blots analysis*

Western blot protocol was described in our previous works[24,25]. In brief, Hep-G2 cells were seeded into a 6-well plate at $1*10^6$/well in MEM for 24-48 h prior to treatment. Various concentrations of SAHA, **STR-V-53**, **STR-V-114,** and **STR-I-195** solutions in DMSO were added to the cell culture such that the final DMSO level was 0.1%. Cells were treated for 24 h, washed with cold PBS, and lysed with RIPA buffer (120 µL) (VWR, VWRVN653-100ML) containing phosphatase inhibitor (Fisher Thermo, A32957) and protease inhibitor (Fisher Thermo, A32955). The cell lysates were scraped, collected, and vortexed for 15 s, followed by sonication for 60 s. The lysate was centrifuged at 14000x rpm for 10 min, and the supernatants were collected. The total protein concentration was determined using a BCA protein assay kit (BioVision, K813-2500). Based on the results from the BSA assay, the lysates were diluted to make equal protein concentration, and 40 µg of each lysate was loaded to each well of the TGX

MIDI 4-20% gel (Biorad, cat. 5671093) and ran at 150 V for 70 min. The gel was then transferred onto the Turbo PDVF membrane (Biorad, 1704273), and after blocking with 5% BSA for 1-2 h, the membrane was incubated overnight with primary antibodies Ac-Tubulin (sc-23950), Ac-H4 (sc-515319) (Santa Cruz Biotechnology), caspase 3 and clv-Caspase 3 (Cell signaling Technology), anti-p21$^{WAF/CIP}$ (ThermoFisher), anti-GAPDH (Aldrich-Sigma). On the second day, the membrane was washed with TBST for 3x5 min; a secondary antibody (LiCOR) was added, and the membrane was incubated with agitation for 1 h. After washing with TBS-T 3x5 mins, bands were quantified using the Odyssey CLx Image system.

*GLUT-2 uptake experiment*

Hep-G2 or VERO cells were seeded (2500 to 4,000 counts/well) into 96-well plates, in which each well contains 100 µL media. The cells were cultured overnight in the respective media described in the cell culture section above. On day 2, the cells were treated with 25 µM Phloretin for 24 h. On day 3, 200x stock solutions of **STR-V-53** and SAHA were used to prepare 2x treatment solution by adding 1% stock solution in the media. Then, without removal of the media in 96-well plates, the same volumes of drug as the media were added directly to the wells with gentle pipetting to reach 1x drug solution with 25 µM Ph. After another 72 h, the media were removed, and new media (100 µL) added to the wells. MTS solution (20 µL) was added to each well. After 2.5-4 h incubation, the plates were read using a multiplate reader at 490 nm. The data were processed using GraphPad Prism 8.

*Flow cytometry*

Hep-G2 cells ($5*10^6$) were seeded in a 10 cm plate with MEM and incubated until 50% confluency before drug treatment.  Cells were treated, with DMSO (control) and DMSO solutions of SAHA (5 µM) and **STR-V-53** (15 µM) such that the final DMSO level was 0.1%, for another 48 h. Cells were washed with cold 1x PBS solution twice and trypsinized. Subsequently, cells were collected using 5 mL 1x PBS buffer and fixed overnight at -20ºC using 70% ethanol. Cells were centrifuged in 1x PBS on day 2 at 4 ºC with a spin speed of 3,000g, re-suspended in 1x PBS, and centrifuged again under the same condition. The supernatant was removed, and the cell pellets were re-suspended with 500 µL of 200 ug/mL RNase for 30 min. Then cells were treated with another 500 µL of 100 ug/mL PI staining at room temperature for 30 min. The cell cycle was analyzed with BD FACS Aria Illu analyzer, and the data were processed using FlowJo.

*Plasma stability*

The stability of **STR-V-53** in murine and human plasmas was determined through a contract agreement with Cyprotex, an Evotec Company (https://www.cyprotex.com)

*Maximum tolerated dose determination*

C57BL/6 mice, 6–8 weeks old, 3 males and 3 females per treatment group, were injected intraperitoneally with **STR-V-53** at concentrations of 25, 50, and 100 mg/kg once daily for 7 days in Kolliphor/DMA/water vehicle. We measured body weight and monitored daily food consumption, overall health, and mortality. On the eighth day, mice were sacrificed by cervical dislocation, and organs were harvested and frozen at −80 ◦C.

*Tissue distribution studies*

C57Bl/6 mice, 6–8 weeks old, 3 males and 3 females per treatment group, were injected intraperitoneally with **STR-V-53 at** 50 mg/kg in Kolliphor/DMA/water vehicle. Eight hours after the injection, organs were harvested and frozen at −80ºC for subsequent quantification by LC-MS analyses[18].

*Immunofluorescence (IF)*

$CD4^+$ or $CD8^+$ T-cells were detected by IF using an anti-CD4 antibody (rat monoclonal, clone RM4-5, BD Pharmingen) and an anti-CD8 antibody (rabbit monoclonal, clone D4W2Z, Cell Signaling Technology), respectively, for immunostaining. To detect regulatory T cells (Tregs), an anti-Foxp3 antibody (rabbit monoclonal, clone D6O8R, Cell Signaling Technology) was used to identify $Foxp3^+CD4^+$ cells. For IF analyses of endothelial, we used antibodies against CD31 (Armenian Hamster monoclonal antibody, clone 2H8, Millipore). Positive cells were counted from five random fields for each HCC tissue section. All images were obtained using a confocal microscope (FLUOVIEW FV100, OLYMPUS, Center Valley, PA). These image data were analyzed using ImageJ (US NIH) and Photoshop (Adobe Systems Inc., San Jose, CA) software.

*In vivo efficacy studies*

The *in vivo* efficacy of **STR-V-53**, alone or in combination with sorafenib, was evaluated in a murine orthotopic xenograft model of Hep-G2-Red-Flu human HCC, as previously described[18].

Next, we evaluated the *in vivo* efficacy of **STR-V-53** alone or combined with an anti-PD1 antibody in a p53-null murine HCC model. RIL-175 murine HCC cells in Matrigel were injected into the subcapsular region of the liver parenchyma in C57Bl/6 mice, and tumor growth was monitored by high-frequency ultrasonography. When tumors reached ~5 mm in diameter, mice were randomly assigned to one of the four treatment groups (n=12 mice): DMA/ CRH/Water (10%/20%/70%) (control), **STR-V-53** (25mg/kg daily by i.p. for 3 weeks), anti-PD1 antibody (10mg/kg thrice a week for 3 weeks), or combination of **STR-V-53** and anti-PD1 antibody. Anti-PD1 antibody treatment was administered i.p. for a total of 9 injections. Tumor growth and body weight were measured every three days by ultrasound imaging, and tumor size was calculated as

$0.5 \times \text{length} \times \text{width}^2$. Per protocol, moribund status was used as the endpoint, and moribundity was defined as symptoms of prolonged distress, more than 15% of weight loss compared with the starting date, body condition score of more than 2, and tumor size of more than 15 mm in diameter. Data were processed using GraphPad Prism 8, and survival analysis was performed using JMP Pro 13.

RNA-seq Protocol

HepG2 cells were seeded in a 6-well plate at a density of 2 x $10^5$ cells per well in complete MEM media and allowed to grow for 48 hours. The cells were then treated with 10 and 20 µM **STR-V-53** solutions dissolved in 100% DMSO and diluted with media, ensuring that the final DMSO concentration was 1%. After a 24-hour incubation period, the cells were trypsinized, followed by centrifugation for 5 min at 500 RCF. The resulting cell pellets were resuspended in cold 1X PBS and centrifuged again for 5 min at 500 RCF, with this washing step repeated twice. Next, using RNeasy Plus Mini Kit and Qiagen QIAcube Connect RNA Purification system, RNA samples were extracted from the cells and the concentrations determined using a DeNovix spectrophotometer. Subsequently, quality control of the RNA samples was carried out using Agilent RNA 6000 Nano Kit which is designed for use with Agilent 2100 Bioanalyzer instrument only. High quality RNA samples having RNA integrity numbers of 7 or above were used for the next step. RNA-seq libraries were prepared based on NEBNext Ultra II Directional RNA Library Prep Kit for Illumina using NEBNext Poly(A) mRNA Magnetic Isolation Module (NEB #E7490). RNA sequencing was conducted on Illumina NovaSeq X Plus PE150 bp reads at the Molecular Evolution Core, Georgia Institute of Technology. The quality of sequencing was measured using Phred quality score (Q score) and it was established that more than 99% of the sequencing reads attained over 99.99% base call accuracy (base call error rate of $1\text{x}10^{-39}$).

FASTQ files of samples were analyzed through the RNA-seq pipeline provided by the Gryder lab[1]. Reads were aligned to hg38 using STAR version 2.5.3a, and gene expression was calculated as Transcripts per Million mapped reads (TPM) using RSEM version 1.3.3 and filtered for coding genes. Differentially expressed genes lists were generated for each sample by calculating log2 fold change of the averaged sample TPM versus the averaged DMSO-treated control cells TPM and filtering for genes with |log2 fold change| > -1 and false discovery rate < 0.25. Hallmark[2] and gene ontology biological processes (GOBP)[3] gene set enrichment analysis (GSEA) was performed on the differentially expressed gene lists using the GSEA software (parameters: Run GSEAPreranked, Collapse/Remap- No_Collapse)[4]. GSEA results were interpreted and visualized using scripts provided by the Gryder lab[1].Volcano plots were generated using DESeq2v1.41.1 and visualized with the ggplot2 R packages[5,6].

#### Synthesis Protocol

*Published/Reported Agents*

We have described the synthesis and biological characterization of the control compound **STR-V-48** (N-hydroxy-7-(4-(6-methoxynaphthalen-2-yl)-1H-1,2,3-triazol-1-yl)heptanamide) in a previous publication[27]. Intermediates **STR-V-51** (Compound CID: 102084446) and **STR-I-190** have been described in the patents (WO2014100158A1). **STR-V-49** was reported in a patent WO2014100158A1. n-(Trityloxy)acrylamide (**5**) is a commercially available chemical (CAS: 79-06-1).

(2R,3R,4S,5R,6R)-2-(Acetoxymethyl)-6-(4-(1-(7-(((tert-butyldiphenylsilyl)oxy)amino)-7-oxoheptyl)-1H-1,2,3-triazol-4-yl)phenoxy)tetrahydro-2H-pyran-3,4,5-triyl triacetate **(STR-V-**

**167).** **STR-V-51** (1.00 g, 2.20 mmol) and **3** (0.88 g, 2.07 mmol) were dissolved in a 1:1 mixture of ethanol and tert-butanol (12 mL). Copper (II) sulfate (99 mg, 0.40 mmol) and sodium ascorbate (313 mg. 1.58 mmol) were added to the solution together with 0.5 mL water. The was stirred at rt for 3 h during which TLC revealed a complete consumption of the starting materials. The crude was partitioned between DCM (30 mL) and water (50 mL) and two layers separated. The organic layer was washed with water (30 mL), dried over $Na_2SO_4$ and solvent evaporated *in vacuo*. The residue was purified using column chromatography, eluting with EtOAc: hexane 8:2 to furnish the TDBPS protected intermediate (1.10 g, 84 %). Then, the intermediate (1.10g, 1.27mmol) was dissolved into methanol and treated with cesium fluoride (330 mg, 2.20 mmol) at rt for 15 min. The final product was yielded as a brownish-white solid (Yield: 730 mg, 66.1%). $^{1}$H NMR (700 MHz, $CDCl_3$) δ 7.75 (d, *J* = 8.0 Hz, 2H), 7.68 (s, 1H), 7.13 (d, *J* = 7.8 Hz, 2H), 5.75 (d, *J* = 3.5 Hz, 1H), 5.69 (t, *J* = 9.8 Hz, 1H), 5.15 (t, *J* = 9.8 Hz, 1H), 5.04 (dd, *J* = 10.3, 3.5 Hz, 1H), 4.38 (s, 1H), 4.24 (d, *J* = 4.6 Hz, 1H), 4.11 (ddd, *J* = 10.6, 4.7, 2.1 Hz, 1H), 4.04 (dd, *J* = 12.4, 2.2 Hz, 1H), 2.14 (s, 2H), 2.07 – 1.98 (m, 12H), 1.92 (s, 2H), 1.65 (s, 2H), 1.39 – 1.21 (m, 6H). $^{13}$C NMR (176 MHz, $CD_3OD$) δ 171.1, 170.5, 170.0, 156.0, 147.1, 127.0, 125.4, 119.9, 117.1, 94.2, 70.4, 70.1, 68.3, 68.0, 61.7, 50.3, 32.5, 29.9, 28.1, 25.8, 25.1, 20.3. HRMS (ESI) m/z Calcd. for $C_{29}H_{39}O_{12}N_4$ [M+H$^+$]: 635.2559, found 635.2540. Retention time: 16.5 min.

N-Hydroxy-7-(4-(4-(((2R,3R,4S,5S,6R)-3,4,5-trihydroxy-6-(hydroxymethyl)tetrahydro-2H-pyran-2-yl)oxy)phenyl)-1H-1,2,3-triazol-1-yl)heptanamide **(STR-V-53).** A mixture of **STR-V-167** (730 mg, 1.15 mmol) and sodium methoxide (0.5 mL, 2.3 mmol) was stirred in MeOH (2 mL) at rt for 14 h during which TLC revealed a complete consumption of the starting material. The reaction was treated with Ambolite IR120 Plus resin to adjust the pH to 1. [**Important**

**Note:** some precipitates crashed out when methoxide was added if the reaction was in larger scale (>500mg), however, the addition of Resin rapidly eliminated the precipitation.] The resin was filtered, and the filtrate was collected and evaporated to dryness *in vacuo* to furnish the crude **STR-V-53**. To purify, the crude **STR-V-53** was re-dissolved in water (15 mL) and washed with 15% MeOH in DCM (30 mL). The aqueous layer was collected and evaporated to dryness. To completely dry the product, it was dispersed in acetonitrile (1 mL) and water (0.1 mL) and lyophilized to furnish **STR-V-53** as white solid was (Yield: 460 mg, 85.2 %). $^{1}$H NMR (700 MHz, DMSO-*d6*) δ 10.33 (s, 1H), 8.48 (s, 1H), 7.76 (d, *J* = 8.6 Hz, 2H), 7.16 (d, *J* = 8.7 Hz, 2H), 5.76 (s, 0H), 5.42 (d, *J* = 3.5 Hz, 1H), 4.36 (t, *J* = 7.1 Hz, 2H), 3.63 (t, *J* = 9.2 Hz, 1H), 3.57 (m, 3.59-3.55, 4.2 Hz, 1H), 3.47 (m, 3.49-3.42, 2H), 3.38 (dd, *J* = 9.7, 3.5 Hz, 1H), 3.20 (t, *J* = 9.2 Hz, 1H), 1.93 (t, *J* = 7.4 Hz, 2H), 1.85 (p, *J* = 7.1 Hz, 2H), 1.48 (p, *J* = 7.3 Hz, 2H), 1.27 (m, 1.32-1.22, 5H). $^{13}$C NMR (176 MHz, $CD_3OD$) δ 171.5, 157.3, 147.2, 126.5, 124.5, 120.3, 117.1, 97.9, 73.5, 71.9, 70.1, 61.0, 50.0, 48.5, 32.2, 29.7, 28.0, 25.7, 25.1. HRMS (ESI) m/z Calcd. for $C_{21}H_{31}O_8N_4$ [M+H$^+$]: 467.2136, found 467.2123. Retention time: 2.1 min.

(2R,3R,4S,5R,6R)-2-(Acetoxymethyl)-6-((6-bromonaphthalen-2-yl)oxy)tetrahydro-2H-pyran-3,4,5-triyl triacetate (**STR-V-55**). Beta-D-glucose pentaacetate (3.31 g, 8.49 mmol) and 6-bromo-2-naphthol (2.34 g, 10.18 mmol) were dissolved in DCM (25 mL) at 0 ºC in ice bath. Boron trifluoride etherate (1.6 mL, 12.74 mmol) was added drop-wise, the reaction was removed from ice and stir at rt for 30 min. Subsequently, the reaction was heated under reflux for 48 h, partitioned between water (30 mL) and DCM (30 mL) and the two layers separated. The organic layer was dried over $Na_2SO_4$ and evaporated to dryness. The crude was purified via column chromatography (EtOAc: hexane 3:7), to furnish **STR-V-55** (Yield: 523 mg, 80%). $^{1}$H NMR

(400 MHz, $CDCl_3$) δ 7.95 (d, *J* = 2.0 Hz, 1H), 7.71 (d, *J* = 9.0 Hz, 1H), 7.61 (d, *J* = 8.7 Hz, 1H), 7.53 (dd, *J* = 8.8, 2.0 Hz, 1H), 7.43 (d, *J* = 2.5 Hz, 1H), 7.33 – 7.25 (m, 2H), 5.88 (d, *J* = 3.6 Hz, 1H), 5.75 (t, *J* = 10.3, 1H), 5.19 (t, *J* = 9.4 Hz, 1H), 5.10 (dd, *J* = 10.3, 3.6 Hz, 1H), 4.30-4.23 (m, 1H), 4.13 (ddd, *J* = 10.2, 4.6, 2.1 Hz, 1H), 4.05 (d, *J* = 12.3 Hz, 1H), 2.13 – 2.00 (m, 9H), 1.99 (s, 3H).

(2R,3R,4S,5R,6R)-2-(Acetoxymethyl)-6-((6-ethynylnaphthalen-2-yl)oxy)tetrahydro-2H-pyran-3,4,5-triyl triacetate (**STR-V-111**). **STR-V-55** (354 mg, 0.63 mmol), TMS-acetylene (0.13 mL, 0.95 mmol), Bis(triphenylphosphine)palladium(II) dichloride (9 mg, 0.01 mmol), copper iodide (3.4 mg, 0.02 mmol) and TPP (2.5 mg, 0.01 mmol) were dissolved into THF (3 mL) and TEA addition (0.17 mL). The reaction was kept stirring at rt for 72 h. The solution was filtered through celite and evaporated off. The crude product was purified via column chromatoghraphy, eluting with EtOAc: hexane 1:2 to furnish the silyl intermediate (Yield: 250 mg, 79.3%). The silyl intermediate (45 mg, 0.08 mmol) was dissolved into DCM and TBAF (0.08 mL, 0.08 mmol) was added to the solution. The reaction was stirred at rt for 30 min, solvent was evaporated off and the residue was purified via column chromatography (EtOAc: hexane 3:7) to furnish **STR-V-111** as brownish-white solid (Yield: 30 mg, 75 %). $^1$H NMR (700 MHz, $CDCl_3$) δ 7.94 (d, *J* = 13.6 Hz, 1H), 7.73 (d, *J* = 9.0 Hz, 1H), 7.67 (dd, *J* = 15.7, 8.7 Hz, 1H), 7.59 (d, *J* = 8.7 Hz, 1H), 7.50 (dd, *J* = 13.6, 8.6 Hz, 1H), 7.41 (t, *J* = 3.3 Hz, 1H), 7.30 – 7.25 (m, 1H), 5.86 (dd, *J* = 10.2, 3.5 Hz, 1H), 5.72 (td, *J* = 9.9, 2.4 Hz, 1H), 5.16 (t, *J* = 9.9 Hz, 1H), 5.08 (dt, *J* = 10.4, 3.4 Hz, 1H), 4.26 – 4.21 (m, 1H), 4.11 (dd, *J* = 10.6, 4.4 Hz, 1H), 4.03 (d, *J* = 12.5 Hz, 1H), 3.11 (s, 1H), 2.06 – 2.01 (m, 9H), 1.96 (s, 3H).

N-Hydroxy-7-(4-(6-(((2R,3R,4S,5S,6R)-3,4,5-trihydroxy-6-(hydroxymethyl)tetrahydro-2H-pyran-2-yl)oxy)naphthalen-2-yl)-1H-1,2,3-triazol-1-yl)heptanamide (**STR-V-114**). A mixture of **STR-V-111** (29 mg, 0.06 mmol), **3** (30 mg, 0.07 mmol), copper (II) sulfate (1.5 mg, 0.01 mmol), and sodium ascorbate (4.75 mg, 0.02 mmol) in ethanol (1.5 mL) and tert-butanol (1.5mL) was reacted, work-up and the crude purified as described for **STR-V-167** to furnish the intermediate silyl product (53 mg, 96%). The intermediate silyl product was treated with CsF (17 mg, 0.11 mmol) in MeOH (2 mL) at rt for 30 min. Solvent was evaporated off and the residue was partitioned between ethyl acetate (40 mL) and water (30 mL), the two layers were separated, the organic layer was dried over $Na_2SO_4$, and solvent was evaporated off. The residue was stirred in methanol (2 mL) and NaOMe (25% in methanol) (0.08 mL, 0.34 mmol) at rt for 14 h and the crude was purified as described for **STR-V-53** to furnish **STR-V-114** as white solid (Yield: 9 mg, 31 %). $^1$H NMR (700 MHz, DMSO-*d6*) δ 10.33 (s, 1H), 8.14 (s, 1H), 7.95-7.89 (m, 1H), 7.89-7.76 (m, 3H), 7.58 (d, *J* = 9.6 Hz, 2H), 7.35-7.29 (m, 2H), 5.56 (t, *J* = 2.6 Hz, 1H), 3.67 (t, *J* = 9.5 Hz, 1H), 3.56 (d, *J* = 10.5 Hz, 1H), 3.47 (d, *J* = 9.1 Hz, 2H), 3.42 (dt, *J* = 9.8, 2.7 Hz, 1H), 3.31 (t, *J* = 7.0 Hz, 2H), 3.21 (t, *J* = 9.2 Hz, 1H), 3.17 (d, *J* = 1.9 Hz, 0H), 1.93 (t, *J* = 7.4 Hz, 2H), 1.50 (dp, *J* = 22.8, 7.4 Hz, 4H), 1.28 (dq, *J* = 32.1, 7.7 Hz, 6H). HRMS (ESI) m/z Calcd. for $C_{25}$ $H_{33}$ $O_8$ $N_4$ [M+H$^+$]: 517.2293, found 517.2272.

(E)-N-Hydroxy-3-(4-(6-(((2R,3R,4S,5S,6R)-3,4,5-trihydroxy-6-(hydroxymethyl)tetrahydro-2H-pyran-2-yl)oxy)naphthalen-2-yl)-1H-1,2,3-triazol-1-yl)acrylamide (**STR-V-115**). **STR-V-55** (156 mg, 0.28 mmol), compound **5** (186 mg, 0.56 mmol), Pd(OAc) (9.4 mg, 0.04 mmol), tri(o-tolyl)phosphine (26.3 mg, 0.084 mmol) dissolved in acetonitrile (3 mL) and the mixture was flushed with argon for about 5 min. TEA (0.1 mL. 0.74 mmol) was added and the mixture was heated at 90$^o$C for 5 h. The mixture was filtered over a celite bed and the filtrate was evaporated *in vacuo*. The crude product was purified using column chromatography eluting with EtOAc:hexanes 6:4 to furnish the *O*-trityl protected intermediate (Yield: 57 mg, 25%). The trityl protected intermediate (55 mg, 0.07 mmol) was dissolved into DCM (1.5 mL), TIPS (0.1 mL) and TFA (0.5 mL) and the reaction was stirred at rt for 30 min. Solvent was evaporated off and the residue was stirred in MeOH (2 mL) and NaOMe (0.1 mL) at rt for 14 h and purified as described for the synthesis of **STR-V-53** to furnish **STR-V-115** as brownish-white solid (Yield: 14 mg, 52%). $^1$H NMR (700 MHz, DMSO-*d6*) δ 10.74 (s, 1H), 7.99 (s, 1H), 7.83 (dd, *J* = 45.6, 8.8 Hz, 2H), 7.64 (d, *J* = 8.6 Hz, 1H), 7.55 (d, *J* = 17.3 Hz, 2H), 7.29 (dd, *J* = 8.9, 2.7 Hz, 1H), 6.51 (d, *J* = 15.7 Hz, 1H), 5.73 (d, *J* = 2.7 Hz, 1H), 5.55 (d, *J* = 2.9 Hz, 1H), 3.65 (td, *J* = 9.3, 2.8 Hz, 1H), 3.56 – 3.52 (m, 1H), 3.46 (d, *J* = 9.5 Hz, 3H), 3.42 – 3.37 (m, 1H), 3.21 – 3.17 (m, 1H), 3.14 (d, *J* = 2.8 Hz, 1H). $^{13}$C NMR (176 MHz, $CD_3OD$) δ 156.0, 140.5, 135.3, 130.7, 129.7, 129.6, 128.8, 127.6, 123.5, 119.6, 110.9, 97.9, 74.2, 73.6, 73.2, 71.9, 70.1, 61.0, 48.5. HRMS (ESI) m/z Calcd. for $C_{19} H_{22} O_8$ N [M+H$^+$]: 392.1340, found 392.1340.

(2R,3R,4S,5R,6S)-2-(Acetoxymethyl)-6-(4-(1-(7-(hydroxyamino)-7-oxoheptyl)-1H-1,2,3-triazol-4-yl)phenoxy)tetrahydro-2H-pyran-3,4,5-trityl triacetate (**STR-V-176**). The procedure used to accomplish the synthesis of the title compound was as described for the synthesis of **STR-V-167**. Briefly, the reaction of **STR-I-190** (67 mg, 0.15 mmol), **3** (67 mg, 0.16 mmol), copper (II) sulfate (3.7 mg, 0.015 mmol) and sodium ascorbate (12 mg, 0.06 mmol) in $H_2O$ (0.2 mL), tert-butanol (1.5 mL) and ethanol (1.5mL), after column chromatography eluting with EtOAc: hexane 2:8, furnished the TBDPS protected intermediate (91 mg, 66.7% ). The intermediated was treated with cesium fluoride (32 mg, 0.20 mmol) in MeOH (2 mL) at rt for 30 min and the reaction was worked-up as described for **STR-V-53**. The crude was purified using preparative TLC, eluting with DCM:MeOH 9:1, to give **STR-V-176** as white solid (Yield: 15 mg, 23.6%). $^{1}$H NMR (700 MHz, $CDCl_3$) δ 7.81 – 7.68 (m, 3H), 7.12 (d, *J* = 8.1 Hz, 2H), 5.56 – 5.51 (m, 2H), 5.43 (dd, *J* = 3.6, 1.8 Hz, 1H), 5.35 (t, *J* = 10.1 Hz, 1H), 4.34 (t, *J* = 6.6 Hz, 2H), 4.25 (dd, *J* = 12.2, 5.2 Hz, 1H), 4.10 – 4.02 (m, 2H), 2.18 (s, 3H), 2.14 – 2.08 (m, 2H), 2.04 – 1.98 (m, 9H), 1.90 – 1.86 (m, 2H), 1.60 (s, 2H), 1.31 (d, *J* = 26.4 Hz, 6H). $^{13}$C NMR (176 MHz, $CD_3OD$) δ 171.3, 171.1, 170.4, 170.3, 170.2, 155.5, 147.1, 127.0, 125.3, 120.1, 116.9, 95.7, 77.7, 77.5, 77.3, 69.2, 69.2, 69.1, 65.8, 62.1, 50.2, 32.4, 29.8, 28.1, 25.8, 25.1, 20.3, 20.2. HRMS (ESI) m/z Calcd. for $C_{29}H_{39}O_{12}N_4$ [M+H$^+$]: 635.2559, found 635.2552.

N-Hydroxy-7-(4-(4-(((2S,3R,4S,5S,6R)-3,4,5-trihydroxy-6-(hydroxymethyl)tetrahydro-2H-pyran-2-yl)oxy)phenyl)-1H-1,2,3-triazol-1-yl)heptanamide (**STR-I-195**). Following the procedure used for the synthesis of **STR-V-53**, **STR-V-176** (13 mg, 0.02 mmol) was treated with NaOMe (0.1 mL, 0.50 mmol) in MeOH (1 mL) to furnish **STR-I-195** as pale-white foam (Yield: 4.2 mg, 42 %). $^{1}$H NMR (400 MHz, DMSO-*d6*) δ 10.31 (s, 4H), 8.47 (s, 1H), 7.73 (d, *J* = 8.7 Hz, 2H), 7.13 (d, *J* = 8.6 Hz, 2H), 5.38 (d, *J* = 1.8 Hz, 1H), 4.34 (t, *J* = 7.1 Hz, 2H), 3.82 (s, 1H), 3.71 – 3.53 (m, 2H), 3.53 – 3.20 (m, 4H), 1.94 – 1.76 (m, 4H), 1.46 (d, *J* = 8.0 Hz, 2H), 1.25 (s, 6H). $^{13}$C NMR (100 MHz, $CD_3OD$) δ 171.5, 156.6, 147.1, 126.6, 124.5, 120.3, 116.8, 98.7, 74.1, 71.0, 70.5, 66.9, 65.5, 61.3, 32.2, 29.7, 28.0, 25.7, 25.1. HRMS (ESI) m/z Calcd. for $C_{21}H_{31}O_8N_4$ [M+H$^+$]: 467.2136, found 467.2136.

(2R,3R,4S,5R,6S)-2-(Acetoxymethyl)-6-((6-bromonaphthalen-2-yl)oxy)tetrahydro-2H-pyran-3,4,5-triyl triacetate (**STR-II-30**). The reaction of mannose pentaacetate (1.48 g, 3.8 mmol), 6-bromo-2-naphthol (3.5 g, 15.21 mmol), and boron trifluoride etherate (0.72 mL, 5.70 mmol) in DCM (20 mL) at 0$^o$C to rt overnight, followed by work-up as described for **STR-V-55** and purification using column chromatography, eluting with EtOAc:hexane 35:65, furnished **STR-II-30** (Yield: 1.34, 64 %). $^1$H NMR (400 MHz, $CDCl_3$) δ 7.94 (d, *J* = 2.3 Hz, 1H), 7.77 – 7.66 (m, 1H), 7.63 – 7.38 (m, 3H), 7.31 – 7.23 (m, 1H), 5.67 (d, *J* = 1.9 Hz, 1H), 5.60 (dd, *J* = 10.0, 3.5 Hz, 1H), 5.49 (dd, *J* = 3.6, 1.9 Hz, 1H), 5.39 (t, *J* = 10.0 Hz, 1H), 4.29 (dd, *J* = 12.0, 5.2 Hz, 1H), 4.16 – 4.03 (m, 2H), 2.22 (s, 3H), 2.05 (s 3H), 1.95 (s, 3H). HRMS (ESI) m/z Calcd. for C24 H25 O10 Br Na [M+Na$^+$]: 707.2535, found 707.2515.

(2R,3R,4S,5R,6S)-2-(Acetoxymethyl)-6-((6-ethynylnaphthalen-2-yl)oxy)tetrahydro-2H-pyran-3,4,5-triyl triacetate (**STR-II-34**). **STR-II-30** (301 mg, 0.54 mmol), Tetrakis(triphenyl-phosphine)palladium(0) (31 mg, 0.03 mmol), TMS-acetylene (0.14 mL, 1.0 mmol) were dissolved in TEA (15 mL) and DMF (1 mL). The mixture was flushed with argon and heated under argon atmosphere at 60ºC for 12 h. The mixture was allowed to cool down, and filtered through a celite bed to remove palladium. The filtrate was evaporated to dryness and purified through column, eluting with EtOAc:hexane 3:7 to furnish the silyl intermediate as yellowish-white solid (Yield: 239 mg, 84 %). The silyl intermediate (233 mg, 0.43 mmol) was dissolved into THF (2 mL), and solution was kept stirring in ice bath. 1M TBAF (solution in methanol, 0.43 mL, 0.43 mmol) and AcOH (0.04 mL) were added, the reaction was allowed to warm to rt and stirring continued at rt for about 24 h. Solvent was evaporated off and the residue was purified via column chromatography, eluting with EtOAc:hexane 4:6 to furnish **STR-II-34**

(Yield: 129 mg, 60.4 %). $^{1}$H NMR (400 MHz, $CDCl_3$) δ 7.96 (t, *J* = 1.1 Hz, 1H), 7.74 (dd, *J* = 8.9, 0.7 Hz, 1H), 7.70 – 7.63 (m, 1H), 7.50 (dd, *J* = 8.5, 1.6 Hz, 1H), 7.45 (d, *J* = 2.5 Hz, 1H), 7.30 – 7.23 (m, 1H), 5.68 (d, *J* = 1.9 Hz, 1H), 5.60 (dd, *J* = 10.0, 3.5 Hz, 1H), 5.50 (dd, *J* = 3.5, 1.8 Hz, 1H), 5.39 (t, *J* = 10.0 Hz, 1H), 4.29 (dd, *J* = 12.1, 5.3 Hz, 1H), 4.15 – 4.03 (m, 2H), 3.13 (s, 1H), 2.22 (s, 3H), 2.05 (d, *J* = 1.7 Hz, 6H), 1.94 (s, 3H). HRMS (ESI) m/z Calcd. for $C_{26}$ $H_{26}$ $O_{10}$ Na [M+Na$^{+}$]: 707.2535, found 707.2515.

(2R,3R,4S,5R,6S)-2-(Acetoxymethyl)-6-((6-(1-(7-(hydroxyamino)-7-oxoheptyl)-1H-1,2,3-triazol-4-yl)naphthalen-2-yl)oxy)tetrahydro-2H-pyran-3,4,5-triyl triacetate (**STR-V-177**). **STR-II-34** (65 mg, 0.13 mmol), compound **3** (61 mg, 0.14 mmol), copper (II) sulfate (3.2 mg, 0.01 mmol) and sodium ascorbate (10.3 mg, 0.05 mmol) were stirred in $H_2O$ (0.2 mL), tert-butanol (1.5 mL) and ethanol (1.5 mL) for 3h at rt. After work-up, the crude was reacted with cesium fluoride (32 mg, 0.20 mmol) in MeOH (2 mL) at rt for 30 min and the reaction was worked-up as described for **STR-V-53**. The crude was purified using preparative TLC, eluting with 7 % MeOH in DCM to give **STR-V-177** as white solid (Yield: 15 mg, 24 %). $^{1}$H NMR (700 MHz, $CDCl_3$) δ 8.24 (s, 1H), 7.85 (d, *J* = 9.7 Hz, 2H), 7.80 (d, *J* = 8.8 Hz, 1H), 7.75 (d, *J* = 8.2 Hz, 1H), 7.45 (d, *J* = 2.4 Hz, 1H), 7.23 (s, 3H), 5.58 (dd, *J* = 10.0, 3.6 Hz, 1H), 5.47 (dd, *J* = 3.6, 1.8 Hz, 1H), 5.37 (t, *J* = 10.1 Hz, 1H), 4.38 (s, 2H), 4.27 (dd, *J* = 12.3, 5.5 Hz, 1H), 4.11 (ddd, *J* = 10.2, 5.4, 2.3 Hz, 1H), 4.05 (dd, *J* = 12.3, 2.3 Hz, 1H), 3.21 (t, *J* = 6.8 Hz, 4H), 2.19 (s, 3H), 2.12 (s, 3H), 2.02 (s, 3H), 1.92 (s, 3H), 1.60 (s, 4H), 1.54 (p, *J* = 7.1 Hz, 4H), 1.31 (q, *J* = 14.1, 12.6 Hz, 4H), 1.21 (s, 3H). $^{13}$C NMR (101 MHz, $CD_3OD$) δ 171.5, 171.0, 170.3, 170.3, 170.2, 153.5, 147.5, 134.0, 130.1, 129.9, 127.7, 126.6, 124.3, 124.0, 120.9, 119.0, 110.9, 95.7, 69.3, 69.2, 65.8,

51.1, 47.9, 47.8, 47.6, 29.8, 28.4, 28.4, 26.1, 25.3, 19.8, 19.7, 19.7 .HRMS (ESI) m/z Calcd. for $C_{33} H_{40} O_{12} N_4 Na$ [M+Na$^+$]: 707.2535, found 707.2515.

N-Hydroxy-7-(4-(6-(((2S,3R,4S,5S,6R)-3,4,5-trihydroxy-6-(hydroxymethyl)tetrahydro-2H-pyran-2-yl)oxy)naphthalen-2-yl)-1H-1,2,3-triazol-1-yl)heptanamide (**STR-II-36**). Following the procedure used for the synthesis of **STR-V-53**, **STR-V-177** (219 mg, 0.27 mmol) was treated with NaOMe (0.5 mL, 2.19 mmol) in MeOH (2 mL) to furnish **STR-II-36** as whitish foam (90 mg, 43 %). $^1$H NMR (400 MHz, DMSO-*d6*) δ 10.31 (s, 1H), 8.64 (s, 1H), 8.31 (s, 2H), 7.98 – 7.80 (m, 4H), 7.56 (d, *J* = 2.4 Hz, 1H), 7.27 (dd, *J* = 8.9, 2.5 Hz, 1H), 5.53 (s, 1H), 4.39 (t, *J* = 6.9 Hz, 2H), 3.87 (dd, *J* = 3.5, 1.8 Hz, 1H), 3.72 (dd, *J* = 9.1, 3.4 Hz, 1H), 3.59 (d, *J* = 10.8 Hz, 1H), 3.55 – 3.38 (m, 4H), 1.95 – 1.84 (m, 4H), 1.48 (d, *J* = 7.3 Hz, 2H), 1.40 – 1.07 (m, 6H). $^{13}$C NMR (101 MHz, $CD_3OD$) δ 154.6, 134.3, 134.2, 129.7, 129.4, 127.6, 126.1, 123.8, 123.7, 120.9, 119.2, 110.6, 98.8, 74.1, 71.1, 70.6, 67.0, 61.3, 47.8, 47.6, 47.4, 47.2, 47.0, 29.7, 28.0, 25.7, 25.0. HRMS (ESI) m/z Calcd. for $C_{25} H_{33} O_8 N_4$ [M+H$^+$]: 517.2293, found 517.2283.

(E)-N-Hydroxy-3-(4-(6-(((2S,3R,4S,5S,6R)-3,4,5-trihydroxy-6-(hydroxymethyl)tetrahydro-2H-pyran-2-yl)oxy)naphthalen-2-yl)-1H-1,2,3-triazol-1-yl)acrylamide (**STR-V-105**). **STR-II-30** (202 mg, 0.37 mmol), compounds **5** (240 mg, 0.73 mmol), palladium (II) acetate (12.3 mg, 0.05 mmol) and tri(o-tolyl)phosphine (32.3 mg, 0.11 mmol) were dissolved in acetonitrile (3 mL) and the mixture was flushed with argon for 5 min. TEA (0.13 mL, 0.91 mmol) was added and the mixture heated under argon atmosphere at 90°C for 5 h. The mixture was filtered over a celite bed and the filtrate was evaporated *in vacuo*. The residue was purified via column chromatography, eluting with EtOAc:hexane 6:4 to furnish the *O*-trityl protected intermediate (Yield: 87 mg, 29 %). The *O*-trityl intermediate (87 mg, 0.11 mmol was dissolved into DCM (1.5 mL), TIPS (0.1 mL) and TFA (0.5 mL) and the reaction was stirred at rt for 30 min. Solvent was evaporated off and the residue was stirred in MeOH (2 mL) and NaOMe (0.1 mL) at rt for 14 h and purified as described for the synthesis of **STR-V-53** to furnish **STR-V-105** as brownish-white solid (Yield: 24 mg, 54 %). $^{1}$H NMR (700 MHz, DMSO-*d6*) δ 10.77 (s, 1H), 8.01 (s, 1H), 7.89 (d, *J* = 8.9 Hz, 1H), 7.83 (d, *J* = 8.6 Hz, 1H), 7.67 (dd, *J* = 8.6, 1.7 Hz, 1H), 7.60 – 7.56 (m, 2H), 7.30 (dd, *J* = 8.9, 2.5 Hz, 1H), 6.54 (d, *J* = 15.8 Hz, 1H), 5.56 (d, *J* = 1.8 Hz, 1H), 3.89 (dd, *J* = 3.4, 1.9 Hz, 1H), 3.73 (dd, *J* = 9.2, 3.4 Hz, 1H), 3.60 (dd, *J* = 11.7, 2.1 Hz, 1H), 3.52 (t, *J* = 9.4 Hz, 1H), 3.48 (dd, *J* = 11.7, 6.1 Hz, 1H), 3.43 (m, 3.44-3.41, 1H). $^{13}$C NMR (176 MHz, MeOD) δ 155.3, 140.5, 135.3, 130.7, 129.8, 129.6, 128.8, 127.6, 123.5, 119.2, 116.4, 110.6, 98.8, 74.2, 71.0, 70.6, 67.0, 61.3, 48.5. HRMS (ESI) m/z Calcd. for $C_{19} H_{22} O_8 N$ [M+H$^{+}$]: 392.1340, found 392.1338.

(2S,3R,4S,6R)-4-(Dimethylamino)-6-methyltetrahydro-2H-pyran-2,3-diyl diacetate (**STR-V-160**) Desosamine (2.00 g, 11.41 mmol) and acetic anhydride (3.24 mL, 34.24 mmol) were dissolved in DCM (60 mL), DMAP (558 mg, 4.57 mmol) was added and the mixture was stirred at rt overnight. $H_2O$ (50 mL) was added and the two layers separated. The organic layer was dried over $Na_2SO_4$ and the solvent was evaporated off. The crude was purified via column chromatography, eluting with DCM:MeOH 9.5:0.5 to furnish **STR-V-160** as white solid (Yield: 2.48 g, 80 %). $^{1}$H NMR (400 MHz, $CDCl_3$) δ 6.22 (d, *J* = 3.6 Hz, 1H), 5.03 (ddd, *J* = 11.1, 3.7, 0.8 Hz, 1H), 4.04 (dqd, *J* = 12.3, 6.1, 2.2 Hz, 1H), 3.64 (d, *J* = 0.8 Hz, 1H), 3.13 (td, *J* = 11.6, 3.9

Hz, 1H), 2.29 (d, *J* = 0.8 Hz, 6H), 2.13 (d, *J* = 0.9 Hz, 3H), 2.04 (d, *J* = 0.8 Hz, 3H), 1.85 (ddd, *J* = 13.2, 4.2, 2.5 Hz, 1H), 1.48 – 1.33 (m, 1H), 1.21 (dd, *J* = 6.2, 0.9 Hz, 3H).

(2S,3R,4S,6R)-4-(Dimethylamino)-2-(4-iodophenoxy)-6-methyltetrahydro-2H-pyran-3-yl acetate (**STR-V-161**). **STR-V-160** (104 mg, 0.4 mmol) and 4-iodophenol (135 mg, 0.6 mmol) were dissolved in DCM (3 mL). The mixture was cooled to ice bath and flushed with argon for 10 min. Boron trifluoride etherate (1 mL, 8 mmol) was added slowly and dropwisely, ice bath was removed after 10 min stirring and the reaction was stirred at rt 12 h. The reaction was partitioned between DCM (30 mL) and water (30 mL) and the two layers separated. The organic layer was washed with water (30 mL), dried over $Na_2SO_4$ and the solvent evaporated off. The crude was purified using preparative TLC eluting with ethyl acetate solvent to furnish **STR-V-161** as white solid (Yield: 161 mg, 95 %). $^1$H NMR (400 MHz, $CDCl_3$) δ 7.60 – 7.51 (m, 2H), 6.80 – 6.72 (m, 2H), 5.07 (dd, *J* = 10.6, 7.5 Hz, 1H), 4.89 (dd, *J* = 7.5, 2.1 Hz, 1H), 3.83 – 3.54 (m, 1H), 2.82 (ddd, *J* = 12.3, 10.4, 4.3 Hz, 1H), 2.31 (d, *J* = 2.3 Hz, 6H), 2.16 – 2.03 (m, 3H), 1.87 – 1.78 (m, 1H), 1.61 (s, 1H), 1.46 (q, *J* = 12.8, 12.2 Hz, 1H), 0.89 – 0.77 (m, 1H).

(2S,3R,4S,6R)-4-(Dimethylamino)-2-(4-ethynylphenoxy)-6-methyltetrahydro-2H-pyran-3-yl acetate (**STR-V-163**). **STR-V-161** (221 mg, 0.53 mmol) was reacted with TMS-acetylene (0.1 mL, 0.65 mmol) in the presence of CuI (4mg, 0.02mmol) and bis(triphenylphosphine)palladium (II) dichloride (7.6 mg, 0.01 mmol) in TEA (1.85 mL, 13.25 mmol) and THF (3 mL) as described for the 1$^{st}$ step of the synthesis of **STR-V-111** except that reaction was stopped after 15 h. The mixture was filtered through a celite bed and the filtrate was evaporated of to furnish the crude product (198 mg, 96.2 %). The crude product was treated with potassium carbonate (140 mg, 1.02 mmol) in MeOH (2 mL) for 2h, the solvent was evaporated off and the crude purified using preparative TLC, eluting with 5 % MeOH in DCM to furnish **STR-V-163** as yellowish-white solid (105 mg, 74 %). $^1$H NMR (700 MHz, DMSO-*d6*) δ 7.43 (d, *J* = 8.7 Hz, 2H), 7.08 (d, *J* = 8.8 Hz, 2H), 5.53 (d, *J* = 3.5 Hz, 1H), 4.00 (s, 1H), 3.90 (dqd, *J* = 12.5, 6.2, 2.2 Hz, 1H), 3.72 (d, *J* = 9.2 Hz, 1H), 2.46 (s, 7H), 2.42 – 2.39 (m, 3H), 1.88 (d, *J* = 12.8 Hz, 1H), 1.25 (s, 1H), 1.18 (d, *J* = 6.2 Hz, 1H), 1.09 (d, *J* = 6.2 Hz, 4H).

(2S,3R,4S,6R)-4-(Dimethylamino)-2-(4-(1-(7-(hydroxyamino)-7-oxoheptyl)-1H-1,2,3-triazol-4-yl)phenoxy)-6-methyltetrahydro-2H-pyran-3-yl acetate (**STR-V-165**). Following the procedure for the s **STR-V-167**, the reaction of **STR-V-163** (47 mg, 0.17 mmol), compound **3** (76 mg, 0.18 mmol), copper (II) sulfate (4.26 mg, 0.017 mmol), sodium ascorbate (13 mg, 0.07 mmol), in H2O (0.2 mL), tert-butanol (1 mL) and ethanol (1 mL), after preparative TLC eluting with DCM:MeOH:$NH_4OH$ 8.5:1.5:0.2, furnished the TBDPS protected intermediate (Yield: 62 mg, 52 %). The TBDPS protected intermediate (58 mg, 0.08 mmol) was reacted with CsF (25 mg, 0.17 mmol) in MeOH (2 mL) at rt for 30 min, solvent was evaporated and the crude was purified on preparative TLC, eluting with solution of 15% MeOH in DCM containing 2% $NH_4OH$ (1M) to furnish **STR-V-165** (Yield: 18.1 mg, 48 %). $^1$H NMR (700 MHz, DMSO-*d6*) δ 8.48 (s, 1H), 7.76 (dq, *J* = 8.8, 2.6, 2.1 Hz, 2H), 7.16 – 7.10 (m, 2H), 5.48 (d, *J* = 3.5 Hz, 1H), 4.35 (t, *J* = 7.1 Hz, 2H), 3.90 (dqd, *J* = 12.6, 6.2, 2.1 Hz, 1H), 3.60 (dd, *J* = 10.6, 3.5 Hz, 1H), 3.00 (ddd, *J* = 12.0, 10.5, 3.9 Hz, 1H), 2.28 (s, 5H), 1.83 (td, *J* = 7.3, 3.5 Hz, 4H), 1.74 (ddd, *J* = 12.8, 4.1, 2.3 Hz, 1H), 1.44 (p, *J* = 7.2 Hz, 2H), 1.29 – 1.21 (m, 6H), 1.06 (d, *J* = 6.2 Hz, 2H). $^{13}$C NMR (176 MHz, $CD_3OD$) δ 171.7, 157.6, 147.9, 135.8, 134.7, 133.7, 130.7, 128.3, 127.3, 120.3, 117.6,

98.3, 69.1, 66.0, 60.7, 53.8, 52.4, 50.7, 40.6, 32.9, 31.2, 30.3, 28.6, 26.8, 25.6, 21.2, 19.4. HRMS (ESI) m/z Calcd. for $C_{23}$ $H_{36}$ $O_5$ $N_5$ [M+H$^+$]: 462.2711, found 462.2696.

**Figure S2.** Molecular docking predicted interactions of **STR-V-53** and **STR-V-165** at the active site of (a) HDAC2 (PDB:4LXZ); **STR-V-53** (b1) and **STR-V-165** (b2) at the active site of HDAC6 (5G0G); and the **STR-V-53** (c) and **STR-V-165** (d) at the active site of GLUT-1 (PDB: 4PYP).

A.

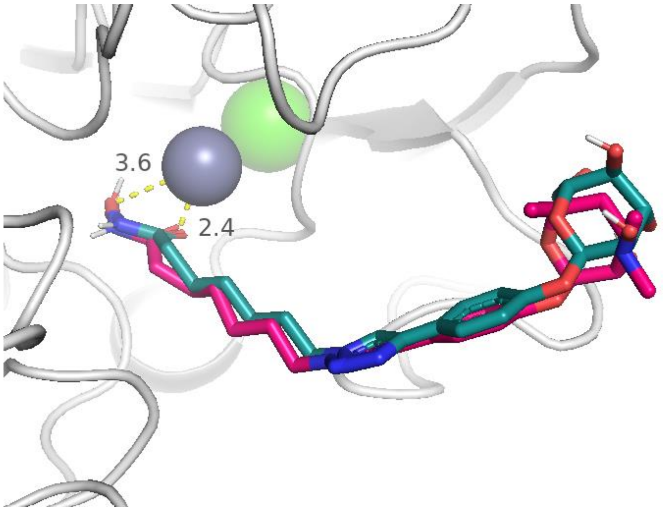


B1.

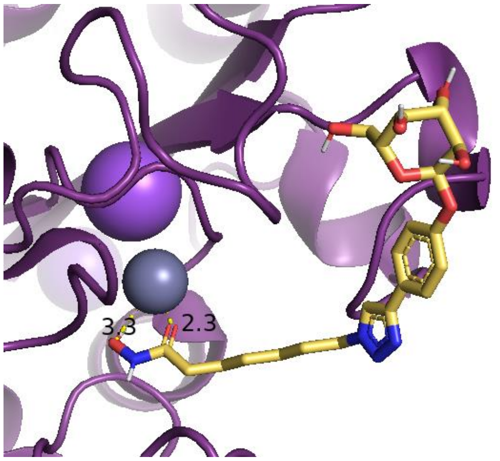


B2.

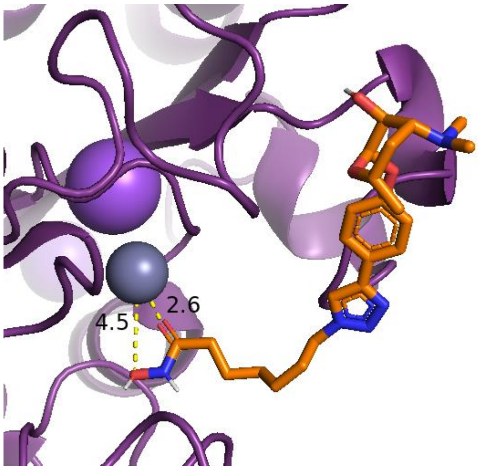

C.

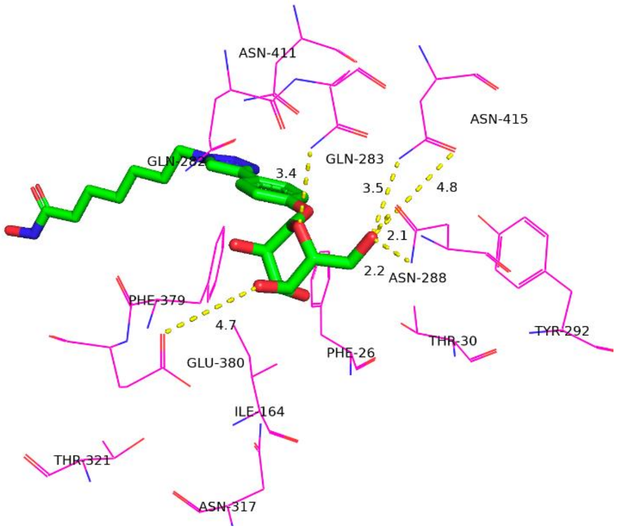
ASN-411
ASN-415
GLN-282
3.4
GLN-283
3.5
4.8
2.1
2.2
ASN-288
PHE-379
4.7
GLU-380
PHE-26
THR-30
TYR-292
ILE-164
THR-321
ASN-317

D.

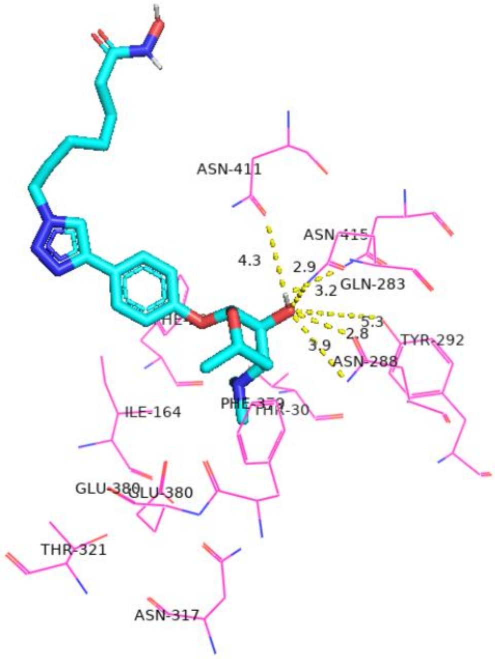
ASN-411
ASN-415
4.3
2.9
GLN-283
3.2
5.3
2.8
3.9
TYR-292
ASN-288
ILE-164
PHE-379
THR-30
GLU-380
GLU-380
THR-321
ASN-317

**Table S1**. HDAC isoform inhibition activities of glycosylated HDACi ($IC_{50}$ in nM)

| | **HDAC1** | **HDAC2** | **HDAC6** | **HDAC8** |
|---|---|---|---|---|
| **STR-V-167** 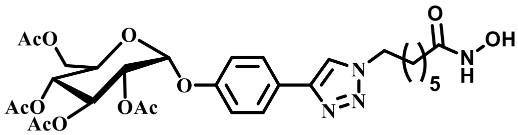 | 31.8 | 66.4 | 2.61 | 311 |
| **STR-I-195** 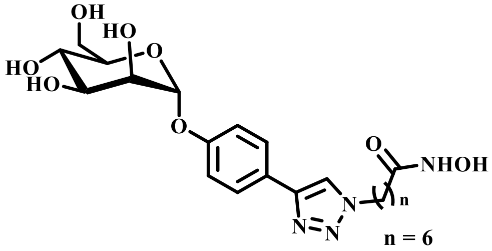 | 14.6 | 29.1 | 2.51 | 464 |
| **STR-V-165** 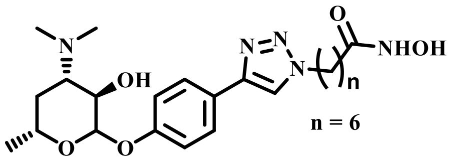 | 10.3 | 27.5 | 5.22 | 806 |
| **STR-V-176** 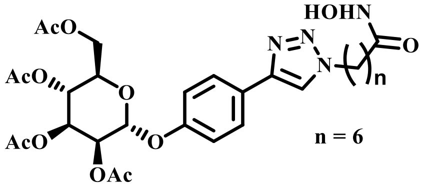 | 1.43 | 4.2 | 1.05 | 271 |
| **STR-V-177** 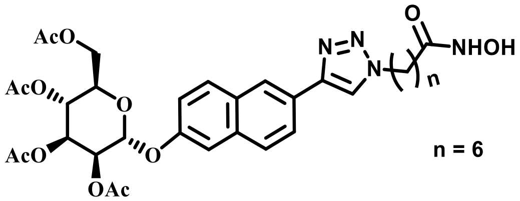 | 2.20 | 13.2 | 1.65 | 718 |
| **STR-II-36** 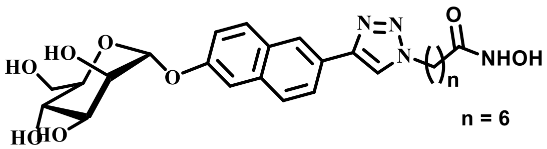 | 3.28 | 10.4 | 0.72 | 319 |
| **STR-V-114** 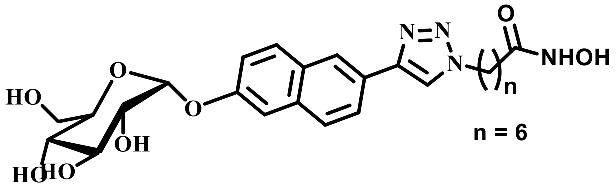 | 20.3 | 61.5 | 4.5 | 2130 |

| | | | | |
|---|---|---|---|---|
| **STR-V-115** 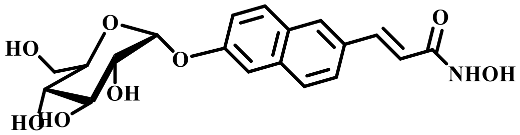 | 427 | 1030 | 16.5 | 461 |
| **STR-V-105** 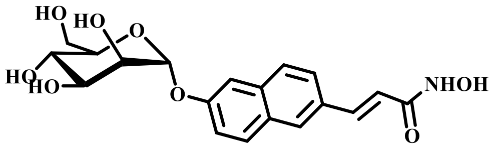 | 697 | 1460 | 30.9 | 771 |
| **STR-V-48** 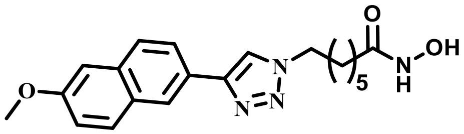 | NT* | NT* | NT* | NT* |
| **TSA** 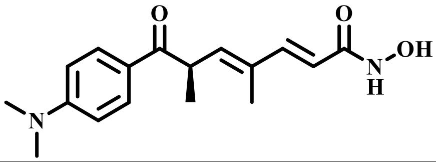 | 3.56 | 10.1 | 1.70 | 620 |

*Reported $IC_{50}$ = 15. 3 nM. Assayed using for HeLa Nuclear extract in a *Fluor de Lys* assay[27]

**Table S2.** Anti-proliferation effects of other glycosylated HDACi ($IC_{50}$ in μM). *NI=no inhibition up to 100μM. *NT= not tested

| Compound name | Hep-G2 (μM) | A549 (μM) | VERO (μM) |
|---|---|---|---|
| **STR-V-114** | 6.7±1.6 | 77.2±4.3 | 34.1±3.0 |
| **STR-V-176** | 8.0±0.9 | 64.7±7.0 | 77.4±5.6 |
| **STR-I-195** | 22.3±2.0 | NI | 65.5±8.3 |
| **STR-V-177** | 7.4±0.9 | 39.6±0.6 | 19.2±0.9 |
| **STR-II-36** | 7.5±0.8 | 20.1±1.1 | 33.6±5.5 |
| **STR-V-105** | 78.5±11.6 | NI | NI |
| **STR-V-115** | 66.1±10.3 | NI | 82.1 |
| **STR-V-165** | 9.6±3.4 | 33.7±1.8 | 76.4 |
| **STR-V-167** | 10.1±1.2 | NI | NI |
| **STR-V-48** | 0.4±0.04 | 1.4±0.2 | 0.2±0.05 |
| **SORA** | 3.7±0.4 | 18.1±2.9 | 12.2±0.1 |

**Figure S3.** Effects of **STR-V-53** on the growth of cancer cells in the NCI-60 panel at 10µM single dose.

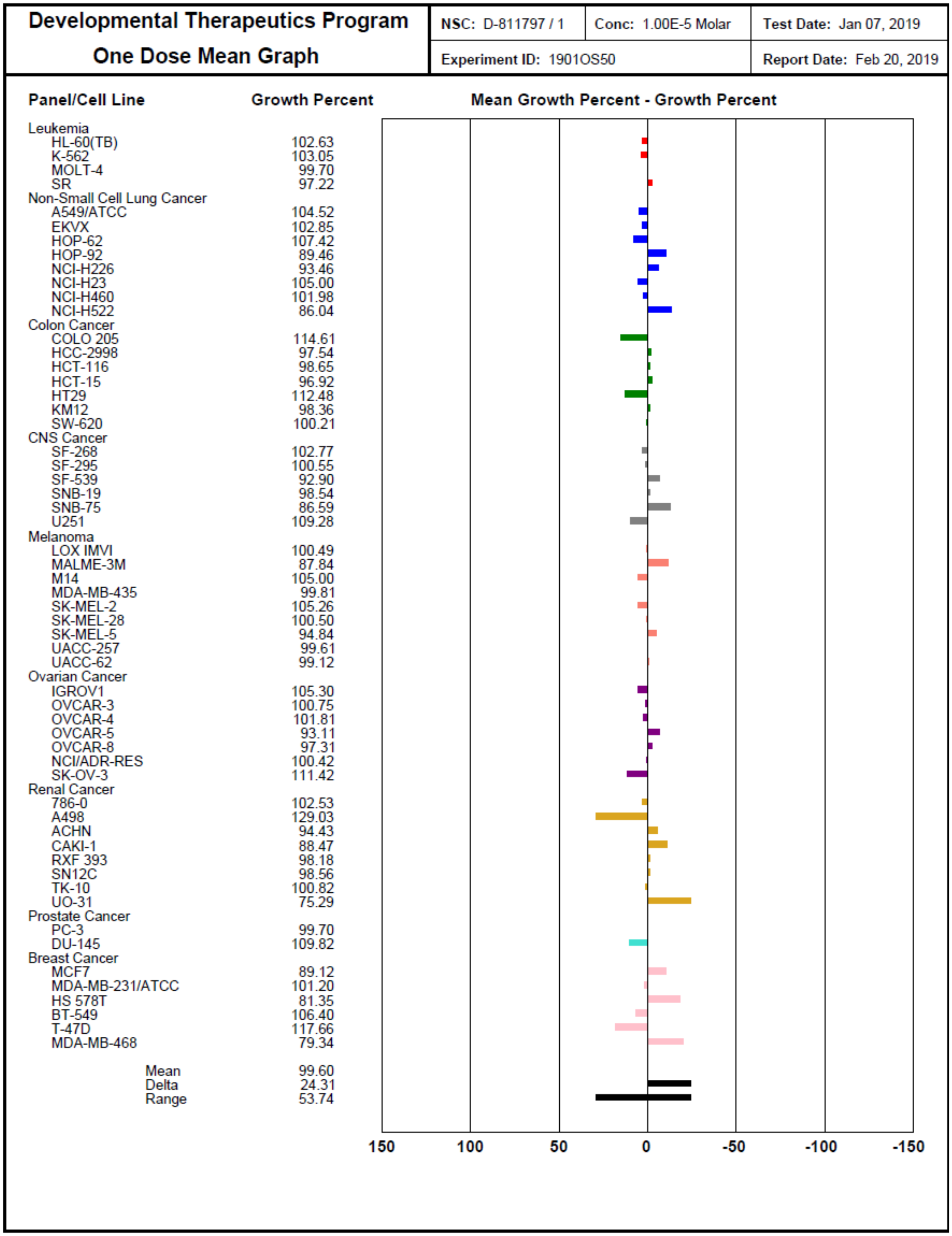

**Developmental Therapeutics Program**
**One Dose Mean Graph**

| | |
|---|---|
| **NSC:** D-811797 / 1 | **Conc:** 1.00E-5 Molar |
| **Test Date:** Jan 07, 2019 | **Experiment ID:** 1901OS50 |
| **Report Date:** Feb 20, 2019 | |

| Panel/Cell Line | Growth Percent |
|---|---|
| Leukemia | |
| HL-60(TB) | 102.63 |
| K-562 | 103.05 |
| MOLT-4 | 99.70 |
| SR | 97.22 |
| Non-Small Cell Lung Cancer | |
| A549/ATCC | 104.52 |
| EKVX | 102.85 |
| HOP-62 | 107.42 |
| HOP-92 | 89.46 |
| NCI-H226 | 93.46 |
| NCI-H23 | 105.00 |
| NCI-H460 | 101.98 |
| NCI-H522 | 86.04 |
| Colon Cancer | |
| COLO 205 | 114.61 |
| HCC-2998 | 97.54 |
| HCT-116 | 98.65 |
| HCT-15 | 96.92 |
| HT29 | 112.48 |
| KM12 | 98.36 |
| SW-620 | 100.21 |
| CNS Cancer | |
| SF-268 | 102.77 |
| SF-295 | 100.55 |
| SF-539 | 92.90 |
| SNB-19 | 98.54 |
| SNB-75 | 86.59 |
| U251 | 109.28 |
| Melanoma | |
| LOX IMVI | 100.49 |
| MALME-3M | 87.84 |
| M14 | 105.00 |
| MDA-MB-435 | 99.81 |
| SK-MEL-2 | 105.26 |
| SK-MEL-28 | 100.50 |
| SK-MEL-5 | 94.84 |
| UACC-257 | 99.61 |
| UACC-62 | 99.12 |
| Ovarian Cancer | |
| IGROV1 | 105.30 |
| OVCAR-3 | 100.75 |
| OVCAR-4 | 101.81 |
| OVCAR-5 | 93.11 |
| OVCAR-8 | 97.31 |
| NCI/ADR-RES | 100.42 |
| SK-OV-3 | 111.42 |
| Renal Cancer | |
| 786-0 | 102.53 |
| A498 | 129.03 |
| ACHN | 94.43 |
| CAKI-1 | 88.47 |
| RXF 393 | 98.18 |
| SN12C | 98.56 |
| TK-10 | 100.82 |
| UO-31 | 75.29 |
| Prostate Cancer | |
| PC-3 | 99.70 |
| DU-145 | 109.82 |
| Breast Cancer | |
| MCF7 | 89.12 |
| MDA-MB-231/ATCC | 101.20 |
| HS 578T | 81.35 |
| BT-549 | 106.40 |
| T-47D | 117.66 |
| MDA-MB-468 | 79.34 |
| Mean | 99.60 |
| Delta | 24.31 |
| Range | 53.74 |

**Figure S4.** Effects of representative glycosylated HDACi, **STR-V-114** (i) and **STR-I-195** (ii) on the acetylation status of H4 and tubulin in Hep-G2 cells.

(i)

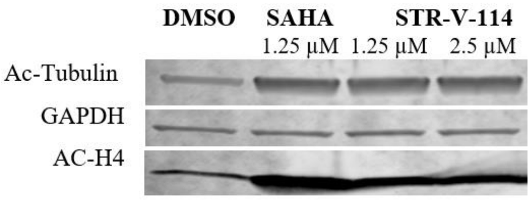


(ii)

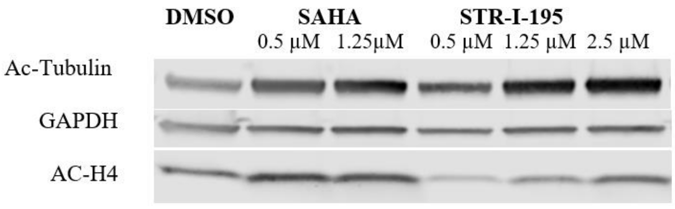


**Figure S5.** (a) Quantification of PD-1 and PD-L1 expression in Hep-G2. (b) Quantification of CD54 in Hep-G2. Bars show mean plus standard deviation; * P < 0.0332; ** P < 0.0021;***P<0.0002; ****P<0.00001.

a.

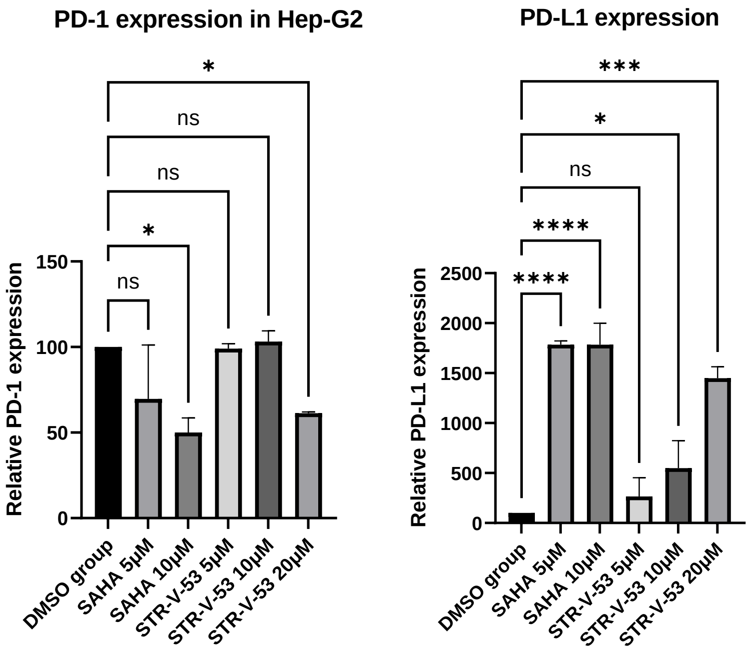

b.

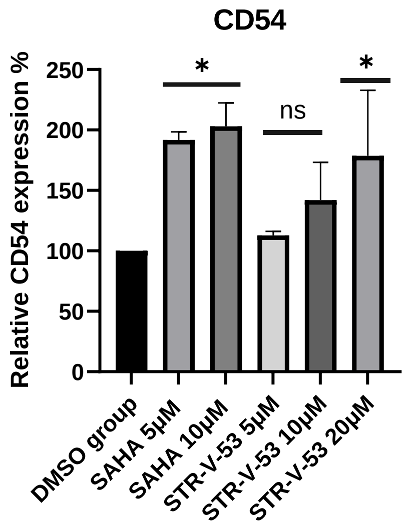


**Figure S6.** Similar to **STR-V-53**, compound **STR-I-195** induced apoptosis in Hep-G2 cells (a) and not in VERO cells (b).

a.

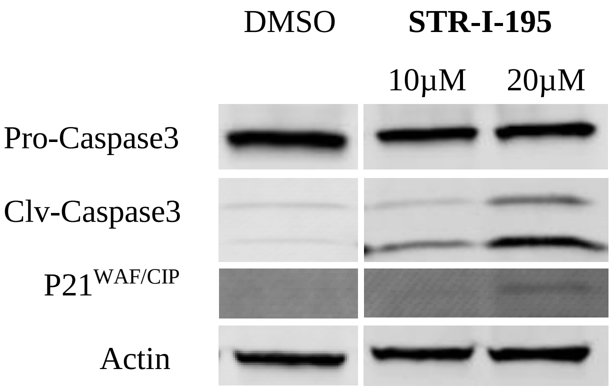


b.

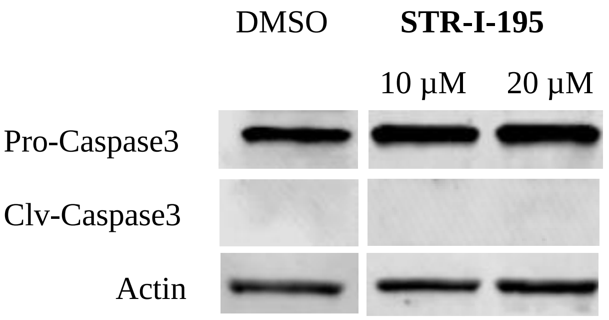

**Figure S7.** (a) Quantification of **STR-V-53**- and **STR-I-195**-induced changes to p21$^{WAF/CIP}$ expression and the ratio of pro-caspase3 to cleaved-caspase3 in the Hep-G2 cell. (b) Quantification of of **STR-V-53**- and **STR-I-195**-induced changes to the ratio of pro-caspase3 to cleaved-caspase3 of VERO cell. Bars show mean plus standard deviation; * P < 0.0332; ** P < 0.0021;***P<0.0002; ****P<0.00001.

a.

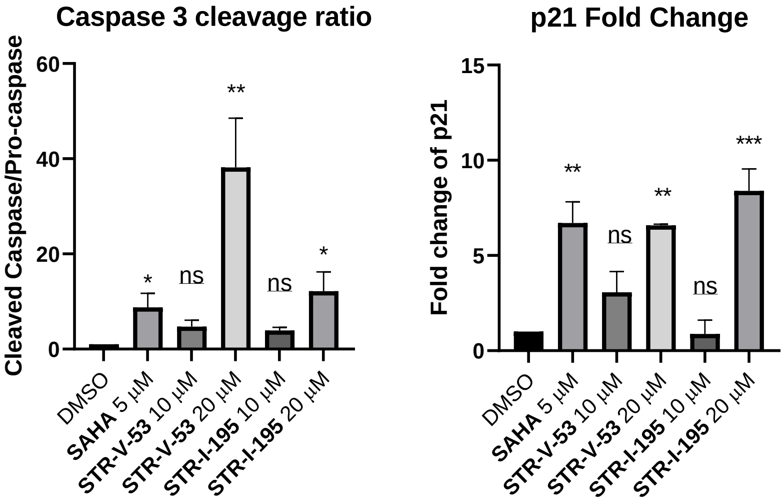


b.

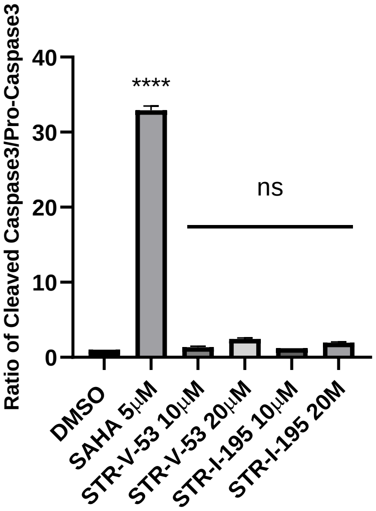

**Figure S8.** Stability of **STR-V-53** in mouse and human Plasmas.

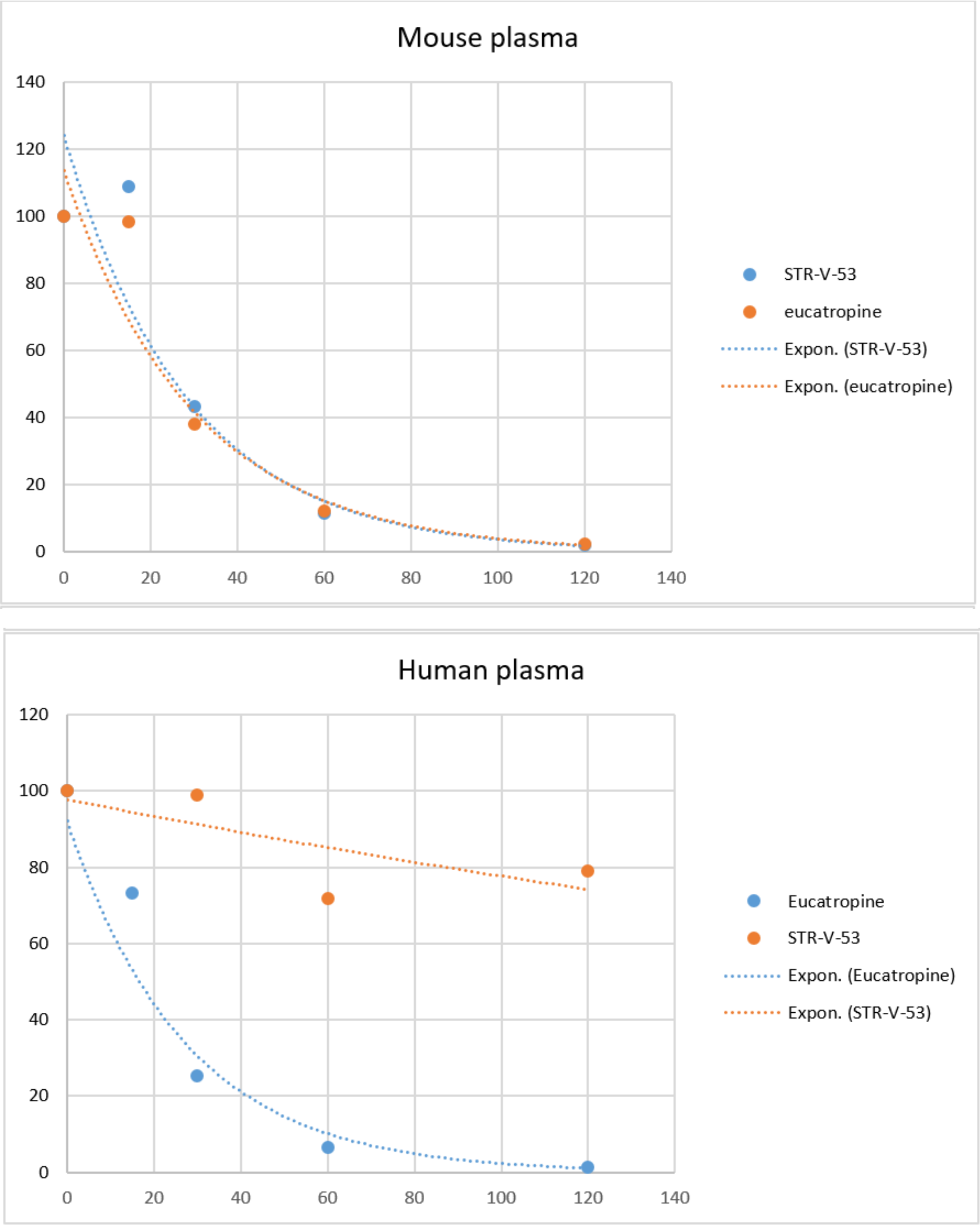

**Figure S9.** Sex-dependence in the plasma and liver distribution of **STR-V-53**. Data from 4 randomly selected mice (2 male and 2 female).

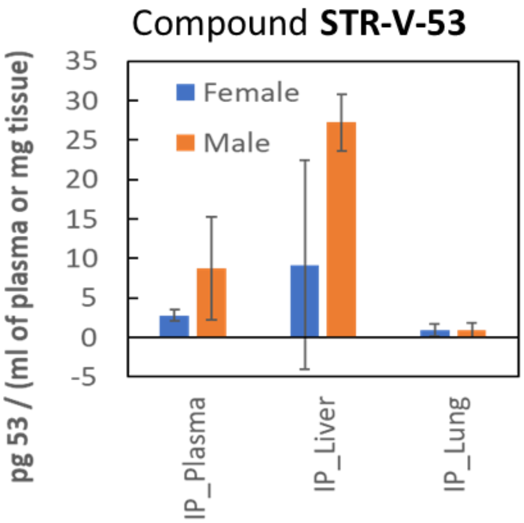

### NMRs

STR-V-167 ($^{1}H$)

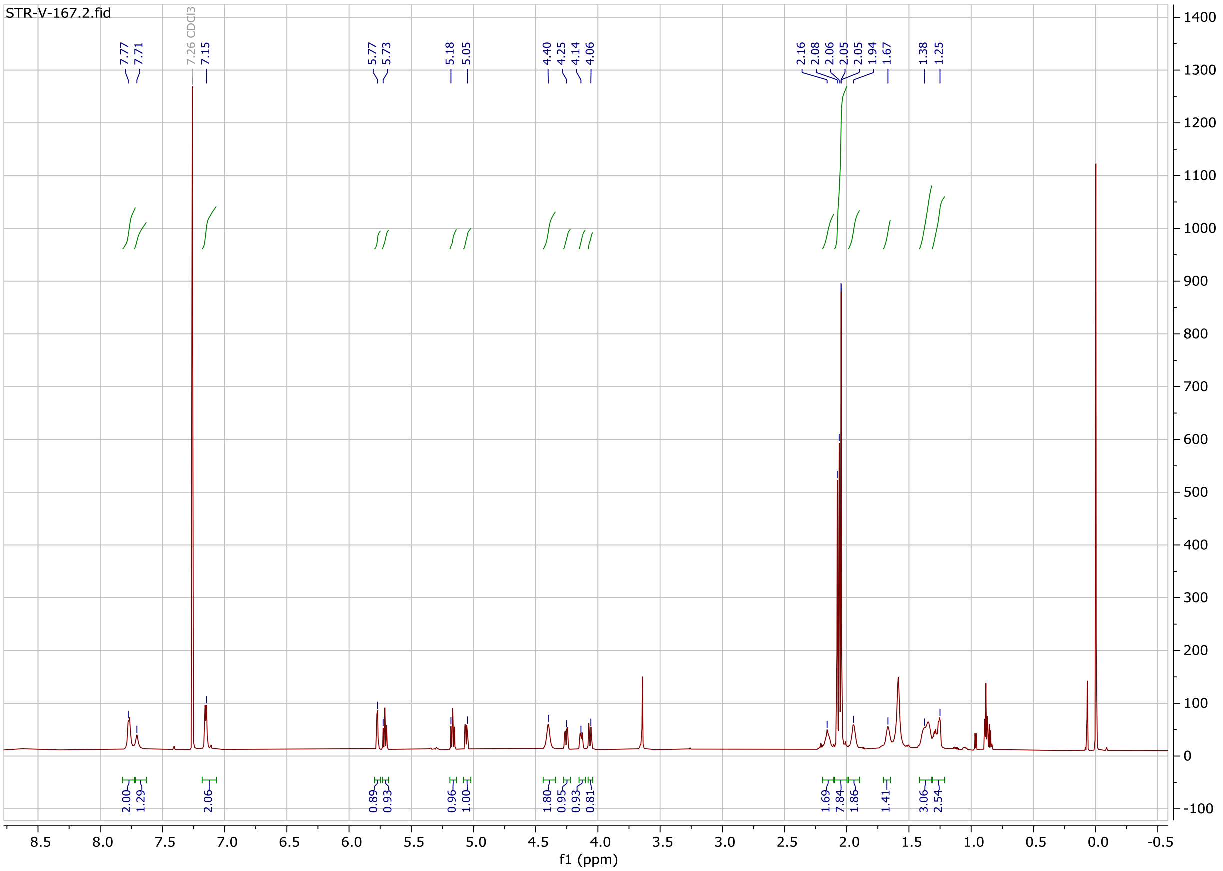

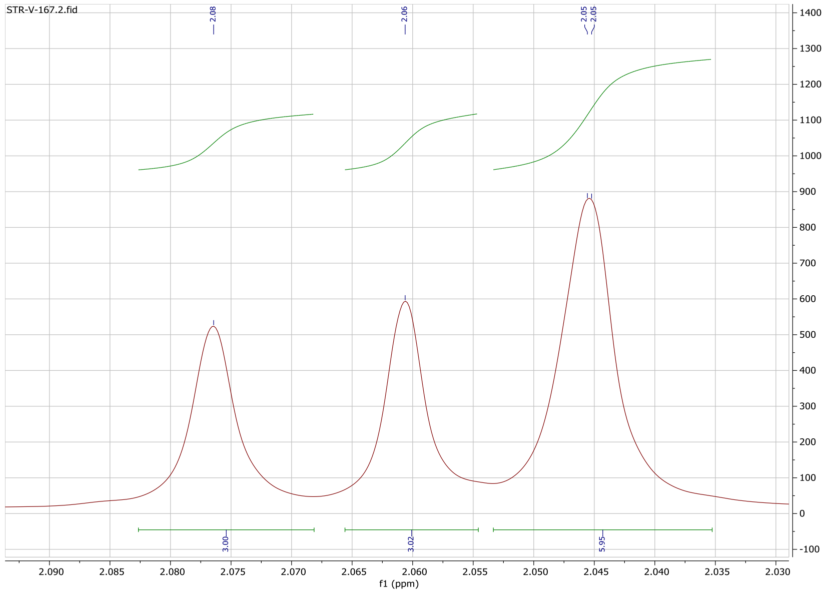

STR-V-167 ($^{13}C$)

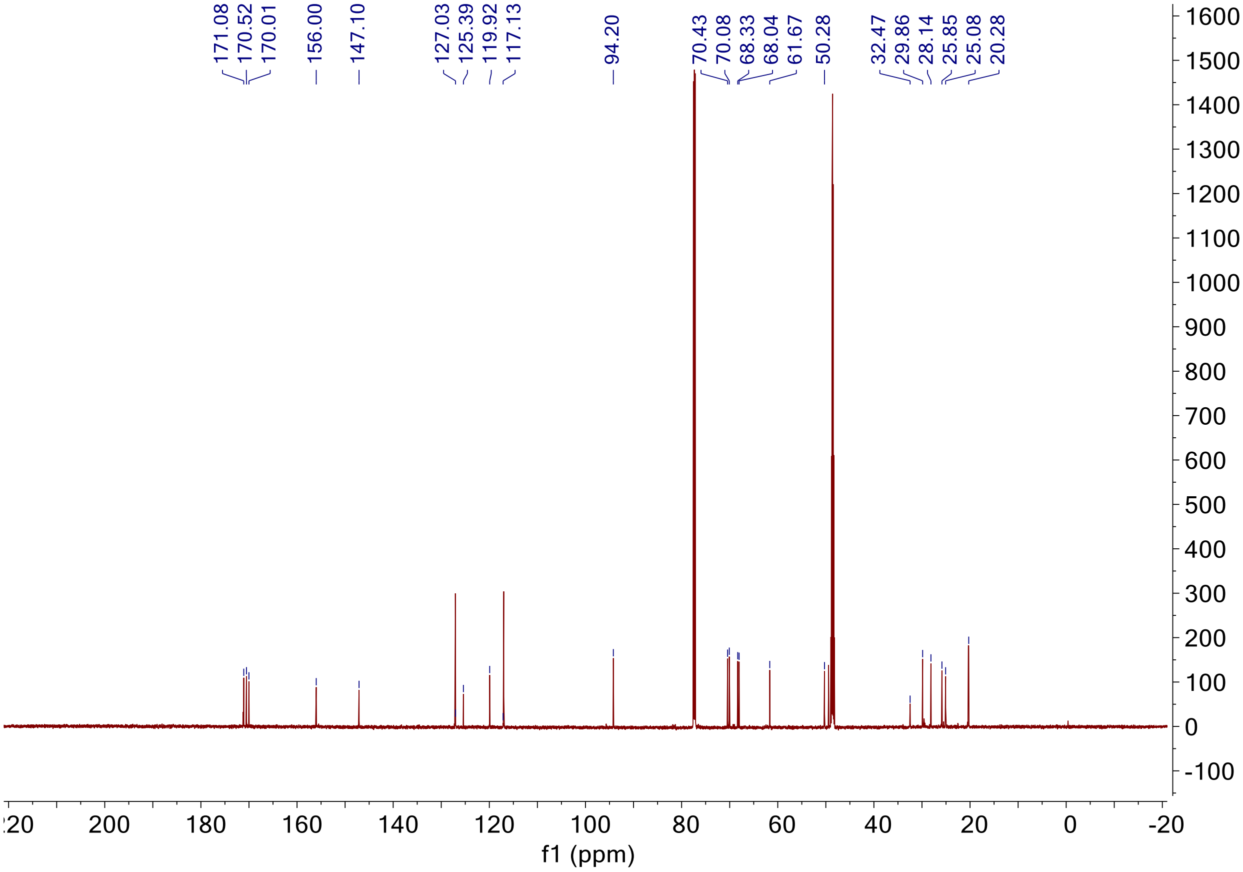
171.08
170.52
170.01
156.00
147.10
127.03
125.39
119.92
117.13
94.20
70.43
70.08
68.33
68.04
61.67
50.28
32.47
29.86
28.14
25.85
25.08
20.28
f1 (ppm)

STR-V-53 ($^{1}$H)

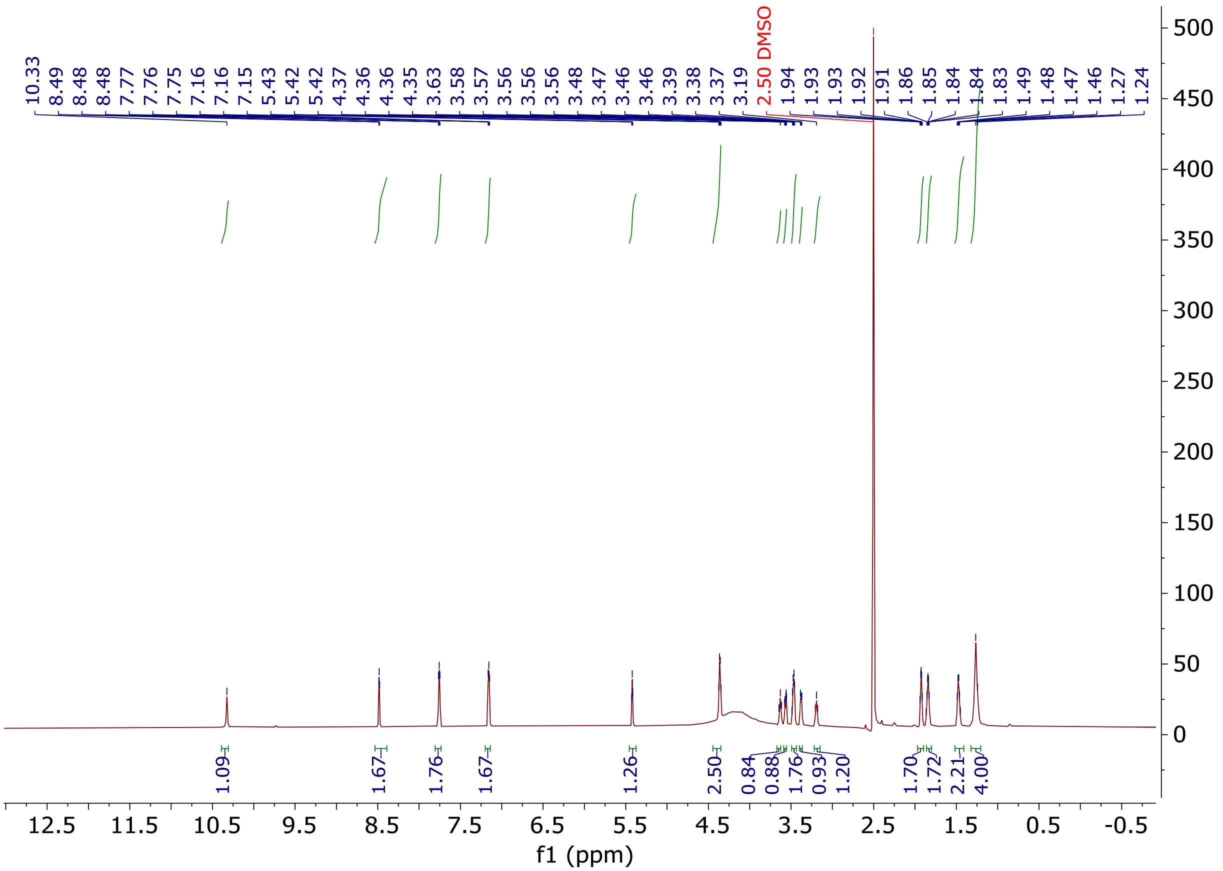

STR-V-53 ($^{13}C$)

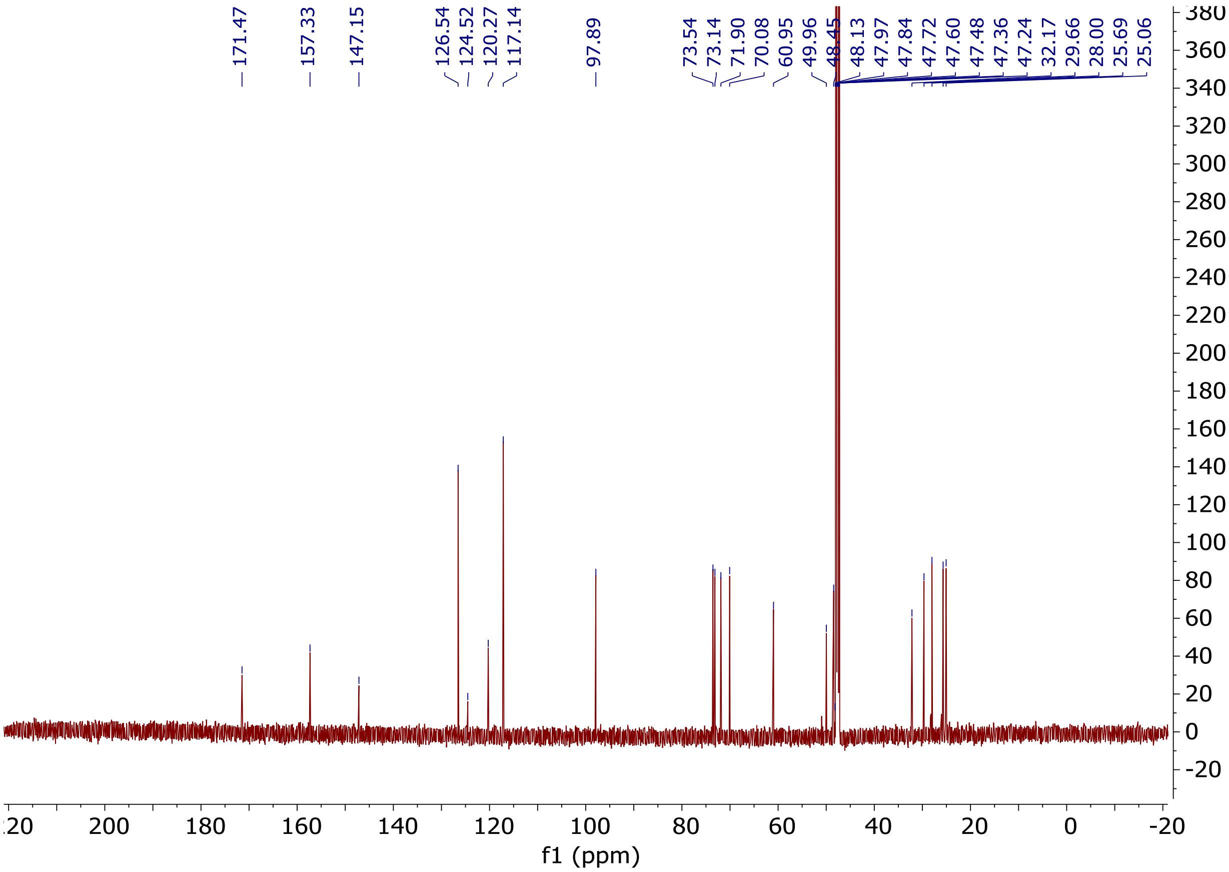
171.47
157.33
147.15
126.54
124.52
120.27
117.14
97.89
73.54
73.14
71.90
70.08
60.95
49.96
48.45
48.13
47.97
47.84
47.72
47.60
47.48
47.36
47.24
32.17
29.66
28.00
25.69
25.06
f1 (ppm)

STR-V-55 ($^{1}H$)

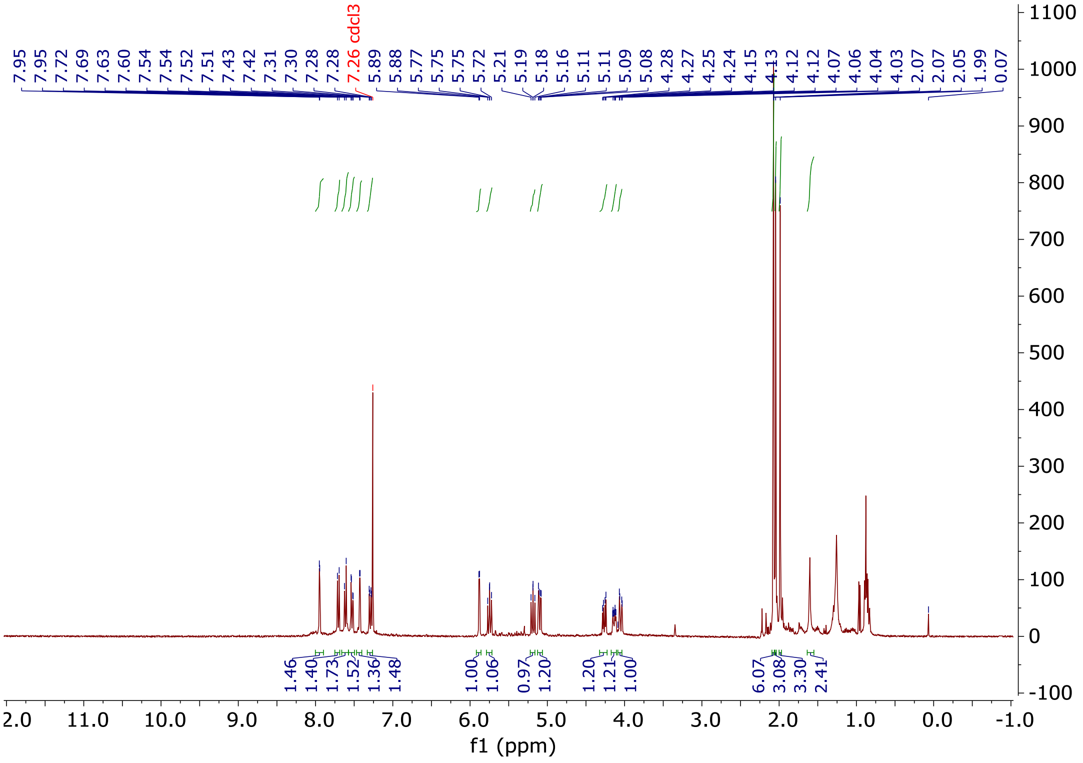
7.95
7.95
7.72
7.69
7.63
7.60
7.54
7.54
7.52
7.51
7.43
7.42
7.31
7.30
7.28
7.28
7.26 cdcl3
5.89
5.88
5.77
5.75
5.75
5.72
5.21
5.19
5.18
5.16
5.11
5.11
5.09
5.08
4.28
4.27
4.25
4.24
4.15
4.13
4.12
4.12
4.07
4.06
4.04
4.03
2.07
2.07
2.05
1.99
0.07
1.46
1.40
1.73
1.52
1.36
1.48
1.00
1.06
0.97
1.20
1.20
1.21
1.00
6.07
3.08
3.30
2.41
f1 (ppm)

STR-V-111 ($^1H$)

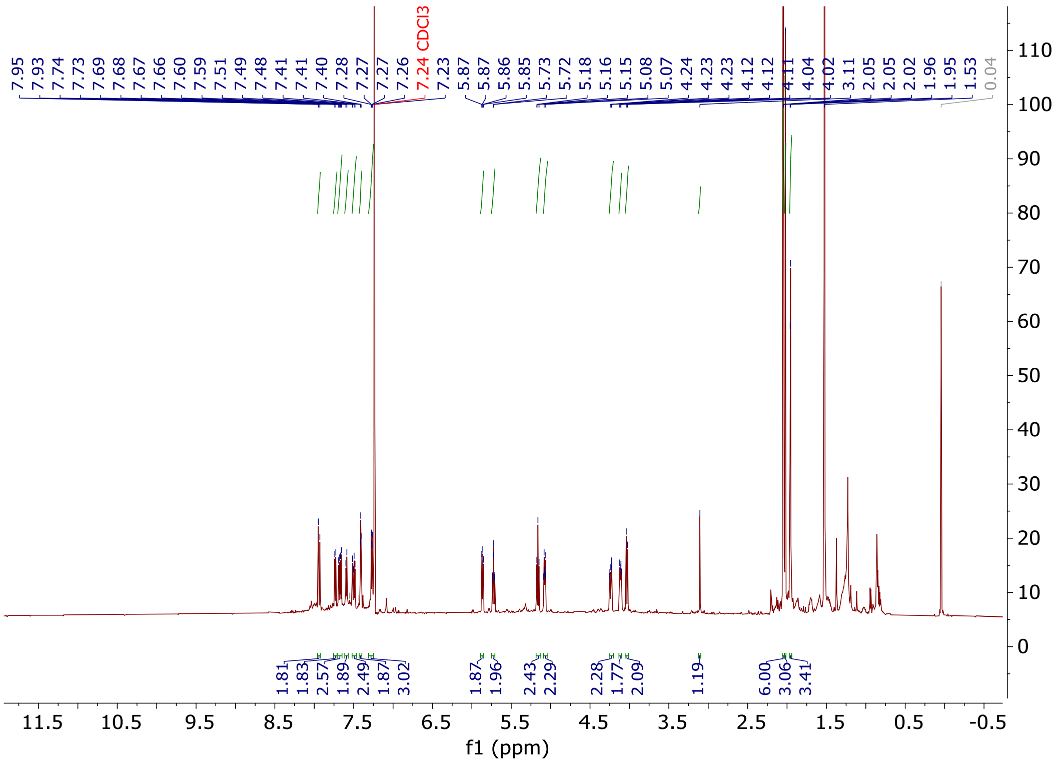

STR-V-114 ($^{1}$H)

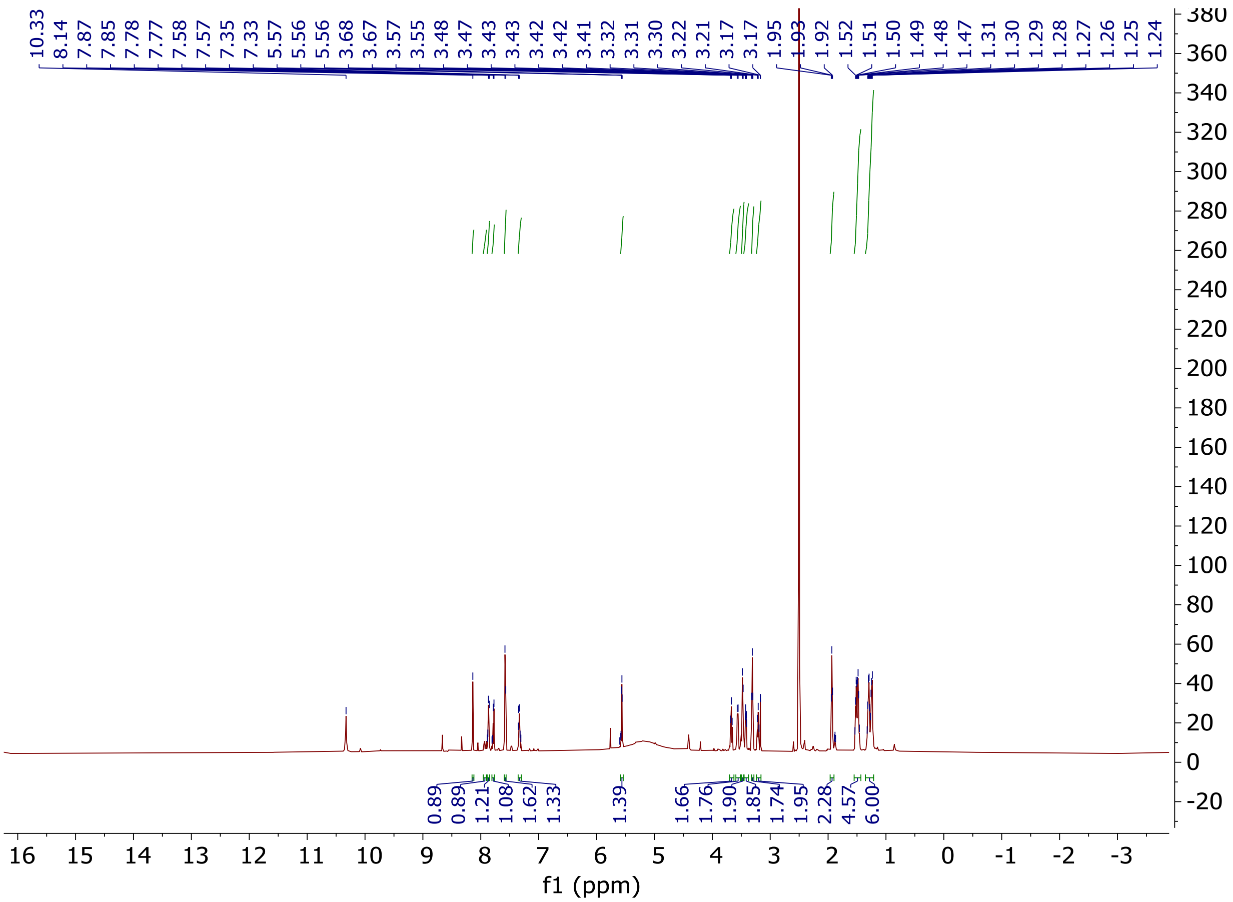

STR-V-115 ($^{1}$H)

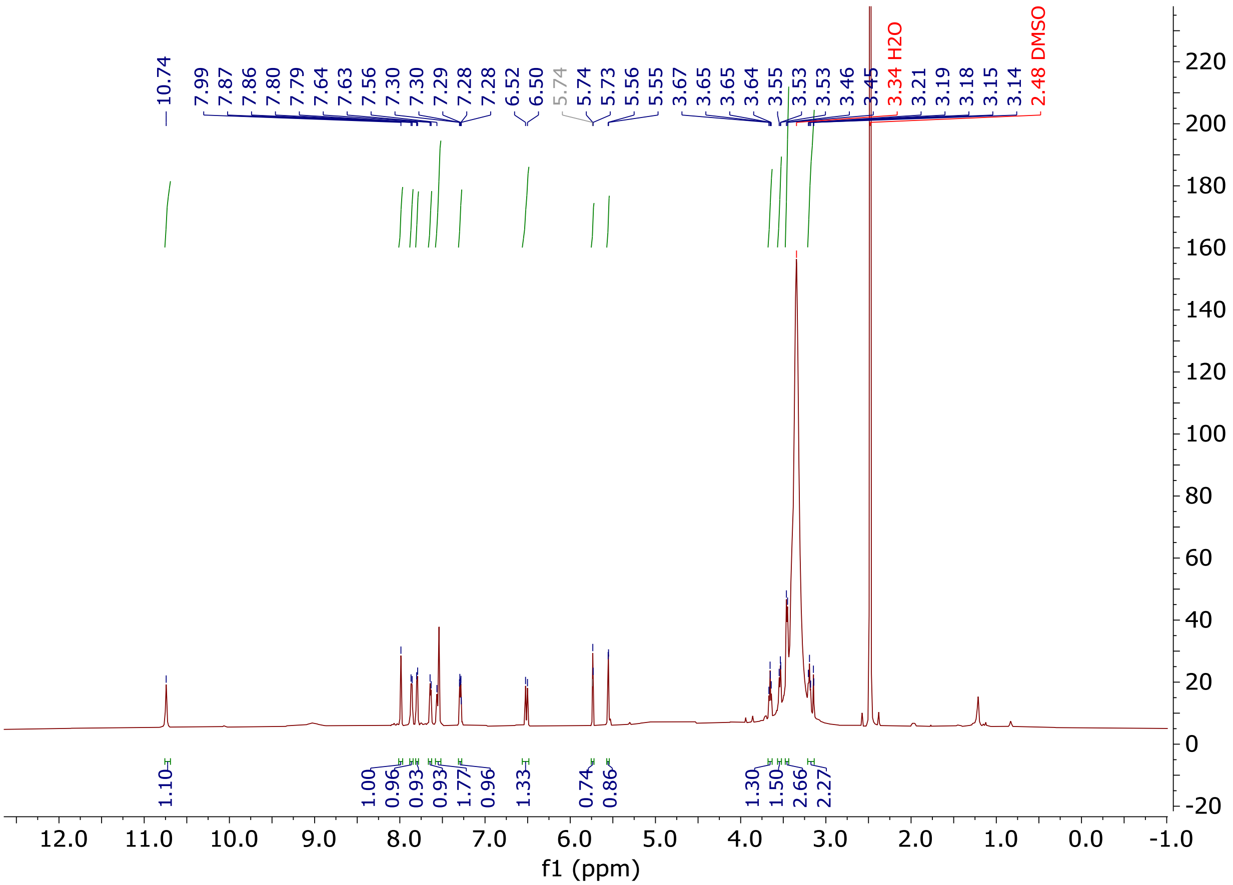

STR-V-115 ($^{13}C$)

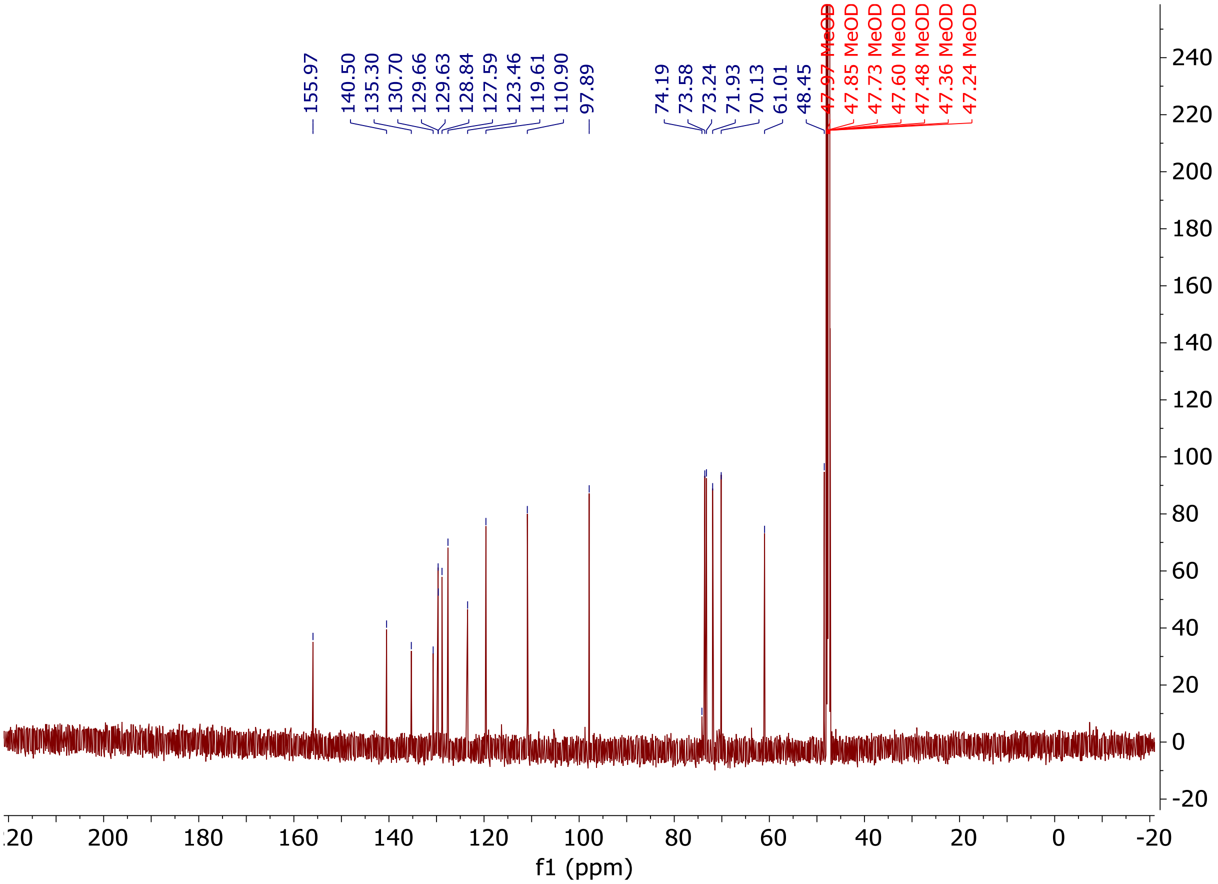
155.97
140.50
135.30
130.70
129.66
129.63
128.84
127.59
123.46
119.61
110.90
97.89
74.19
73.58
73.24
71.93
70.13
61.01
48.45
47.97 MeOD
47.85 MeOD
47.73 MeOD
47.60 MeOD
47.48 MeOD
47.36 MeOD
47.24 MeOD
f1 (ppm)

STR-V-176 ($^{1}H$)

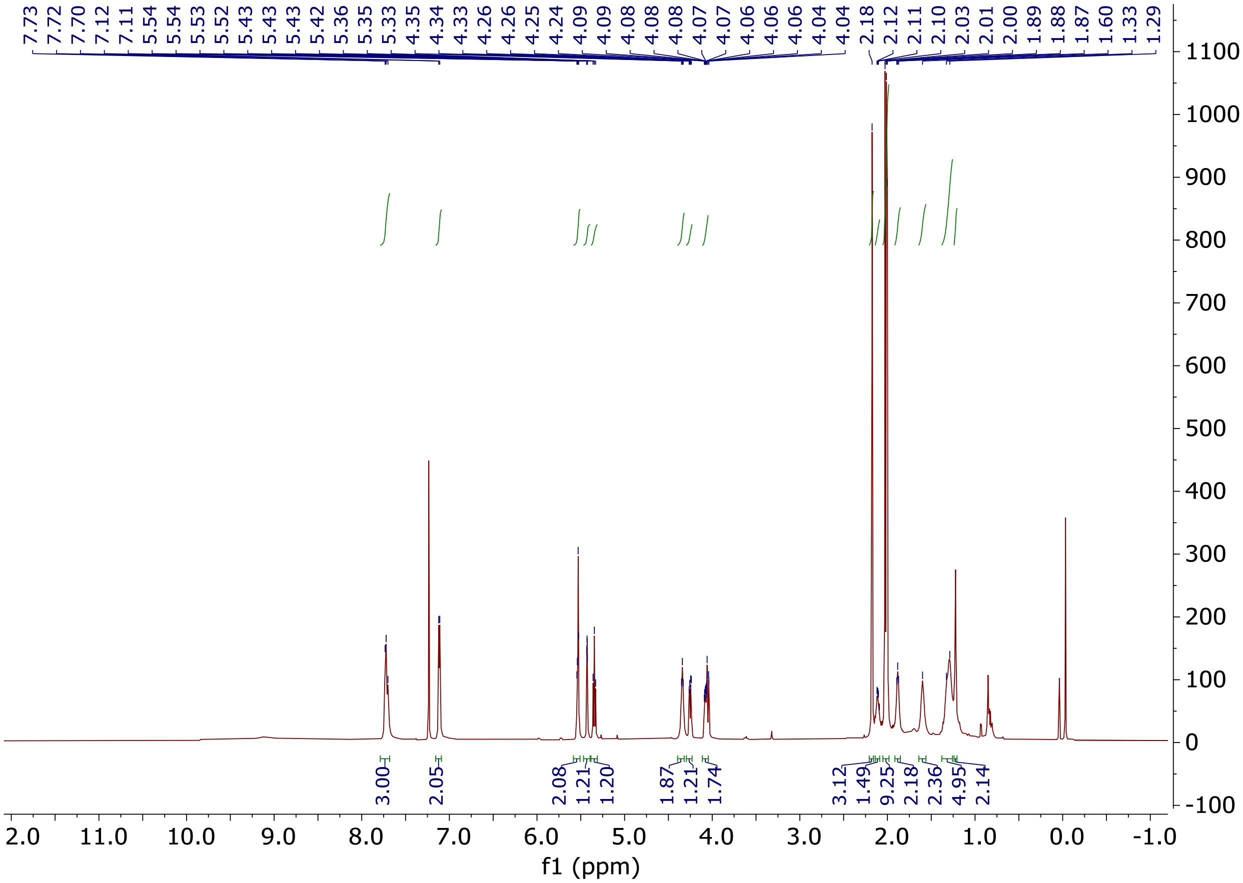

STR-V-176 ($^{13}C$)

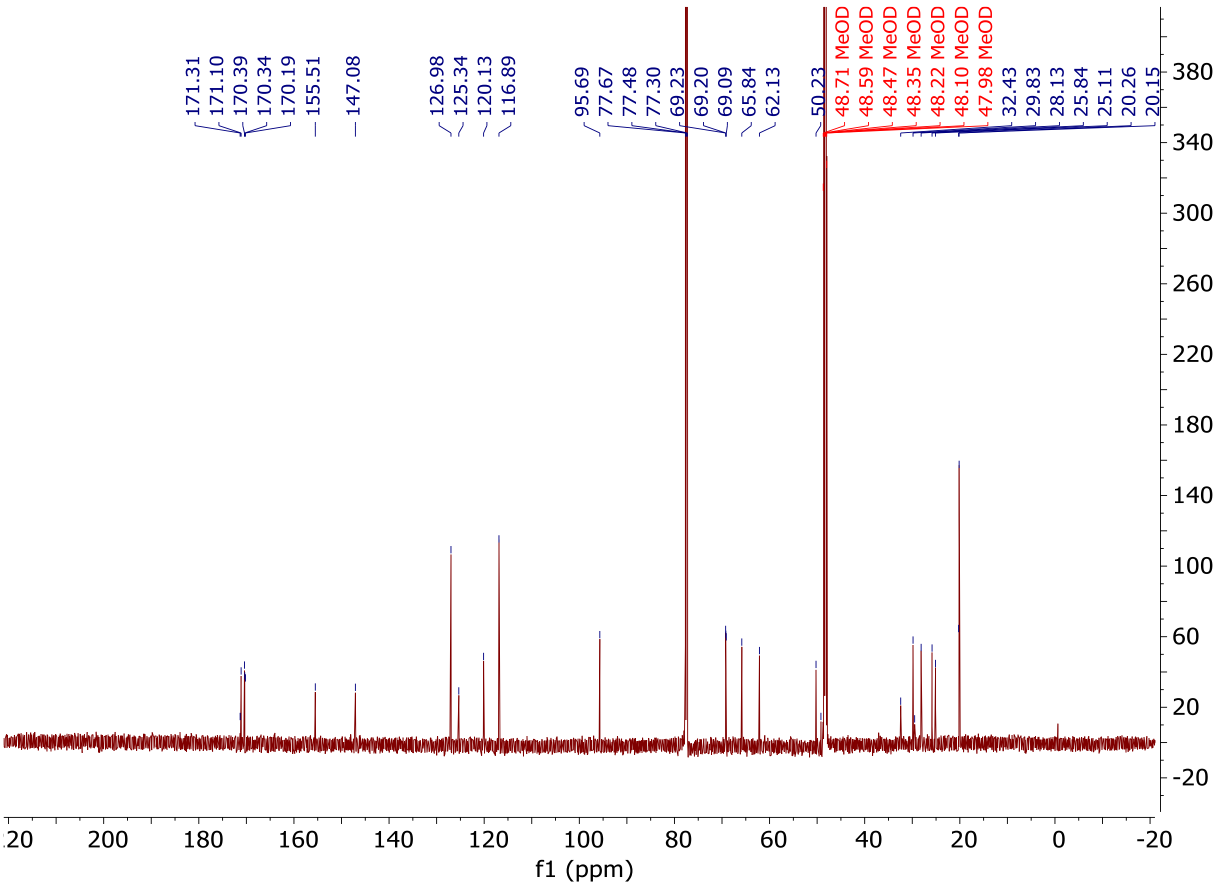
171.31
171.10
170.39
170.34
170.19
155.51
147.08
126.98
125.34
120.13
116.89
95.69
77.67
77.48
77.30
69.23
69.20
69.09
65.84
62.13
50.23
48.71 MeOD
48.59 MeOD
48.47 MeOD
48.35 MeOD
48.22 MeOD
48.10 MeOD
47.98 MeOD
32.43
29.83
28.13
25.84
25.11
20.26
20.15
f1 (ppm)

STR-V-195 ([1]H)

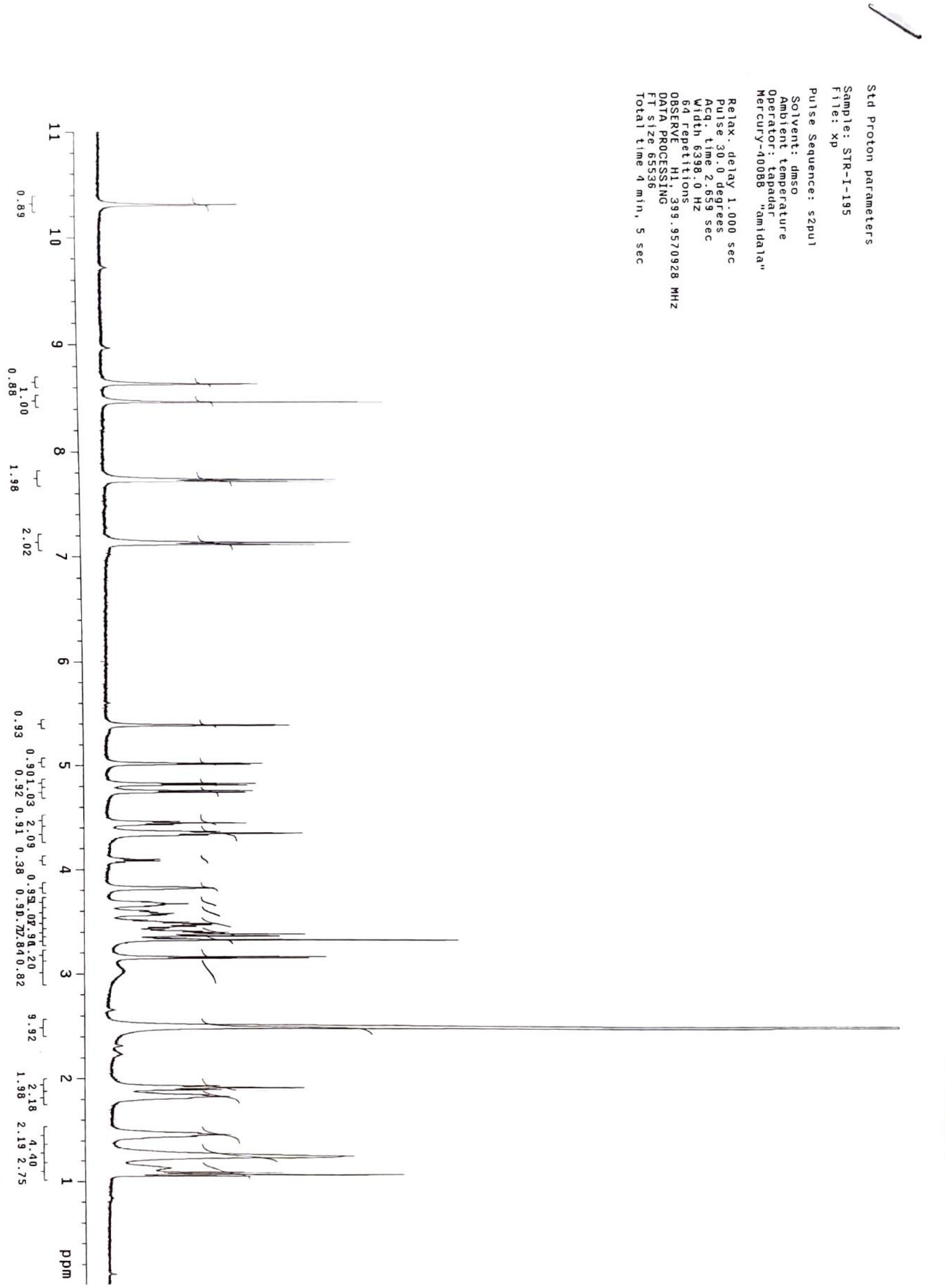
Std Proton parameters
Sample: STR-I-195
File: xp
Pulse Sequence: s2pul
Solvent: dmso
Ambient temperature
Operator: tapadar
Mercury-400BB "amidala"
Relax. delay 1.000 sec
Pulse 30.0 degrees
Acq. time 2.659 sec
Width 6398.0 Hz
64 repetitions
OBSERVE H1, 399.9570928 MHz
DATA PROCESSING
FT size 65536
Total time 4 min, 5 sec
11
10
9
8
7
6
5
4
3
2
1
ppm

STR-V-195 ($^{13}C$)

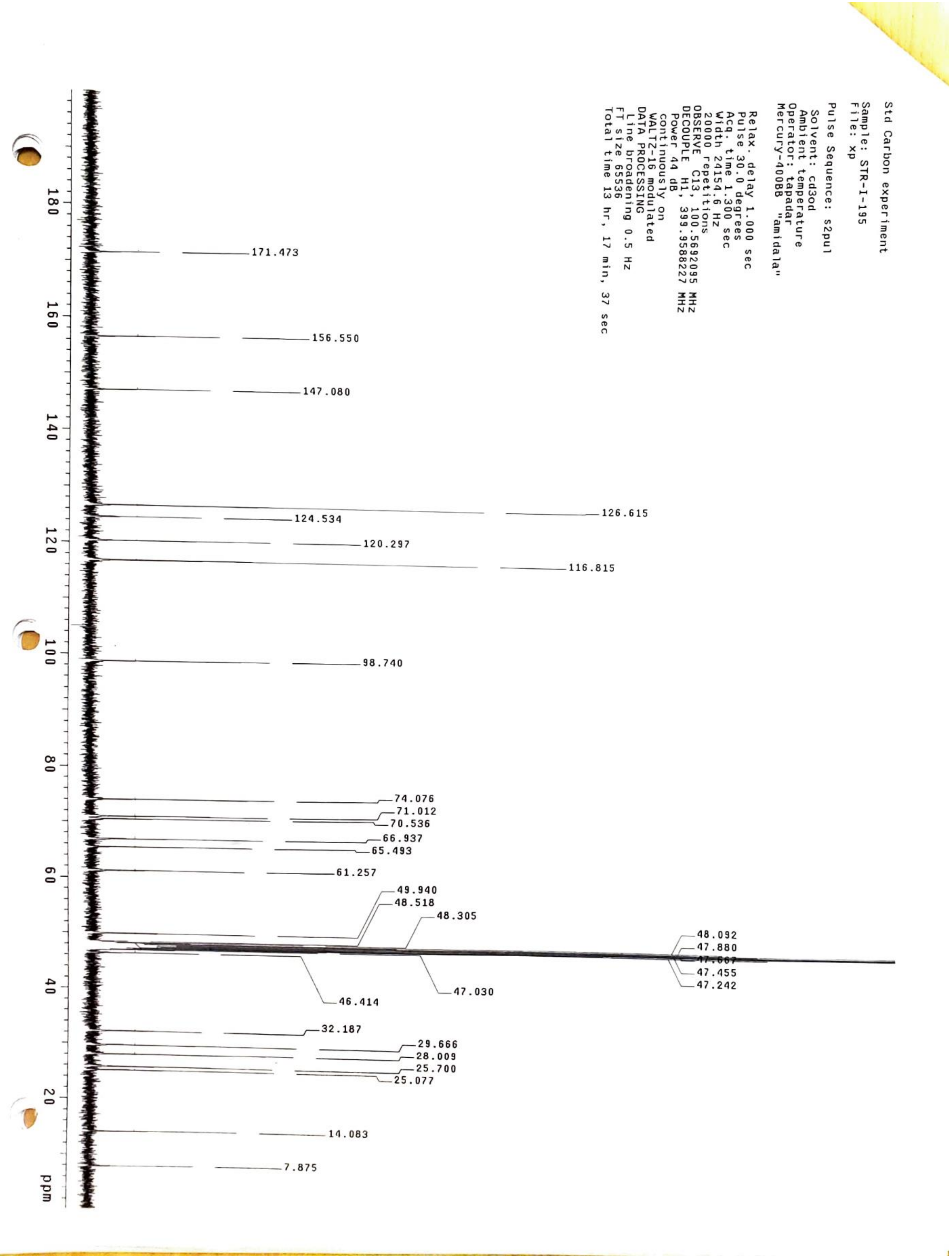

Std Carbon experiment
Sample: STR-I-195
File: xp
Pulse Sequence: s2pul
Solvent: cd3od
Ambient temperature
Operator: tapadar
Mercury-400BB "amidala"
Relax. delay 1.000 sec
Pulse 30.0 degrees
Acq. time 1.300 sec
Width 24154.6 Hz
20000 repetitions
OBSERVE C13, 100.5692095 MHz
DECOUPLE H1, 399.9588227 MHz
Power 44 dB
continuously on
WALTZ-16 modulated
DATA PROCESSING
Line broadening 0.5 Hz
FT size 65536
Total time 13 hr, 17 min, 37 sec
171.473
156.550
147.080
126.615
124.534
120.297
116.815
98.740
74.076
71.012
70.536
66.937
65.493
61.257
49.940
48.518
48.305
48.092
47.880
47.667
47.455
47.242
47.030
46.414
32.187
29.666
28.009
25.700
25.077
14.083
7.875
180
160
140
120
100
80
60
40
20
ppm

STR-II-30 ($^{1}H$)

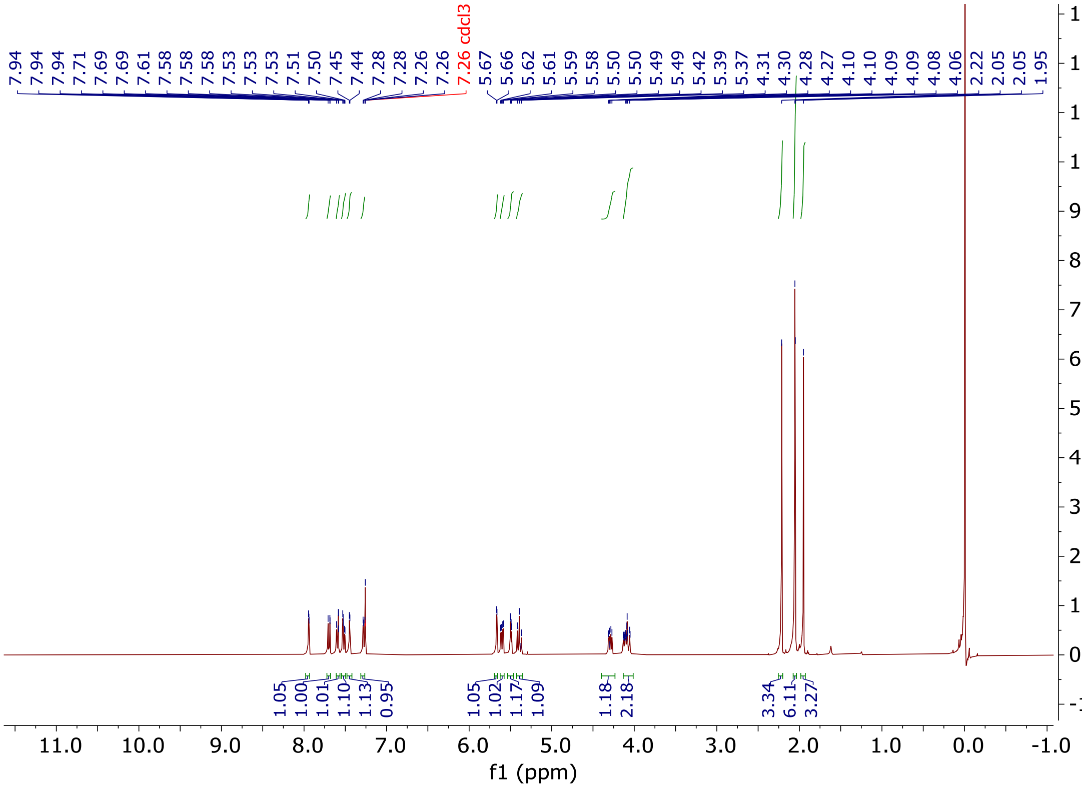

7.94 7.94 7.94 7.71 7.69 7.69 7.61 7.58 7.58 7.58 7.53 7.53 7.53 7.51 7.50 7.45 7.44 7.28 7.28 7.26 7.26
7.26 cdcl3
5.67 5.66 5.62 5.61 5.59 5.58 5.50 5.50 5.49 5.49 5.42 5.39 5.37 4.31 4.30 4.28 4.27 4.10 4.10 4.09 4.09 4.08 4.06 2.22 2.05 1.95
1.05 1.00 1.01 1.10 1.13 0.95 1.05 1.02 1.17 1.09 1.18 2.18 3.34 6.11 3.27
f1 (ppm)

STR-II-34 ($^1H$)

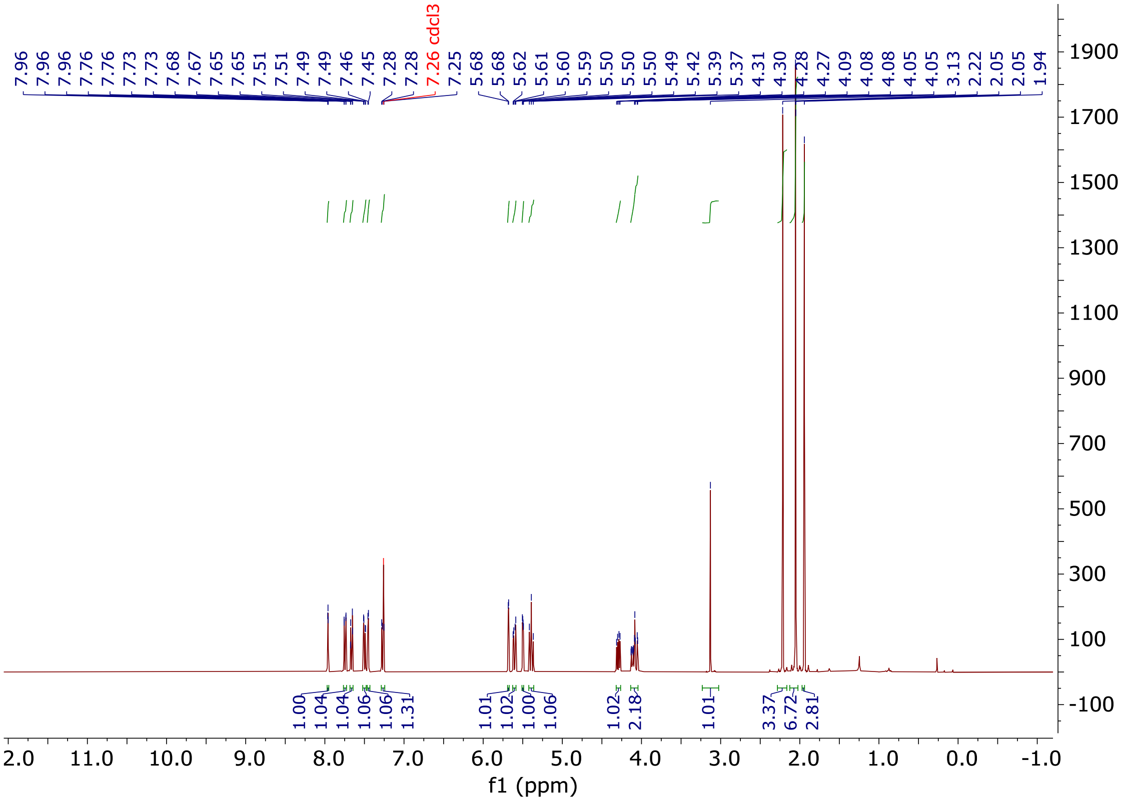

7.96
7.96
7.96
7.76
7.76
7.73
7.73
7.68
7.67
7.65
7.65
7.51
7.51
7.49
7.49
7.46
7.45
7.28
7.28
7.26 cdcl3
7.25
5.68
5.68
5.62
5.61
5.60
5.59
5.50
5.50
5.50
5.49
5.42
5.39
5.37
4.31
4.30
4.28
4.27
4.09
4.08
4.08
4.05
4.05
3.13
2.22
2.05
1.94
1.00
1.04
1.04
1.06
1.06
1.31
1.01
1.02
1.00
1.06
1.02
2.18
1.01
3.37
6.72
2.81
f1 (ppm)

STR-V-177 ($^1$H)

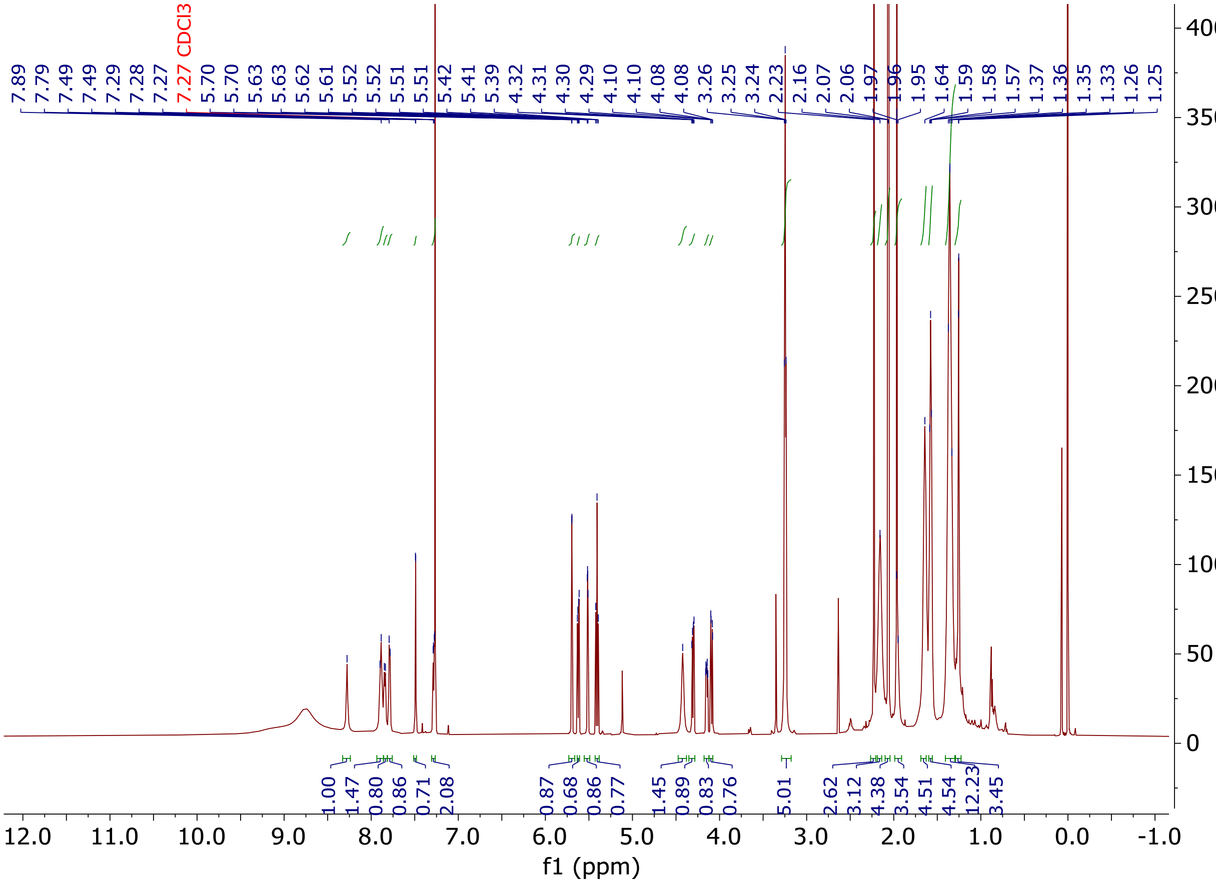

7.89
7.79
7.49
7.49
7.29
7.28
7.27
7.27 CDCl3
5.70
5.70
5.63
5.63
5.62
5.61
5.52
5.52
5.51
5.51
5.42
5.41
5.39
4.32
4.31
4.30
4.29
4.10
4.10
4.08
4.08
3.26
3.25
3.24
2.23
2.16
2.07
2.06
1.97
1.96
1.95
1.64
1.59
1.58
1.57
1.37
1.36
1.35
1.33
1.26
1.25
1.00
1.47
0.80
0.86
0.71
2.08
0.87
0.68
0.86
0.77
1.45
0.89
0.83
0.76
5.01
2.62
3.12
4.38
3.54
4.51
4.54
12.23
3.45
f1 (ppm)

STR-V-177 ($^{13}C$)

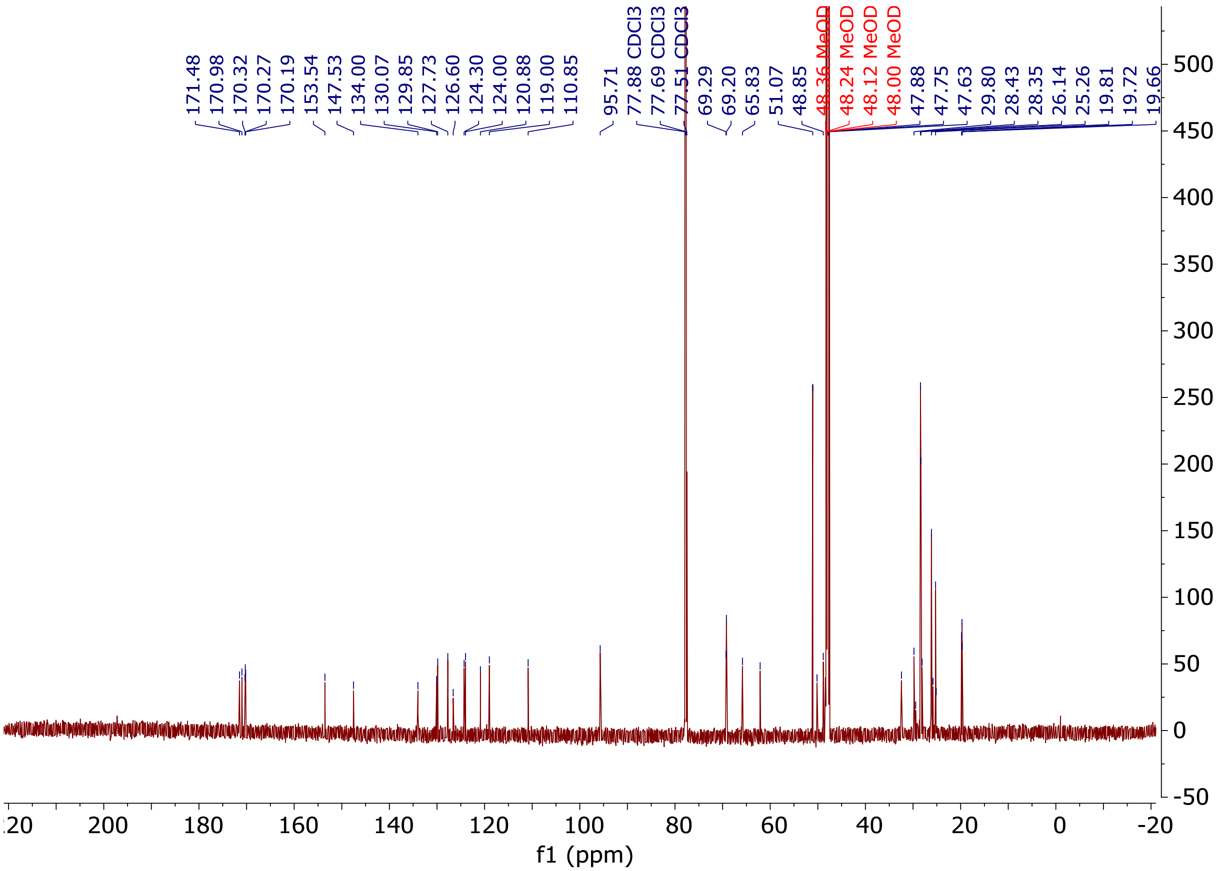
171.48
170.98
170.32
170.27
170.19
153.54
147.53
134.00
130.07
129.85
127.73
126.60
124.30
124.00
120.88
119.00
110.85
95.71
77.88 CDCl3
77.69 CDCl3
77.51 CDCl3
69.29
69.20
65.83
51.07
48.85
48.36 MeOD
48.24 MeOD
48.12 MeOD
48.00 MeOD
47.88
47.75
47.63
29.80
28.43
28.35
26.14
25.26
19.81
19.72
19.66
f1 (ppm)

STR-II-36 ($^{1}H$)

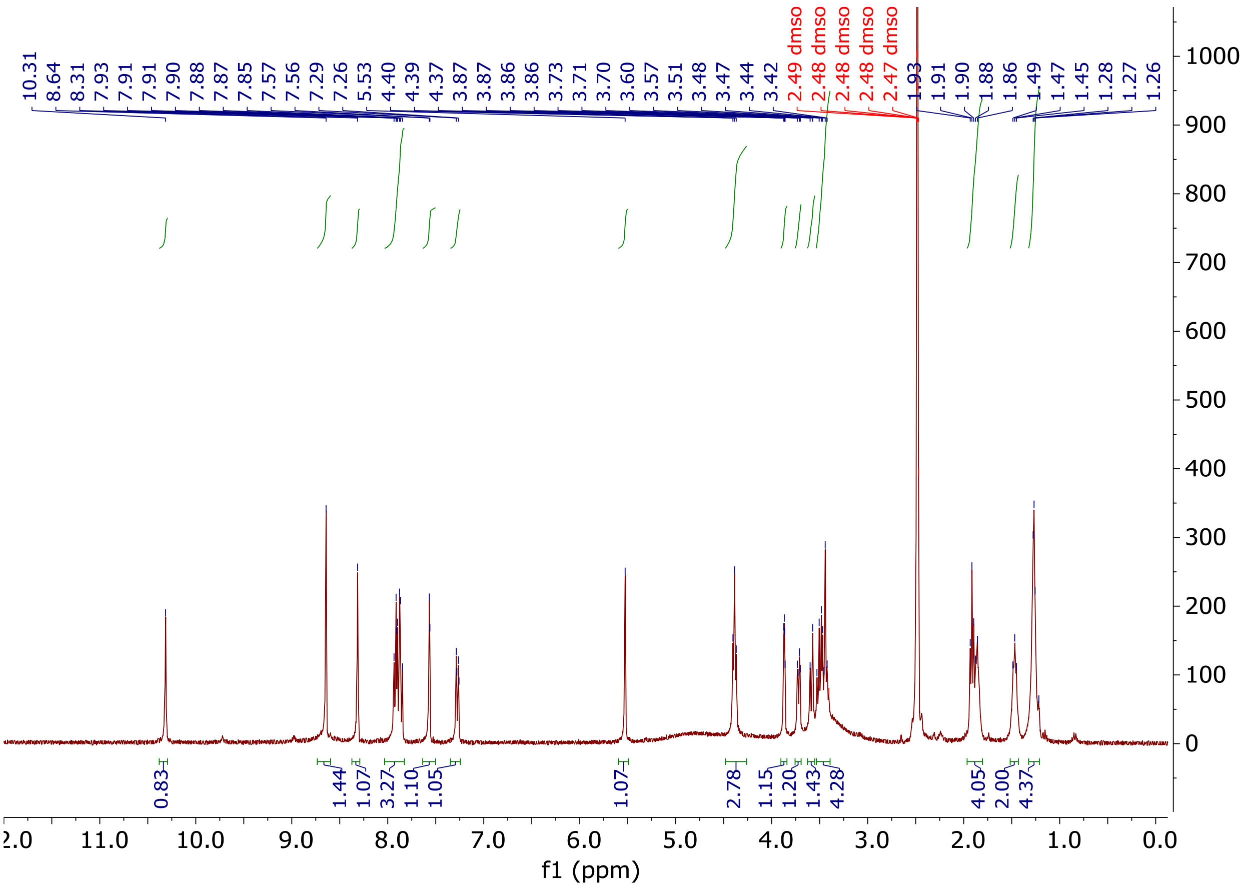

10.31
8.64
8.31
7.93
7.91
7.91
7.90
7.88
7.87
7.85
7.57
7.56
7.29
7.26
5.53
4.40
4.39
4.37
3.87
3.87
3.86
3.86
3.73
3.71
3.70
3.60
3.57
3.51
3.48
3.47
3.44
3.42
2.49 dmso
2.48 dmso
2.48 dmso
2.48 dmso
2.47 dmso
1.93
1.91
1.90
1.88
1.86
1.49
1.47
1.45
1.28
1.27
1.26
0.83
1.44
1.07
3.27
1.10
1.05
1.07
2.78
1.15
1.20
1.43
4.28
4.05
2.00
4.37
f1 (ppm)

STR-II-36 ($^{13}C$)

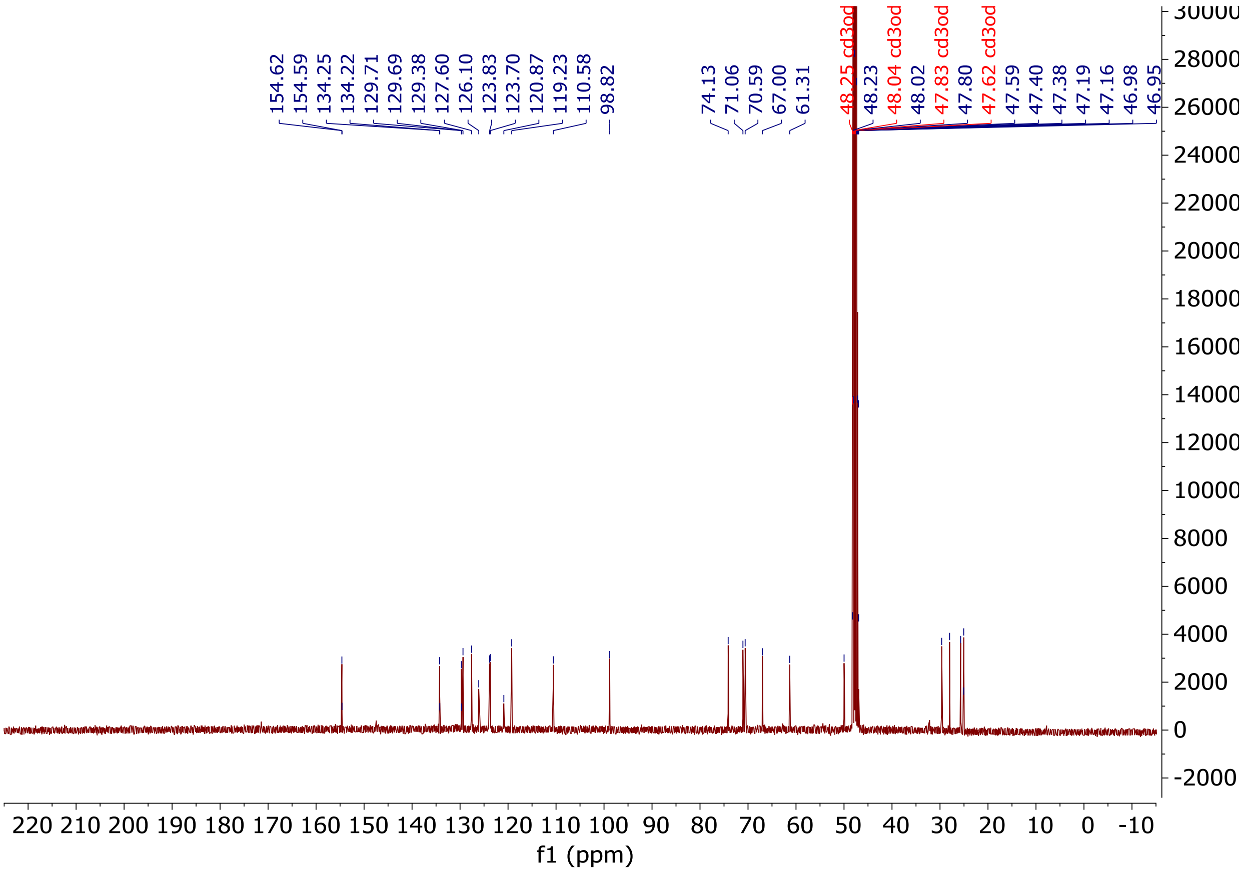
154.62
154.59
134.25
134.22
129.71
129.69
129.38
127.60
126.10
123.83
123.70
120.87
119.23
110.58
98.82
74.13
71.06
70.59
67.00
61.31
48.25 cd3od
48.23
48.04 cd3od
48.02
47.83 cd3od
47.80
47.62 cd3od
47.59
47.40
47.38
47.19
47.16
46.98
46.95
f1 (ppm)

STR-V-105 ($^1H$)

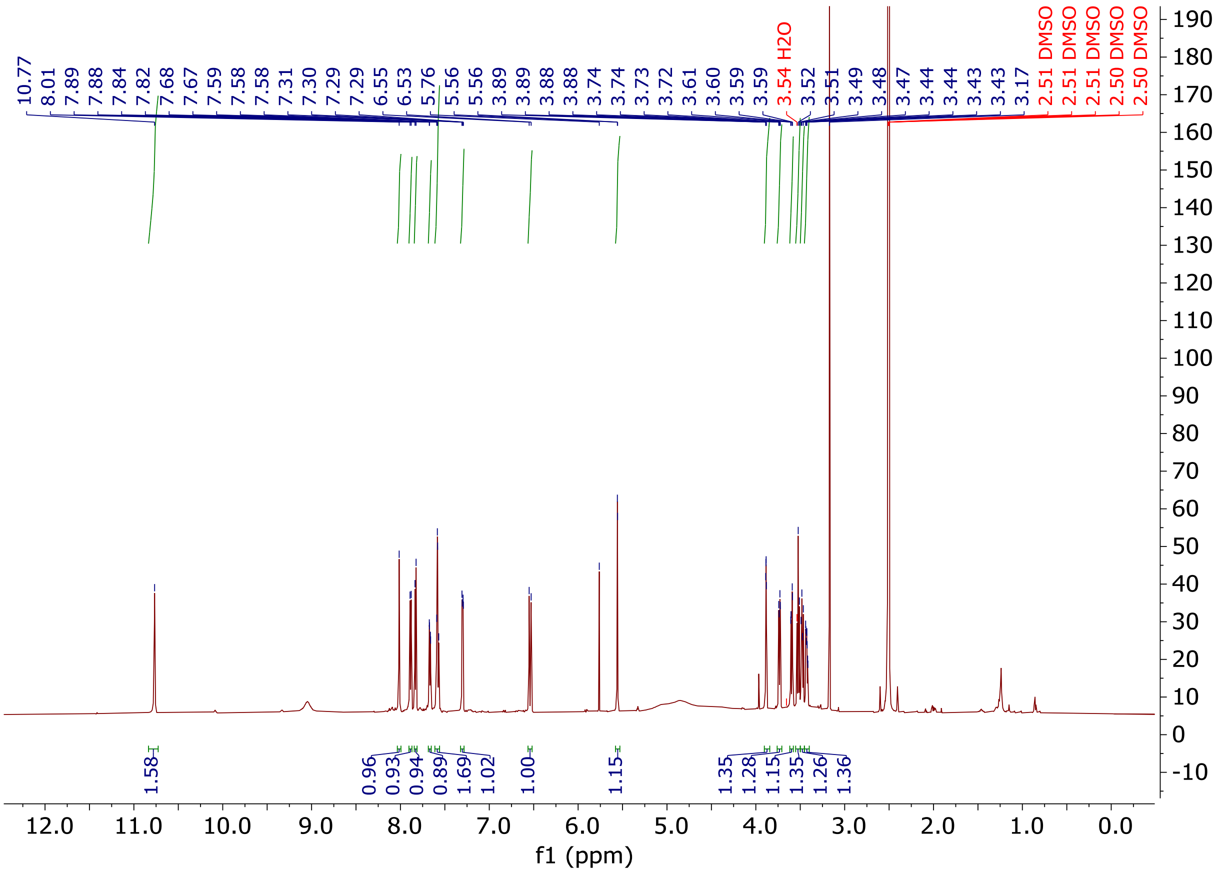
10.77
8.01
7.89
7.88
7.84
7.82
7.68
7.67
7.59
7.58
7.58
7.31
7.30
7.29
7.29
6.55
6.53
5.76
5.56
5.56
3.89
3.89
3.88
3.88
3.74
3.74
3.73
3.72
3.61
3.60
3.59
3.59
3.54 H2O
3.52
3.51
3.49
3.48
3.47
3.44
3.44
3.43
3.43
3.17
2.51 DMSO
2.51 DMSO
2.51 DMSO
2.50 DMSO
2.50 DMSO
1.58
0.96
0.93
0.94
0.89
1.69
1.02
1.00
1.15
1.35
1.28
1.15
1.35
1.26
1.36
f1 (ppm)

STR-V-105 ($^{13}C$)

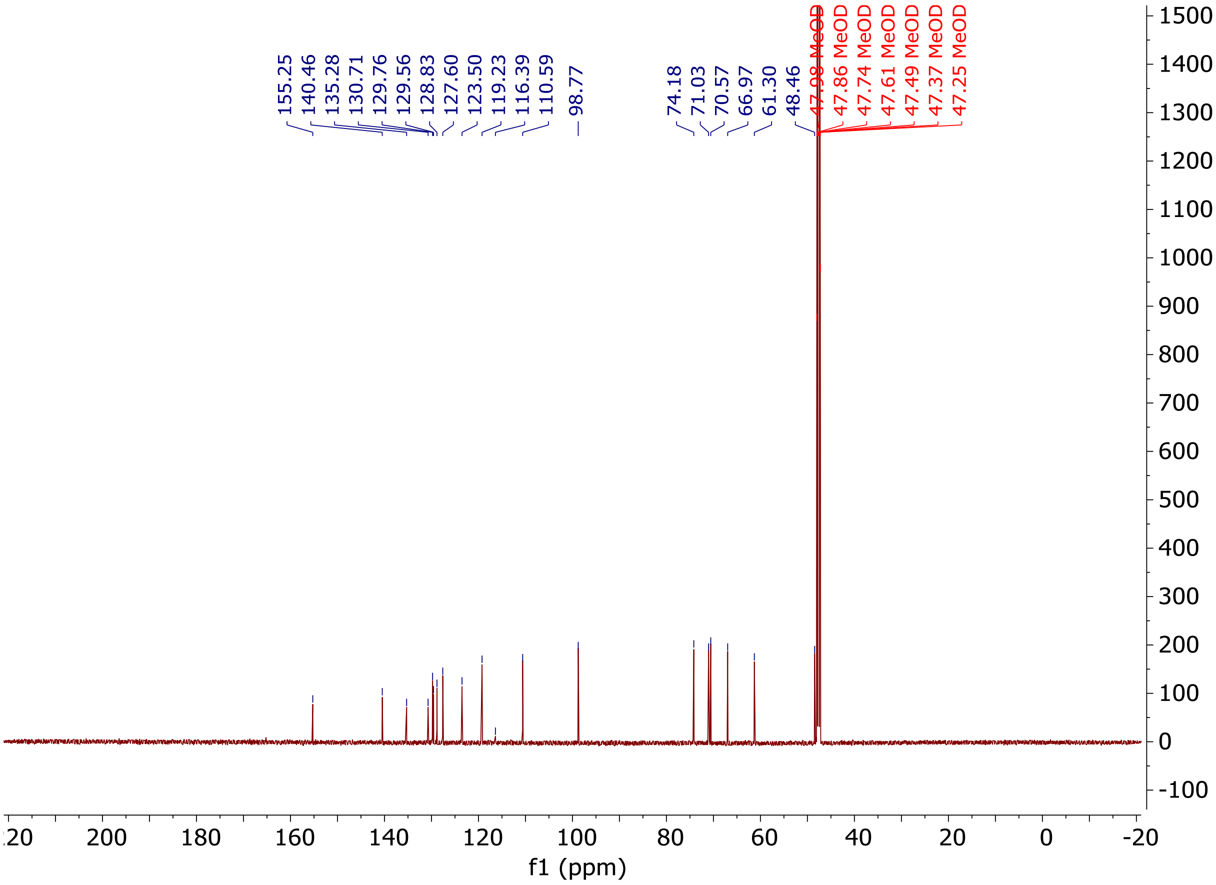
155.25
140.46
135.28
130.71
129.76
129.56
128.83
127.60
123.50
119.23
116.39
110.59
98.77
74.18
71.03
70.57
66.97
61.30
48.46
47.98 MeOD
47.86 MeOD
47.74 MeOD
47.61 MeOD
47.49 MeOD
47.37 MeOD
47.25 MeOD
f1 (ppm)

STR-V-160 ($^{1}H$)

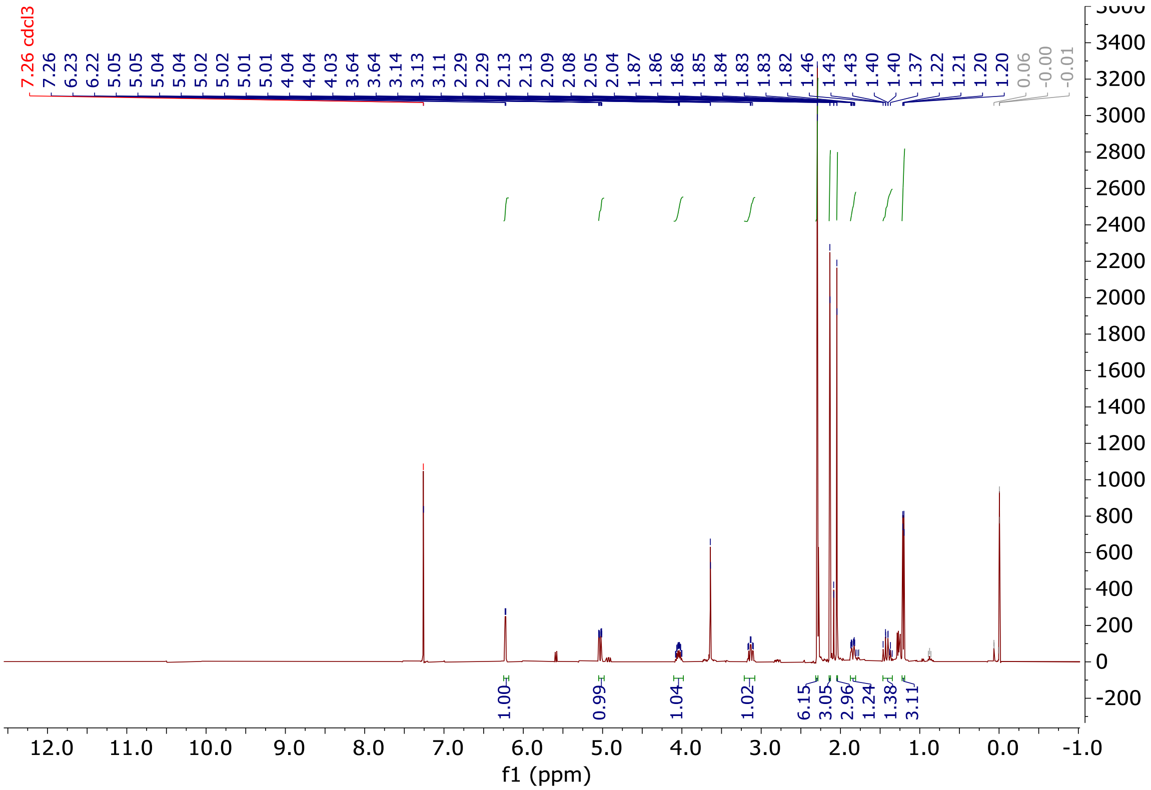

STR-V-161 ($^1$H)

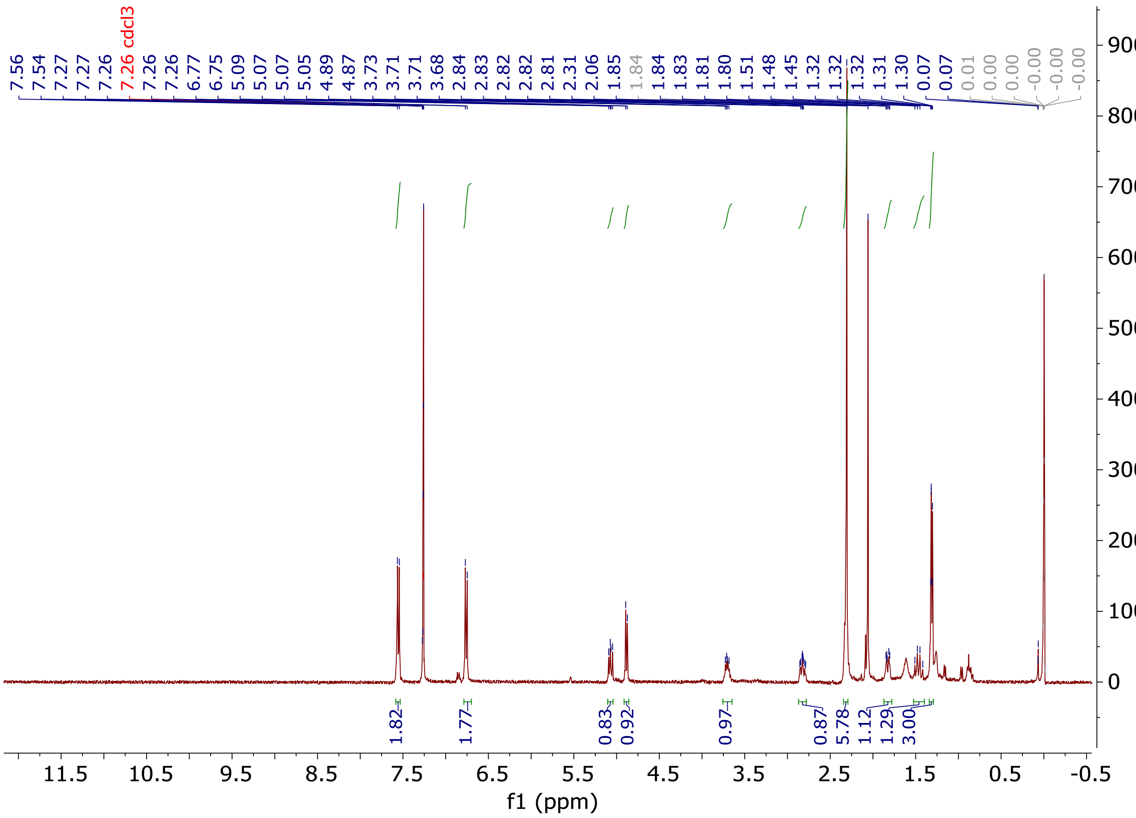

STR-V-163 ($^{1}H$)

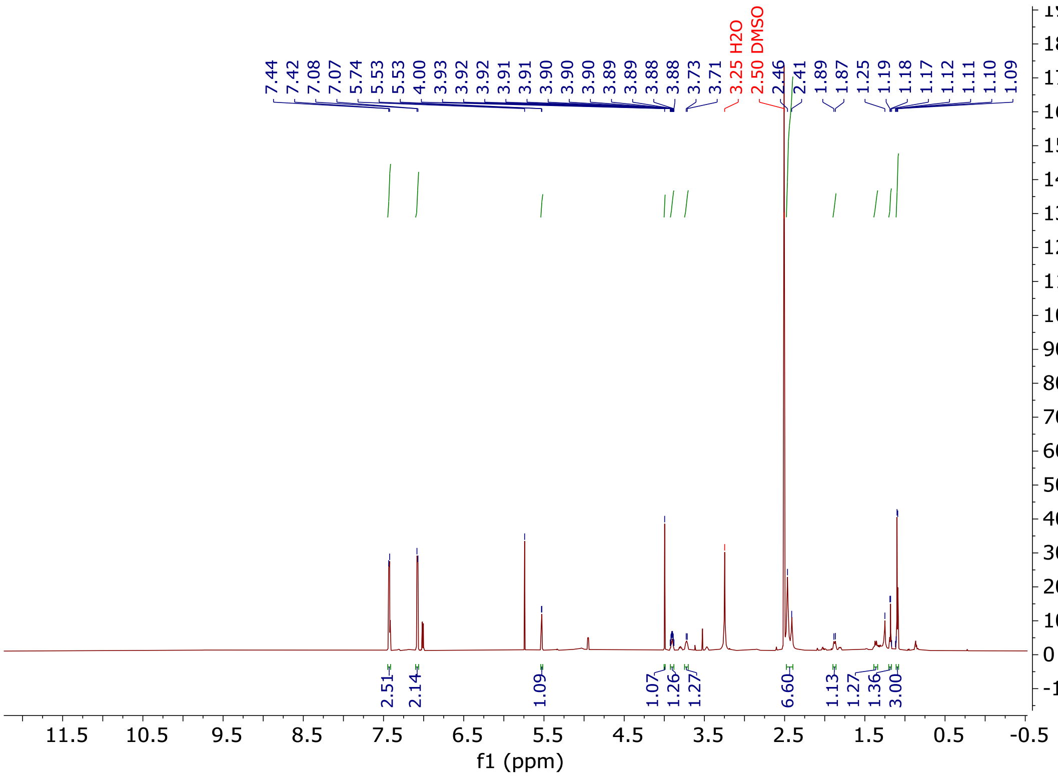
7.44 7.42 7.08 7.07 5.74 5.53 5.53 4.00 3.93 3.92 3.92 3.91 3.91 3.90 3.90 3.90 3.89 3.89 3.88 3.88 3.73 3.71
3.25 H2O
2.50 DMSO
2.46 2.41 1.89 1.87 1.25 1.19 1.18 1.17 1.12 1.11 1.10 1.09
2.51 2.14 1.09 1.07 1.26 1.27 6.60 1.13 1.27 1.36 3.00
11.5 10.5 9.5 8.5 7.5 6.5 5.5 4.5 3.5 2.5 1.5 0.5 -0.5
f1 (ppm)
1900 1800 1700 1600 1500 1400 1300 1200 1100 1000 900 800 700 600 500 400 300 200 100 0 -100

STR-V-165 ($^1$H)

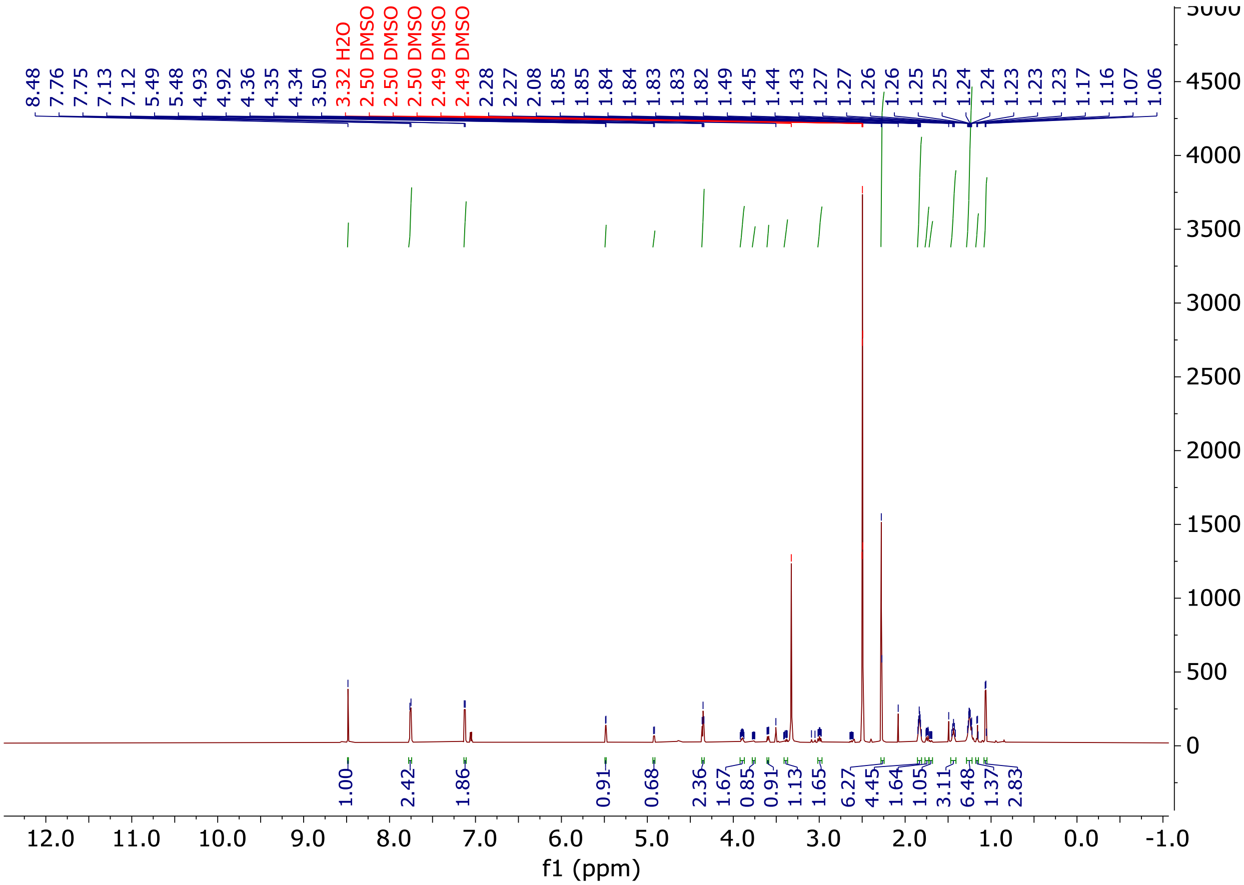
8.48
7.76
7.75
7.13
7.12
5.49
5.48
4.93
4.92
4.36
4.35
4.34
3.50
3.32 H2O
2.50 DMSO
2.50 DMSO
2.50 DMSO
2.49 DMSO
2.49 DMSO
2.28
2.27
2.08
1.85
1.85
1.84
1.84
1.83
1.83
1.82
1.49
1.45
1.44
1.43
1.27
1.27
1.26
1.26
1.25
1.25
1.24
1.24
1.23
1.23
1.23
1.17
1.16
1.07
1.06
1.00
2.42
1.86
0.91
0.68
2.36
1.67
0.85
0.91
1.13
1.65
6.27
4.45
1.64
1.05
3.11
6.48
1.37
2.83
f1 (ppm)

STR-V-165 ($^{13}C$)

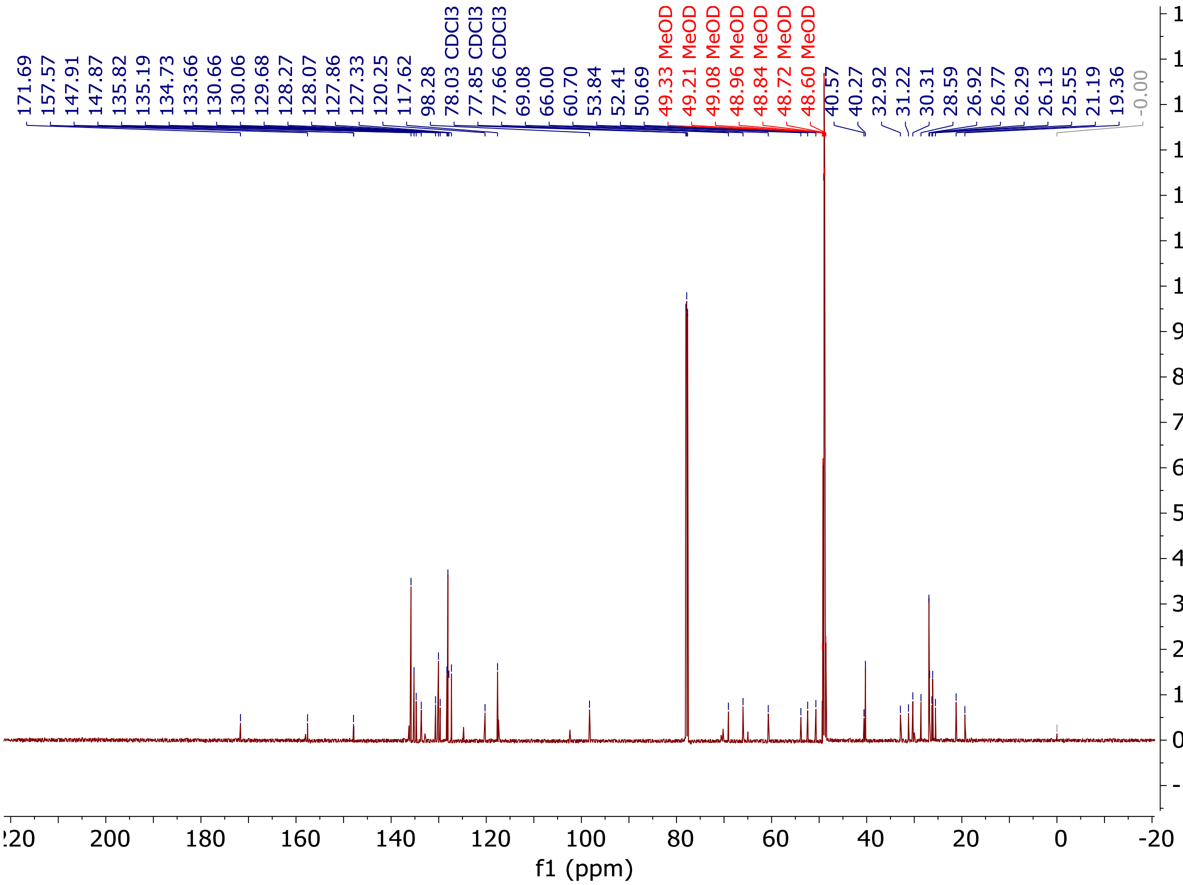
171.69
157.57
147.91
147.87
135.82
135.19
134.73
133.66
130.66
130.06
129.68
128.27
128.07
127.86
127.33
120.25
117.62
98.28
78.03 CDCl3
77.85 CDCl3
77.66 CDCl3
69.08
66.00
60.70
53.84
52.41
50.69
49.33 MeOD
49.21 MeOD
49.08 MeOD
48.96 MeOD
48.84 MeOD
48.72 MeOD
48.60 MeOD
40.57
40.27
32.92
31.22
30.31
28.59
26.92
26.77
26.29
26.13
25.55
21.19
19.36
-0.00
f1 (ppm)

Full gels:

AC-H4 and AC-tubulin

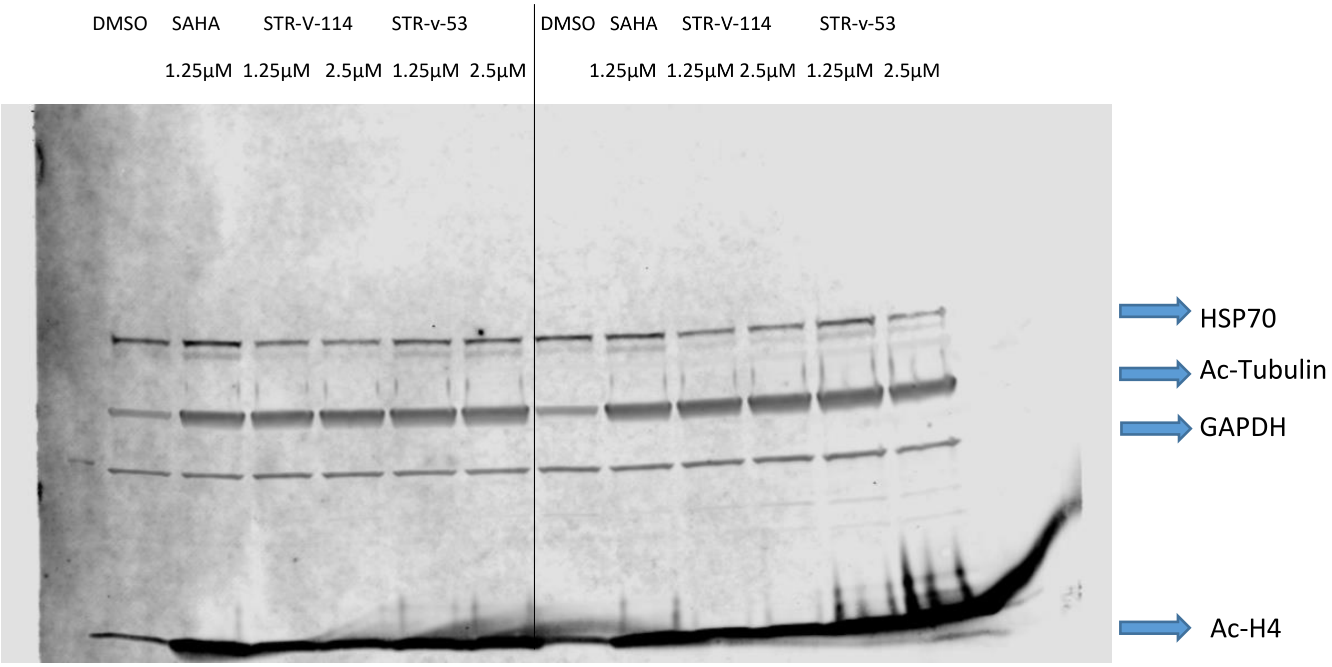


Hep-G2 Caspase 3 cleavage Gel

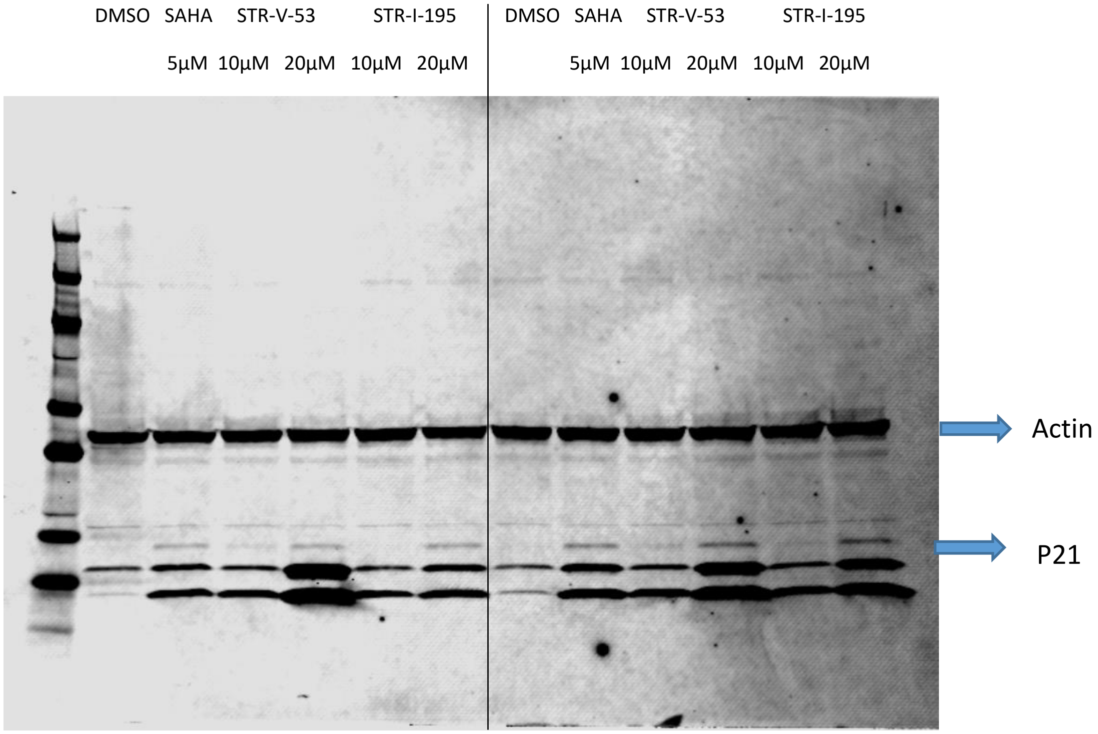

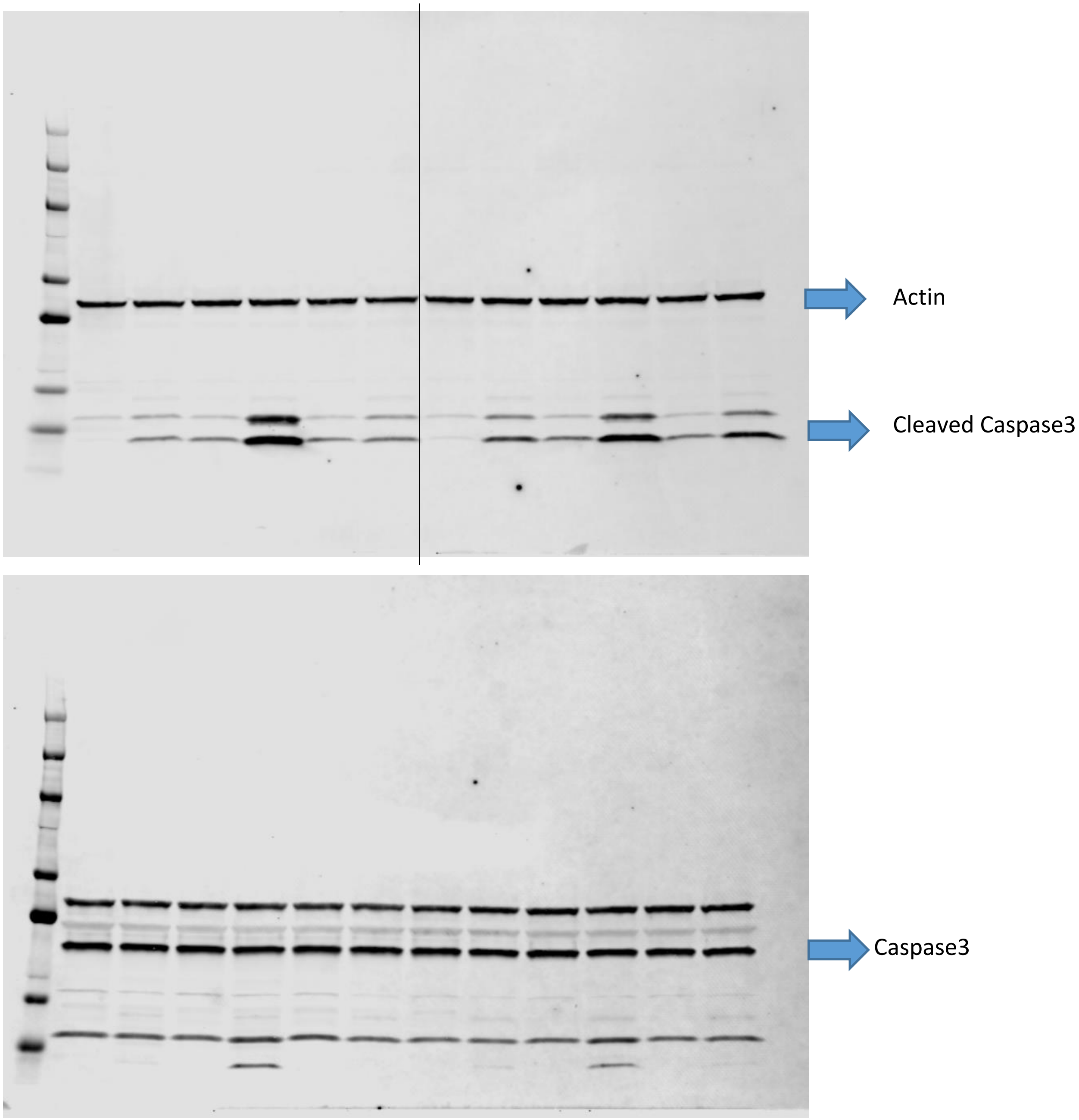

Actin
Cleaved Caspase3
Caspase3

VERO Caspase 3 cleavage Gel

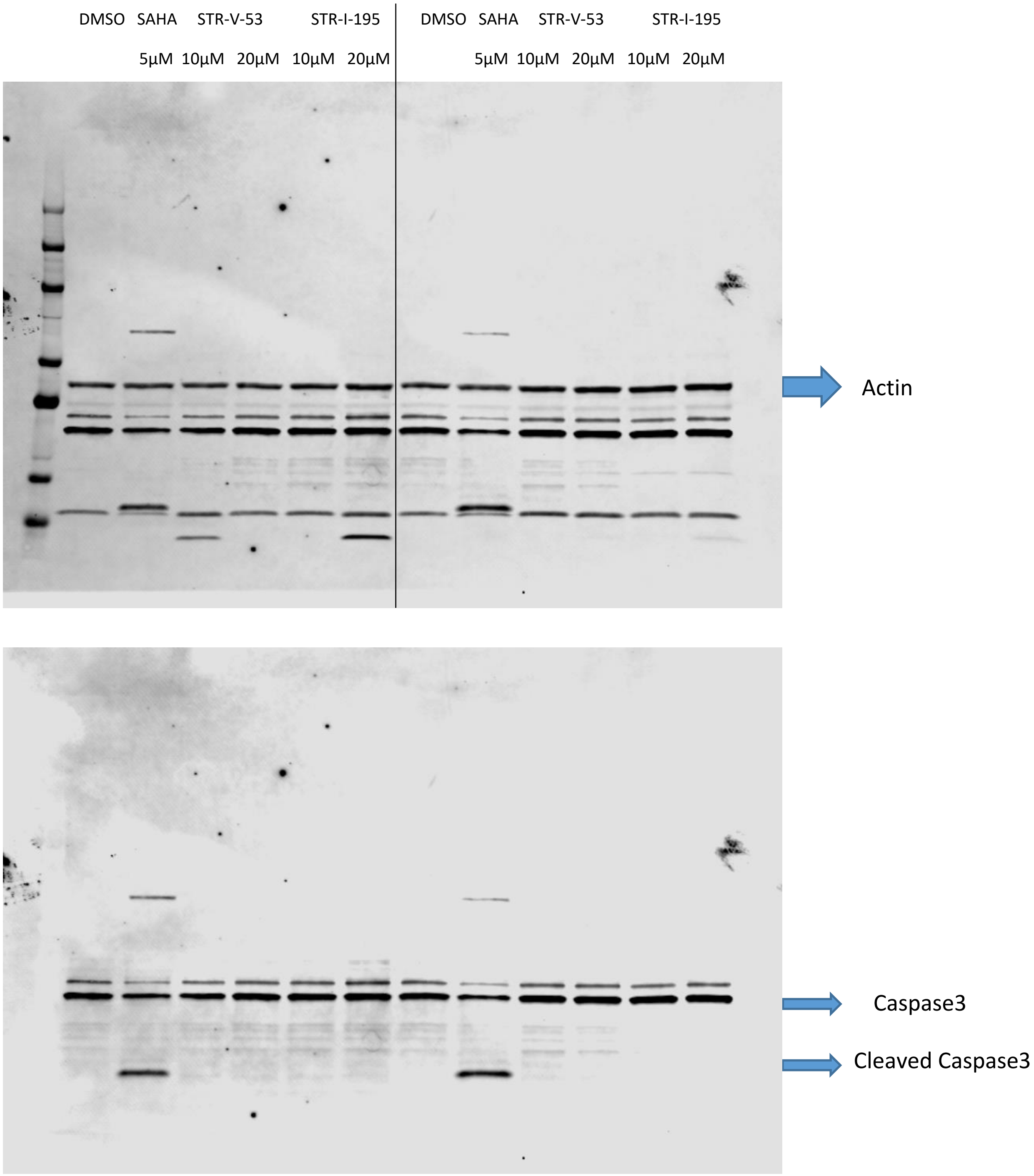

PD1/PD-L1 Gel

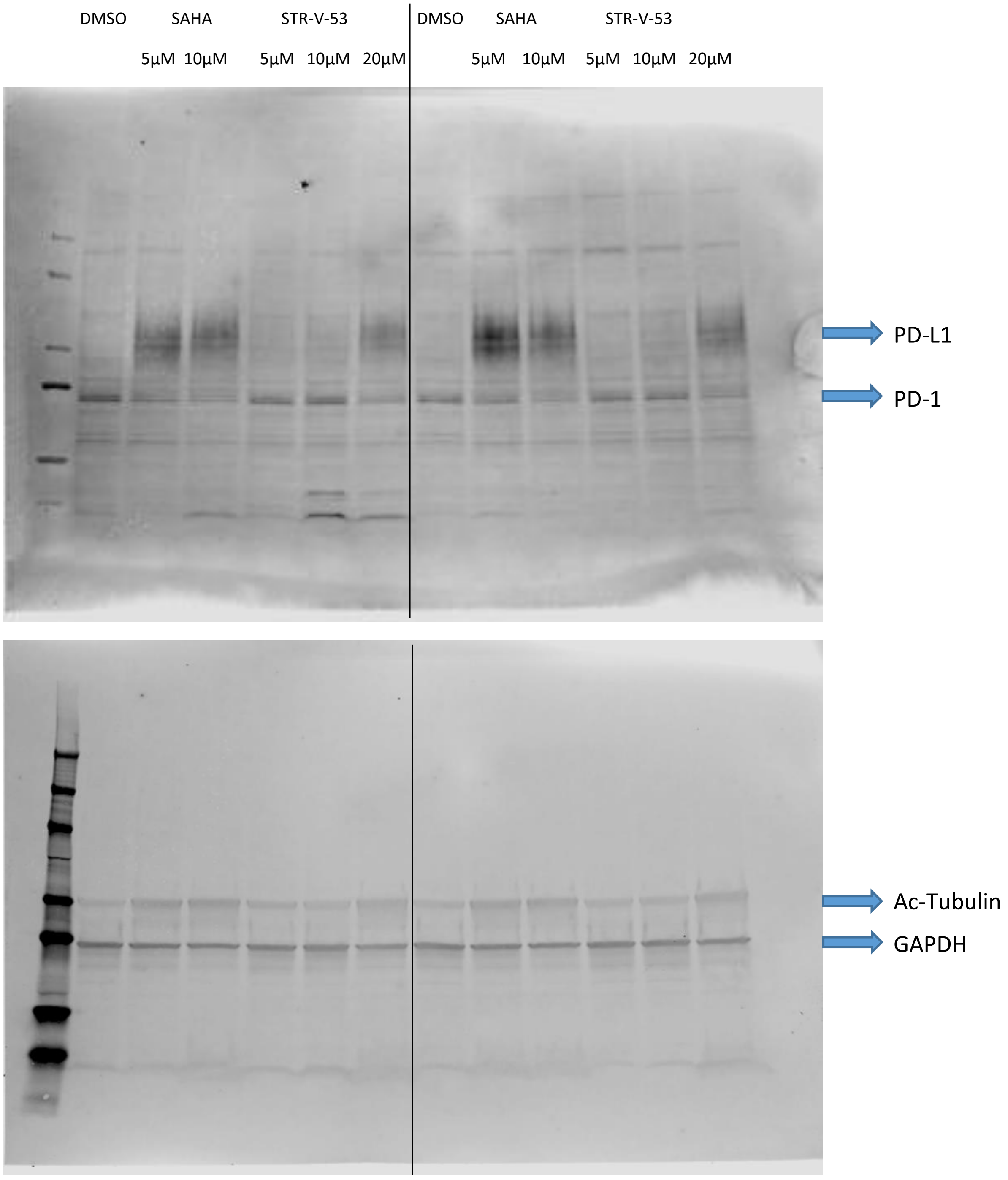
DMSO
SAHA
STR-V-53
5µM
10µM
5µM
10µM
20µM
DMSO
SAHA
STR-V-53
5µM
10µM
5µM
10µM
20µM
PD-L1
PD-1
Ac-Tubulin
GAPDH

CD54

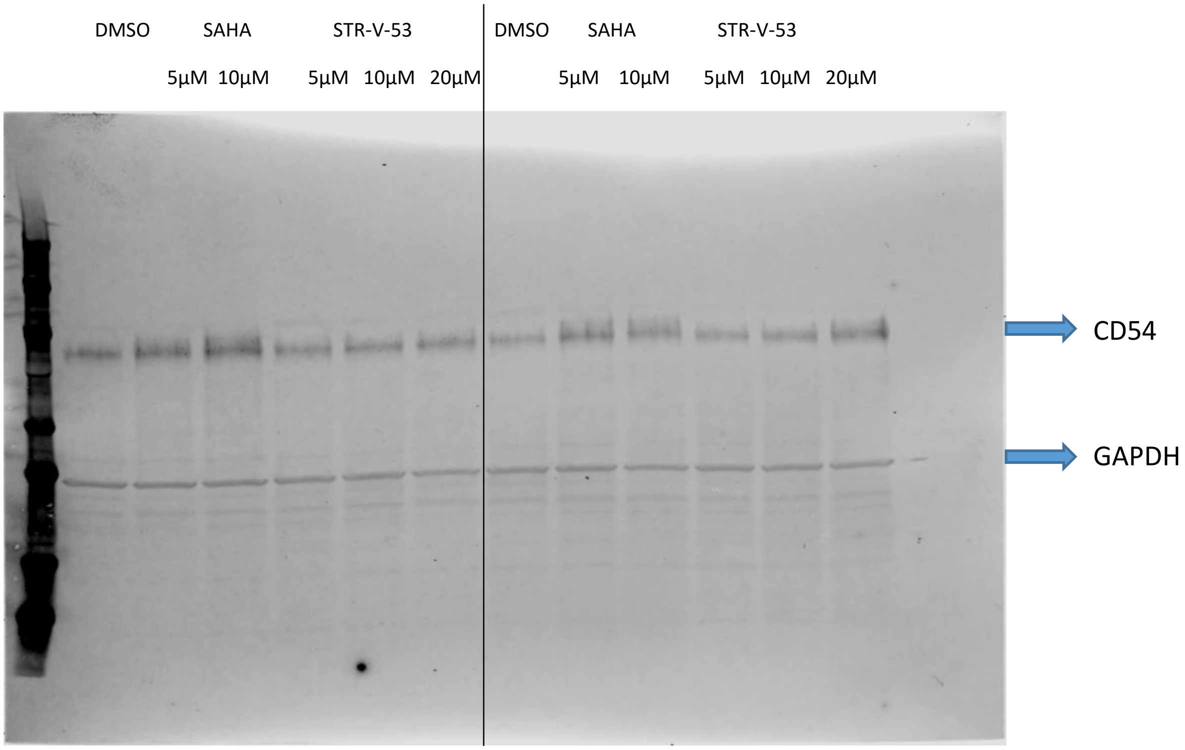


#### Supporting Information References

1. Asante, Y.; Benischke, K.; Osman, I.; Ngo, Q. A.; Wurth, J.; Laubscher, D.; Kim, H.; Udhayakumar, B.; Khan, M. I. H.; Chin, D. H.; Porch, J.; Chakraborty, M.; Sallari, R.; Delattre, O.; Zaidi, S.; Morice, S.; Surdez, D.; Danielli, S. G.; Schäfer, B. W.; Gryder, B. E.; Wachtel, M. PAX3-FOXO1 uses its activation domain to recruit CBP/P300 and shape RNA Pol2 cluster distribution. *Nat. Commun.* **2023**, 14 (1), 8361.
2. Liberzon, A.; Birger, C.; Thorvaldsdóttir, H.; Ghandi, M.; Mesirov, J. P.; Tamayo, P. The Molecular Signatures Database (MSigDB) hallmark gene set collection. *Cell Syst* **2015**, 1 (6), 417-425.
3. Liberzon, A.; Subramanian, A.; Pinchback, R.; Thorvaldsdóttir, H.; Tamayo, P.; Mesirov, J. P. Molecular signatures database (MSigDB) 3.0. *Bioinformatics* **2011**, 27 (12), 1739-1740.
4. Subramanian, A.; Tamayo, P.; Mootha, V. K.; Mukherjee, S.; Ebert, B. L.; Gillette, M. A.; Paulovich, A.; Pomeroy, S. L.; Golub, T. R.; Lander, E. S.; Mesirov, J. P. Gene set enrichment analysis: A knowledge-based approach for interpreting genome-wide expression profiles. *Proc. Natl. Acad. Sci. U.S.A.* **2005**, 102 (43), 15545-15550.
5. Love, M. I.; Huber, W.; Anders, S. Moderated estimation of fold change and dispersion for RNA-seq data with DESeq2. *Genome Biol.* **2014**, 15 (12), 550.
6. Wickham, H., ggplot2: Elegant Graphics for Data Analysis. Springer-Verlag New York: 2016.
